# A parabrachial-hypothalamic parallel circuit governs cold defense in mice

**DOI:** 10.1101/2023.04.19.537288

**Authors:** Wen Z. Yang, Hengchang Xie, Xiaosa Du, Qian Zhou, Yan Xiao, Zhengdong Zhao, Xiaoning Jia, Jianhui Xu, Wen Zhang, Shuang Cai, Zhangjie Li, Xin Fu, Rong Hua, Junhao Cai, Shuang Chang, Jing Sun, Hongbin Sun, Qingqing Xu, Xinyan Ni, Hongqing Tu, Ruimao Zheng, Xiaohong Xu, Hong Wang, Yu Fu, Liming Wang, Xi Li, Haitao Yang, Qiyuan Yao, Tian Yu, Qiwei Shen, Wei L. Shen

## Abstract

Thermal homeostasis is vital for mammals and is controlled by brain neurocircuits. Remarkable advances have been made in understanding how neurocircuits centered in the hypothalamic preoptic area (POA), the brain’s thermoregulation center, control warm defense, whereas mechanisms by which the POA regulates cold defense remain unclear. Here, we confirmed that the pathway from the lateral parabrachial nucleus (LPB) to the POA, is critical for cold defense. Parallel to this pathway, we uncovered that a pathway from the LPB to the dorsomedial hypothalamus (DMH), namely the LPB→DMH pathway, is also essential for cold defense. Projection-specific blockings revealed that both pathways provide an equivalent and cumulative contribution to cold defense, forming a parallel circuit. Specifically, activation of the LPB→DMH pathway induced strong cold-defense responses, including increases in thermogenesis of brown adipose tissue (BAT), muscle shivering, heart rate, and physical activity. Further, we identified a subpopulation of somatostatin^+^ neurons in the LPB that target the DMH to promote BAT thermogenesis. Therefore, we reveal a parabrachial-hypothalamic parallel circuit in governing cold defense in mice. This not only enables resilience to hypothermia but also provides a scalable and robust network in heat production, reshaping our understanding of how neural circuits regulate essential homeostatic behaviors.

## Introduction

Hypothermia caused by cold or famine is one of the major death causes in wild animals and has been a threat to human life throughout history^1–3^. Hypothermia reduces motor function and suppresses immune responses, thereby impairing behavioral reactions and increasing infection risks, respectively^4, 5^. Thus, it is vital to maintain a stable core body temperature (T_core_) or to rapidly recover from a hypothermic state. In response to cold exposure, the brain initiates a series of countermeasures to defend T_core_ from hypothermia, including increases in the thermogenesis of brown adipose tissue (BAT), skeletal muscle shivering, heart rate (HR), and skin vasoconstriction. Several behavioral adaptations are also induced ^6^. However, the neural circuitry underlying these cold-defense activities is not well understood.

The hypothalamic preoptic area (POA) is the known thermoregulation center in mammals^6–10^, and remarkable advances have been made in dissecting POA circuits that mediate warm defense^11–22^. On contrary, the circuitry related the POA in cold defense remains elusive. Previous studies^23^ in rats have suggested that GABAergic neurons in the median preoptic nucleus (MnPO) are critical for cold defense, which receive cold sensory input from the external lateral LPB (LPBel)^24, 25^ and suppress GABAergic neurons in the medial preoptic area (MPA) to disinhibit cold-defense responses. However, in contrast to rat studies, optogenetic activation of the LPB→POA pathway in mice results in hypothermia alone^25, 26^, and activation of GABAergic neurons in the MnPO or MPA in mice could not increase T_core_ (hyperthermic)^13, 14, 19, 27^. Therefore, whether the LPB→POA pathway could function in cold defense in mice is still not clear. Interestingly, recent studies in mice suggested that selective activation of GABAergic arginine vasopressin (AVP) neurons in the MPA or Bombesin-like receptor 3 (Brs3) neurons (mixed glutamatergic and GABAergic) in the MnPO and ventromedial preoptic nucleus (VMPO) could induce hyperthermia^28, 29^, although this induced hyperthermia is rather mild (∼1°C)^30^. Noticeably, it is still debatable the existence of AVP neurons in the POA^20, 31^. Together, the unresolved cold-defense function of the LPB→POA pathway and the mild hyperthermic responses induced by POA neurons in mice raise a concern that the POA may contribute less to cold defense than previously thought. POA-independent mechanisms may also exist to boost cold defense (at least) in mice.

The dorsomedial hypothalamus (DMH) is considered as a thermogenesis center that functions downstream of the POA^10^. DMH lesions selectively impair thermoregulation under cold but not warm challenges^32^. Studies in rats have suggested that DMH neurons are excited or disinhibited by POA cold-defense neurons to increase thermogenesis^23, 33^. Opposite to what is seen in the POA, activation of most thermoregulatory neural types in the DMH triggers hyperthermia (by up to 1.8°C)^14, 34–37^ in mice. To date, only neurons expressing choline acetyltransferase (ChAT) in the DMH, have been shown to induce mild hypothermia (-1°C)^38^. Since diverse types of DMH neurons could induce stronger hyperthermia than POA neurons, it raises an interesting possibility that the DMH may function in parallel to the POA – rather than solely downstream of the POA – to boost cold defense^28, 33, 39^. In retrospect, tracing studies have shown that the LPB sends neural projections directly to the DMH in mice^35, 40^. Therefore, we hypothesized that the LPB directly transmits cold signals to the DMH to boost cold defense cooperatively the LPB→POA pathway.

To test our hypothesis, we first determined the connectivity, neural activity, and function of the LPB→DMH pathway in cold defense. We found that activating this pathway increased T_core_ and promoted a wide spectrum of cold defense activities while blocking this pathway led to profound cold intolerance. This pathway functioned in parallel with the LPB→POA pathway, in that the two pathways provide an equivalent and cumulative contribution to cold defense. Using projection-specific transcriptomic analysis, we identified a somatostatin^+^ (SST^+^) neural subpopulation in the LPB that targets the DMH to promote BAT thermogenesis. Thus, our findings reveal that parabrachial-hypothalamic parallel pathways function cooperatively to promote cold defense in mice, thereby providing a parallel circuit model for understanding thermoregulation.

## Results

### The LPB→POA/DMH pathways are sensitive to cold temperature

To test our hypothesis that the LPB→DMH pathway functions independently of the POA in cold defense, we mapped axonal projection patterns of LPB glutamatergic neurons, the predominant neuronal subtype in the LPB. This was accomplished by injecting AAVs carrying Cre-dependent ChR2-eYFP (AAV9-DIO-ChR2-eYFP) into the LPB of Vglut2-IRES-Cre mice (LPB^Vglut2^ neurons) (**Fig. 1a,b and Extended Data** **Fig. 1**). As expected, LPB^Vglut2^ neurons projected to many regions, including the POA (mostly the MnPO and VMPO) and the DMH. To assess potential collateral projections between axons within the POA and the DMH, we simultaneously injected retrograde AAVs carrying Cre-dependent GFP (Retro-DIO-GFPL10) and Cre-dependent mCherry (Retro-DIO-mCherry) into the POA and the DMH of Vglut2-IRES-Cre mice, respectively (**Fig. 1c**). Three weeks post-injection, these mice were exposed to cold (10°C, 2 h) or warm (38°C, 2 h) stimuli and then sacrificed for cFos immunostaining to evaluate the thermal responsiveness of neurons. We verified that GFPL10 expression was limited to the POA (mostly MnPO and VMPO), and that mCherry expression was limited to the DMH (**Extended Data Fig. 2a,b**). POA-projecting (green) and DMH-projecting (red) LPB^Vglut2^ neurons were more concentrated in the external lateral LPB (LPBel) and dorsal LPB (LPBd) where temperature-responsive neurons reside^24, 25^ (**Fig. 1d**). Among all warm-activated LPB neurons (cFos^+^; 100%), 40% projected to the POA, 13% projected to the DMH, and 7% projected to both regions (**Fig. 1e,f and Extended Data Fig. 2c**). These results suggest that warm-activated LPB neurons mainly innervate the POA compared to the DMH.

**Fig. 1.**
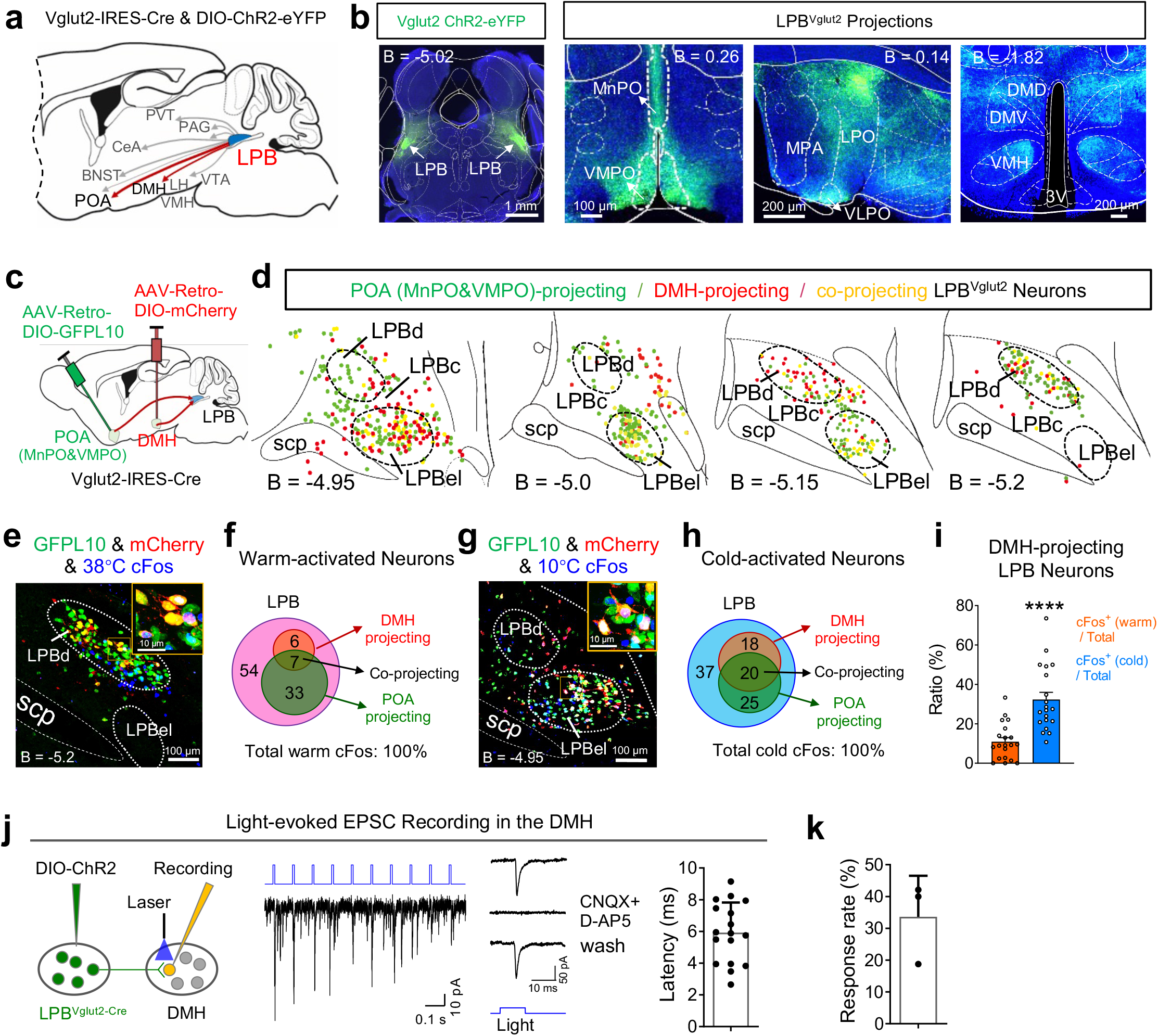
Properties of the LPB→POA/DMH pathways and their sensitivity to temperature. (a-b) Whole-brain projection pattern of LPB^Vglut2^ neurons viewed using AAV-mediated axonal tracing. The injection site and representative axonal images in the POA and the DMH are shown in (b). The scale bar is with the image (as with all brain images). B, bregma; BNST, bed nucleus of the stria terminalis; CeA, central nucleus of the amygdala; 3V, third ventricle; LPB, lateral parabrachial nucleus; PAG, periaqueductal gray matter; VTA, ventral tegmental area; LPO, lateral preoptic area; LH, lateral hypothalamus; PVT, paraventricular thalamic nucleus; MPA, medial preoptic area; MnPO, median preoptic area; VLPO, ventrolateral preoptic area; VMPO, ventromedial preoptic nucleus; VMH, ventromedial hypothalamic nucleus; POA, preoptic area; DMH, dorsomedial hypothalamic nucleus; DMV, dorsomedial hypothalamic nucleus, ventral part; DMD, dorsomedial hypothalamic nucleus, dorsal part. (c) Mapping POA-projecting and DMH-projecting LPB^Vglut2^ neurons. We simultaneously injected two-colored retrograde tracing AAVs, namely AAV-Retro-DIO-GFPL10 and AAV-Retro-DIO-mCherry, into the POA (MnPO and VMPO) and the DMH of Vglut2-IRES-Cre mice, respectively. (d) Distribution of POA-projecting LPB^Vglut2^ neurons (green, GFP^+^), DMH-projecting LPB^Vglut2^ neurons (red, mCherry^+^), and co-projecting LPB^Vglut2^ neurons that send projections to both the POA and DMH (yellow, GFP^+^ & mCherry^+^). Images are from a representative mouse. Each dot represents one neuron. LPBc, lateral parabrachial nucleus, central part; LPBd, lateral parabrachial nucleus, dorsal part; LPBel, lateral parabrachial nucleus, external and internal part; scp, superior cerebellar peduncle. (e-f) Overlap between DMH/POA-projecting LPB^Vglut2^ neurons and warmth-activated cFos. Mice were exposed to 38° C for 2 h before sacrificing for cFos staining. A representative image is shown in (e). Overlap ratios are indicated in (f), where the total number of cFos^+^ neurons is 100%. (g-h) Overlap between DMH/POA-projecting LPB^Vglut2^ neurons and cold-activated cFos. Mice were exposed to 10° C for 2 h before sacrificing for cFos staining. A representative image is shown in (g). Overlap ratios are indicated in (h), where the total number of cFos^+^ neurons is 100%. (i) The ratio of warm- and cold-activated DMH-projecting LPB^Vglut2^ neurons, where the total number of DMH-projecting LPB^Vglut2^ neurons is 100% (20 slices each from 3 mice). (j) Excitatory postsynaptic currents (EPSCs) were recorded randomly from DMH neurons after photoactivation of LPB^Vglut2 & ChR2^ terminals in the DMH (blue, 7 mW, 10 ms). EPSCs were blocked by GluR antagonists D-AP5 and CNQX. The mean latency of responsive neurons is shown on the right (n = 17 neurons). (k) The EPSC response rate of DMH neurons to light stimulation (n = 3 mice). A total of 17 responsive neurons out of 51 randomly recorded DMH neurons were from three mice. All data are the mean ± sem, and were analyzed using the unpaired t-test in (i). P-values were calculated based on statistical tests in Extended Data Table 2. ****p ≤ 0.0001.

Among cold-activated LPB neurons (cFos^+^; 100%), 45% projected to the POA, 38% projected to the DMH, and 20% projected to both regions (**Fig. 1g,h and Extended Data Fig. 2d**). These results suggest that a similar ratio of cold-activated LPB neurons innervates the POA or DMH, and that there are substantial collateral projections between these projection axons. We further analyzed the temperature responsiveness of DMH-projecting LPB neurons. About 30% responded to cold, whereas only 10% responded to warmth (**Fig. 1i**), suggesting a biased response to cold temperature.

To verify whether LPB^Vglut2^ neurons directly innervate DMH neurons, we injected AAVs carrying Cre-dependent ChR2 into the LPB of Vglut2-Cre mice and recorded light-induced excitatory postsynaptic currents (EPSCs) from DMH neurons while photostimulating ChR2-expressing neural terminals in the DMH projected from LPB^Vglut2^ neurons (**Fig. 1j**). Light stimulations faithfully induced EPSCs and could be blocked by the glutamate receptor antagonists CNQX and AP5 (**Fig. 1j**, middle), indicating a glutamatergic transmission. Latency after photoactivation was within the range of a monosynaptic connection (**Fig. 1j**, right)^41^. Notably, nearly 33% of recorded DMH neurons (17/51 neurons from three mice; randomly selected) displayed EPSCs to light stimulations (**Fig. 1k**), suggesting that the DMH receives dense inputs from the LPB.

### Neural activity dynamics of the LPB→ DMH pathway in response to temperature

To visualize neural activity dynamics of the LPB^Vglut2^ → DMH pathway in response to temperature stimuli, we used the calcium reporter GCaMP6s and fiber photometry to record calcium dynamics^42^. We first recorded calcium dynamics at LPB^Vglut2^ ^&^ ^GCaMP6s^ terminals in the DMH, where these terminals were in close proximity to cold-induced cFos (**Fig. 2a,b**). Floor temperature (T_floor_) was controlled by a Peltier device (**Fig. 2a**). As expected, these terminals displayed larger responses to floor cooling (25→10°C) compared to floor warming (25→38°C) (**Fig. 2c-e**).

**Fig. 2.**
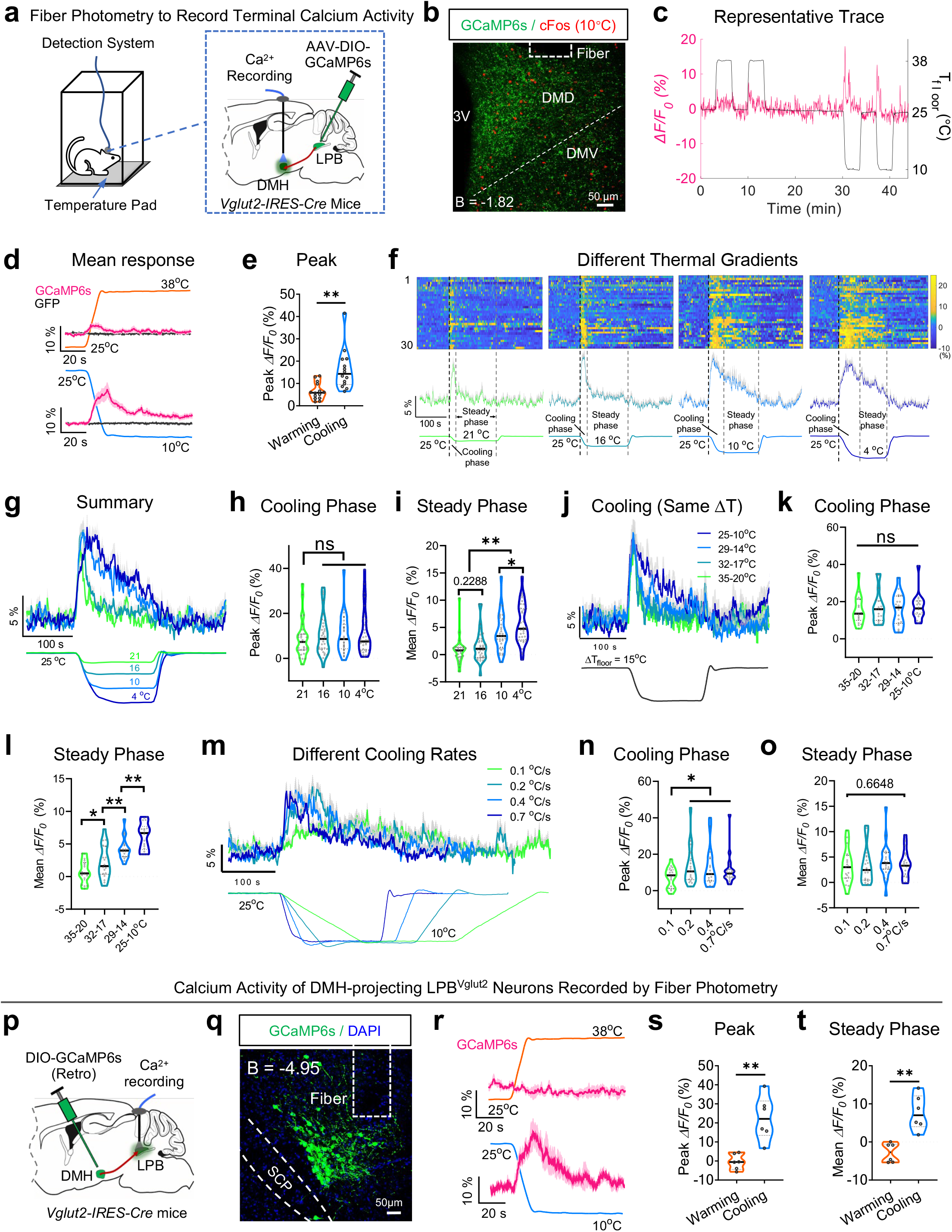
Neural dynamics of LPB→DMH projection in response to thermal stimuli. (a) Fiber photometry to record the calcium dynamics of LPB^Vglut2^ neural terminals in the DMH with GCaMP6s. The calcium indicator GCaMP6s (AAV9-hSyn-DIO-GCaMP6s) was injected into the LPB of Vglut2-IRES-Cre mice, and a recording fiber was implanted above the DMH. Floor temperature (T_floor_) was controlled by a Peltier pad. (b) Representative expression of GCaMP6s from LPB^Vglut2^ neural terminals in the DMH and cold-induced cFos immunoactivity. Mice were exposed to 10° C for 2 h to induce cFos expression. 3V, third ventricle; DMD, dorsomedial hypothalamus, dorsal part; DMV, dorsomedial hypothalamus, ventral part. (c) A representative trace showing calcium dynamics of LPB^Vglut2^ neural terminals in the DMH in response to warm (25→38°C) and cold (25→10°C) stimuli. T_floor_ is plotted in black. ΔF/F_0_ represents the change in GCaMP6s fluorescence from the mean level [t = (−120 to 0 s)] (as to d, f, g, j, m, and r). (d-e) Calcium dynamics after warming (25→38°C) or cooling (25→10° C) the floor. GFP was used as a control (GCaMP6s, n = 14 mice; GFP, n = 5 mice). The peak ΔF/F_0_ in (e) represents the maximum ΔF/F_0_ after warming or cooling the floor. (f-g) Calcium dynamics of LPB^Vglut2^ neural terminals in the DMH in response to different cool/cold floor temperatures, namely 25→21°C, 25→16°C, 25→10°C, and 25→4° C changes. The raster plot in (f, upper part) is the calcium activity from 10 mice and three trials per mouse; the average of signals are plotted in (f, lower part). The cooling and steady phases of T_floor_ are defined in (f). The mean responses under four different T_floor_ changes are replotted together in (g). (h-i) Quantification of calcium signals during the cooling phase (h) and the steady phase (i) defined in (f). The peak ΔF/F_0_ in (h) represents the maximum ΔF/F_0_ during the cooling phase (as to k, n, and s). The mean ΔF/F_0_ in (i) represents the averaged ΔF/F_0_ during the steady phase (as to l, o, and t) (n = 10 mice and 3 trials per mouse). (j) Calcium dynamics of LPB^Vglut2^ neural terminals in the DMH in response to cooling from different starting temperatures but the same temperature reduction (ΔT_floor_ = -15° C, starting from 25, 29, 32, and 35° C). Data are from 6 mice and three trials per mouse. (k-l) Quantification of calcium signals during the cooling (k) and steady (l) phases in response to the same amplitude of cooling shown in (j) (n = 6 mice and 3-5 trials per mouse). (m) Calcium dynamics of LPB^Vglut2^ neural terminals in the DMH in response to different cooling rates. T_floor_ was changed from 25 to 10°C at different rates (0.1, 0.2, 0.4, 0.7° C/sec; n = 10 mice and 2 trials per mouse). (n-o) Quantification of calcium signals during the cooling (n) and steady (o) phases in response to indicated cooling rates (n = 10 mice and 2 trials per mouse). (p-q) Recording from DMH-projecting LPB^Vglut2^ neurons (p) and representative expression (q) of GCaMP6s. Retrograde traveling AAV-Retro-hSyn-Flex-GCaMP6s were injected in the DMH of Vglut2-Cre mice, which traveled to the LPB to drive GCaMP6s expression in the soma of LPB^Vglut2^ neurons. (r) Calcium dynamics of DMH-projecting LPB^Vglut2^ neurons in response to floor warming (25→38°C) or cooling (25→10° C) (n = 6 mice). (s) Peak ΔF/F_0_ values during floor warming or cooling (n = 6 mice, average ΔF/F_0_ of 5 trials per mouse). (t) Mean ΔF/F_0_ values during the steady phase after floor warming or cooling (n = 6 mice, average ΔF/F_0_ of 5 trials per mouse). All data are the mean ± sem (except c), and (h, i, k, l, n, o) were analyzed by one-way RM ANOVA followed by Bonferroni’s multiple comparisons test, (e, s, t) were analyzed by unpaired t-test. P-values are calculated based on statistical tests in Extended Data Table 2. *p ≤ 0.05; **p ≤ 0.01; ns, not significant.

To decode the responses to cold temperatures, we measured calcium dynamics during a series of temperature steps with the same starting T_floor_ (25°C) and cooling rate, but ending at different T_floor_, namely 4, 10, 16, or 21°C (**Fig. 2f**). Interestingly, we observed robust calcium responses during the cooling phases (referred as cooling response) for all four temperature steps (**Fig. 2g**). Yet, peak values of these cooling responses are similar (**Fig. 2h**), suggesting that calcium responses do not encode cooling amplitudes. These results are consistent with the responses seen in cold-responsive spinal cord relay neurons^43^. As the T_floor_ stabilized, neuronal responses gradually reduced (or adapted to the steady temperature). This adaptive feature is also consistent with patterns seen in the majority (∼70%) of peripheral thermosensory neurons^44^. Next, we quantified mean responses during the steady phase (referred as cold response) and found there were no differences between the responses at 16 and 21°C (**Fig. 2i**). Yet, as T_floor_ became less than 10°C, cold responses increased significantly with colder T_floor_ (10 or 4°C) (**Fig. 2i**). Thus, the LPB→DMH projection is sensitive to different cool/cold temperatures, where 10°C appears to be the turning point for faster increases in cold responses. This projection is also responsive to cooling but not sensitive to different cooling amplitudes.

We further measured calcium dynamics in response to four T_floor_ cooling steps, with the same cooling rate and drop in temperature (ΔT = -15°C) but different starting temperatures (from 25 to 35°C) (**Fig. 2j**). These cooling steps induced similar peak cooling responses (**Fig. 2j,k**), suggesting an insensitivity to starting/ending temperatures. However, the cold responses increased with colder ending temperatures (**Fig. 2l**). These data suggest that cold responses, rather than cooling responses, are sensitive to cold temperature values.

To characterize responses to different cooling rates, we measured calcium dynamics in response to a series of cooling rates when cooling from 25 to 10°C. Calcium signals increased in similar dynamics as the temperature changes (**Fig. 2m**), but no peak differences were detected between the three higher cooling rates (0.2, 0.4, or 0.7°C/s) (**Fig. 2n**), which is consistent with the responses seen in spinal cord relay neurons^43^. However, peak responses were smaller at a much slower cooling rate (0.1°C/s) (**Fig. 2n**), suggesting a very weak sensitivity to cooling rates. As the ending temperatures were the same, the cold responses were not different (**Fig. 2o**). Hence, these results suggest that cooling responses of LPB^Vglut2^ terminals in the DMH are weakly sensitive to cooling rates.

Finally, we sought to verify the sensitivity of the LPB^Vglut2^ → DMH pathway to cold temperatures by recording calcium levels within the soma of DMH-projecting LPB^Vglut2^ neurons (**Fig. 2p**). We injected retrograde traveling AAVs carrying Cre-dependent GCaMP6s into the DMH to drive the expression of GCaMP6s in the LPB of Vglut2-Cre mice (**Fig. 2q**). As expected, these DMH-projecting LPB^Vglut2^ neurons were selectively activated by cooling (25→10°C) and cold temperatures (10°C) compared to warming (25→38°C) or warm temperatures (38°C) (**Fig. 2r-t**), suggesting a biased sensitivity to cooling/cold temperatures. Together, we show that the LPB→DMH pathway is responsive to both cooling and cold temperatures. The cold responses are sensitive to different cold temperatures. Yet, akin to spinal cord relay neurons, the peak cooling responses are not sensitive to temperature values and cooling amplitudes, and are only weakly sensitive to cooling rates.

### DMH-projecting LPB neurons that do not collaterally project to the POA (DMH^only^-projecting) are sufficient to mediate cold defense

After confirming the biased response of the LPB→DMH pathway to cool/cold temperatures, we sought to evaluate the function of the LPB→DMH/POA pathways in cold defense. As half of the DMH-projecting cold-responsive LPB neurons also sent collateral projections to the POA (**Fig. 1h** **and Extended Data Fig 3**), we, therefore, tested the cold-defense function of LPB neurons that projected only to the DMH (DMH^only^-projecting) and compared their function to LPB neurons that project only to the POA (POA^only^-projecting). To do so, we used the Cre_off & FlpO_on strategy^45^ to express neurotoxin TeNT (**Fig. 3a,b** **and Extended Data Fig. 4a-b**). On day 1, we injected retrograde AAVs carrying Cre (AAV-Retro-hSyn-Cre) into the POA (MnPO and VMPO) and AAVs carrying Cre_off & Flp_on TeNT into the LPB, where Cre would retrogradely travel to the LPB to turn off TeNT expression (Cre_off). Turning off TeNT expression unblocked POA-projecting LPB neurons. On day 8, we injected retrograde AAVs carrying FlpO (AAV-Retro-hSyn-FlpO) into the DMH to turn on the expression of TeNT in the LPB (FlpO_on), thus blocking DMH^only^-projecting LPB neurons (**Fig. 3a** **and Extended Data Fig. 4a**). The GFP carried in AAV9-hSyn-Cre_off-FlpO_on-eGFP was used as the control for TeNT (**Fig. 3a**). Using the same strategy, we also blocked POA^only^-projecting LPB neurons (**Fig. 3b** **and Extended Data Fig. 4b**). To validate the specificity of this strategy, we co-injected AAV9-hSyn-DIO-mCherry with AAV9-Cre_off & FlpO_on-TeNT-eGFP into the LPB, where mCherry labels POA-projecting LPB neurons after recombination by Cre traveling from the POA and TeNT-GFP labels DMH^only^-projecting LPB neurons. The low amount of overlap (< 2%) between mCherry (POA-projecting) and TeNT-eGFP (DMH^only^-projecting) indicated that this strategy was specific (**Extended Data Fig. 4c**). The overlap between DMH-projecting and POA^only^-projecting neurons was also very low (< 2%; **Extended Data Fig. 4d**).

**Fig. 3.**
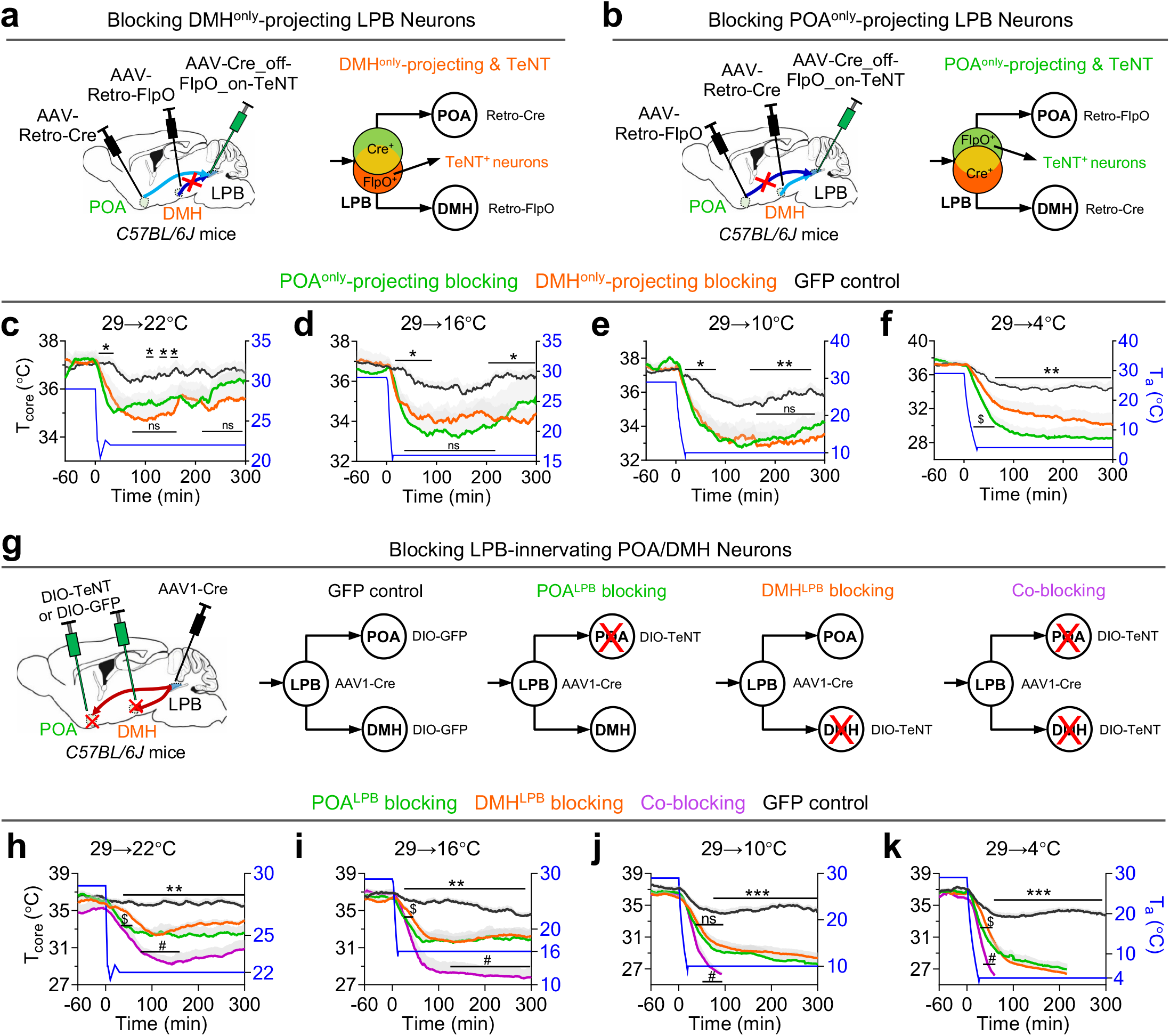
The LPB→POA and LPB→DMH pathways provide an equivalent and cumulative contribution to cold defense. (a) Blocking LPB neurons that project to the DMH but do not send collateral projections to the POA (DMH^only^-projecting LPB neurons). See also Extended Data Fig. 4a for a detailed description. Briefly, AAVs carrying Cre_off-FlpO_on-TeNT were injected in the LPB of C57BL/6J mice. Retrograde traveling AAV-Retro-hSyn-Cre virus was injected into the POA (mainly MnPO and VMPO), which retrogradely traveled to the LPB to turn off TeNT expression. Then, AAV-Retro-hSyn-FlpO virus was injected into the DMH, which retrogradely traveled to the LPB to turn on TeNT expression in Cre^-^ neurons, thus blocking neurons that only projected to the DMH but not to the POA (DMH^only^-projecting TeNT). Cre_off-FlpO_on-EGFP was used as the control for Cre_off-FlpO_on-TeNT. (b) Blocking LPB neurons that project to the POA but do not send collateral projections to the DMH (POA^only^-projecting LPB neurons). A similar strategy was used as described in (a) except for the switch of AAV-Retro-hSyn-FlpO and AAV-Retro-hSyn-Cre during the injections. See also Extended Data Fig. 4b for a detailed description. The same control was used as in (a). (c-f) T_core_ changes in response to a series of cold exposures, namely 29→22° C (c), 29→16°C (d), 29→10°C (e), and 29→4°C (f), after blocking POA^only^-projecting LPB neurons or DMH^only^-projecting LPB neurons. DMH^only^-projecting TeNT, n = 8 mice; POA^only^-projecting TeNT, n = 7 mice; GFP controls, n = 10 mice. *, GFP vs. DMH^only^-projecting blocking group. $, DMH^only^ projecting vs. POA^only^ projecting blocking. (g) Blocking LPB-innervating POA/DMH neurons or both using TeNT. The detailed scheme is shown in Extended Data Fig. 5a. Briefly, anterograde transsynaptic Cre carried by AAV1 (AAV1-hSyn-Cre) was injected in the LPB to drive expression of Cre-dependent TeNT injected in either the POA (POA^LPB^ blocking), or the DMH (DMH^LPB^ blocking), or both (co-blocking). Cre-dependent GFP co-injected in both the POA and DMH was used as the control (GFP control). (h-k) T_core_ changes in response to a series of cold exposures, namely 29→22° C (h), 29→16°C (i), 29→10°C (j), and 29→4°C (k), after blocking POA^LPB^, DMH^LPB^, or both types of neurons (n = 10 mice for GFP and DMH^LPB^ blocking group; n = 9 mice for POA^LPB^ blocking group; n = 6 mice for the co-blocking group). *, DMH^LPB^ blocking vs. GFP; $, DMH^LPB^ vs. POA^LPB^ blocking; #, POA^LPB^ vs. co-blocking. All data are the mean ± SEM, and (c-f, h-k) were analyzed by two-way RM ANOVA followed by uncorrected Fisher’s LSD test. The p-values are calculated based on statistical tests in Extended Data Table 2. *p ≤ 0.05; **p ≤ 0.01; ***p ≤ 0.001; ^$^ p ≤ 0.05; ^#^ p ≤ 0.05; ns, not significant.

T_core_ and physical activity were measured by telemetry probes implanted intraperitoneally. Under a thermoneutral condition (29°C), blocking POA^only^-projecting or DMH^only^-projecting LPB^Vglut2^ neurons did not affect body weight, basal T_core_, energy expenditure (EE), or physical activity compared with controls (**Extended Data Fig. 4e-h**). To evaluate cold defense phenotypes, we switched ambient temperature (T_a_) from 29 to 22, 16, 10, or 4°C. When the T_a_ was dropped to 22, 16, or 10°C, blocking these neurons caused significant but similar degrees of hypothermia (**Fig. 3c-e**). When T_a_ was dropped to 4°C, however, blocking POA^only^-projecting neurons resulted in slightly more severe hypothermia than when DMH^only^-projecting neurons were blocked (**Fig. 3f**). This suggests that DMH^only^-projecting and POA^only^-projecting LPB neurons function independently and contribute equally to defense responses to cool/cold temperatures (>10°C), while POA^only^-projecting LPB neurons contribute more to defense responses in cold/nociceptive temperatures (4°C).

### The LPB→POA/DMH pathways form a parallel circuit in cold defense

After showing that the LPB→DMH pathway could function independently of the POA, we tested whether the LPB→POA/DMH pathways function as parallel circuits where these two pathways function independently and cumulatively in cold defense. Thus, we used TeNT to block neurons downstream of the LPB, including LPB-innervating POA neurons, LPB-innervating DMH neurons, or both (**Fig. 3g** **and Extended Data Fig. 5a**). We injected anterograde transsynaptic AAV1-hSyn-Cre^46^ into the LPB and Cre-dependent TeNT into the POA (MnPO, VMPO, MPA, and lateral preoptic area (LPO)), the DMH, or both. These blocked LPB-innervating POA neurons (POA^LPB^ blocking), LPB-innervating DMH neurons (DMH^LPB^ blocking), or both (co-blocking) (**Fig. 3g** **and Extended Data Fig. 5a**). Injecting Cre-dependent GFP into both the POA and the DMH served as the control (GFP control). TeNT expression heatmaps are shown in **Extended Data Fig. 5b**. Under thermoneutral conditions (29°C), no changes in basal T_core_ were seen for any of the blocking groups (**Extended Data Fig. 5c,d,g**). Physical activity did not change in the DMH^LPB^ blocking and co-blocking groups (**Extended Data Fig. 5f,h**), but POA^LPB^ blocking mice slightly increased physical activity in the light phase and basal EE during both the light and dark phases (**Extended Data Fig. 5e,f**).

Under cool/cold conditions (29→22/16/10/4°C switches), all blocking groups showed substantial hypothermia compared to the control (**Fig. 3h-k**). Notably, there were no differences in the hypothermia between DMH^LPB^ blocking and POA^LPB^ blocking groups, which is reminiscent of phenotypes observed after blocking LPB input neurons (**Fig. 3c-f**). However, the co-blocking group always exhibited more substantial hypothermia, suggesting a cumulative or additive effect (**Fig. 3h-k**). Hence, these results demonstrate that the LPB→POA/DMH pathways provide an equivalent and cumulative contribution to cold defense, forming a parallel circuit to maintain T_core_.

We further performed the cold plate test and found no differences in withdrawal latency or the number of lifts between DMH^LPB^ blocking, POA^LPB^ blocking, and controls (**Extended Data Fig. 5i**), suggesting these pathways do not affect cold nociception. In the temperature preference tests, where mice could choose between 30°C and 6, 10, 16, or 35°C for 5 minutes, no preference differences were detected between these groups (**Extended Data Fig. 5j**). Together, these data suggest that the LPB→POA/DMH pathways form parallel circuits to coordinate cold defense. They play a negligible role in cold nociception or thermotactic behaviors.

### The LPB^Vglut2^ → DMH pathway induces strong hyperthermia

Having shown that the LPB→DMH pathway is necessary for cold defense, we sought to test whether this pathway is sufficient to elicit cold defense responses. We injected AAVs carrying Cre-dependent ChR2 into the LPB of Vglut2-Cre mice and photostimulated ChR2-expressing LPB^Vglut2^ neural terminals in the DMH (**Fig. 4a,b**). As expected, photoactivation of LPB^Vglut2^ neural terminals in the DMH increased T_core_ by 1.81°C (**Fig. 4c**), which was accompanied by an abrupt increase in physical activity (**Fig. 4d**). To rule out antidromic propagation of action potentials during terminal stimulation, we chemogenetically inhibited LPB^Vglut2^ somas via hM4D_i_ while photostimulating their terminals in the DMH (**Fig. 4e-f**). We confirmed the inhibition of somas by calcium fiber photometry (**Extended Data Fig. 6a,b**). This inhibition did not affect the induced hyperthermia, suggesting the hyperthermic effect was specific to terminal activation (**Fig. 4g**).

**Fig. 4.**
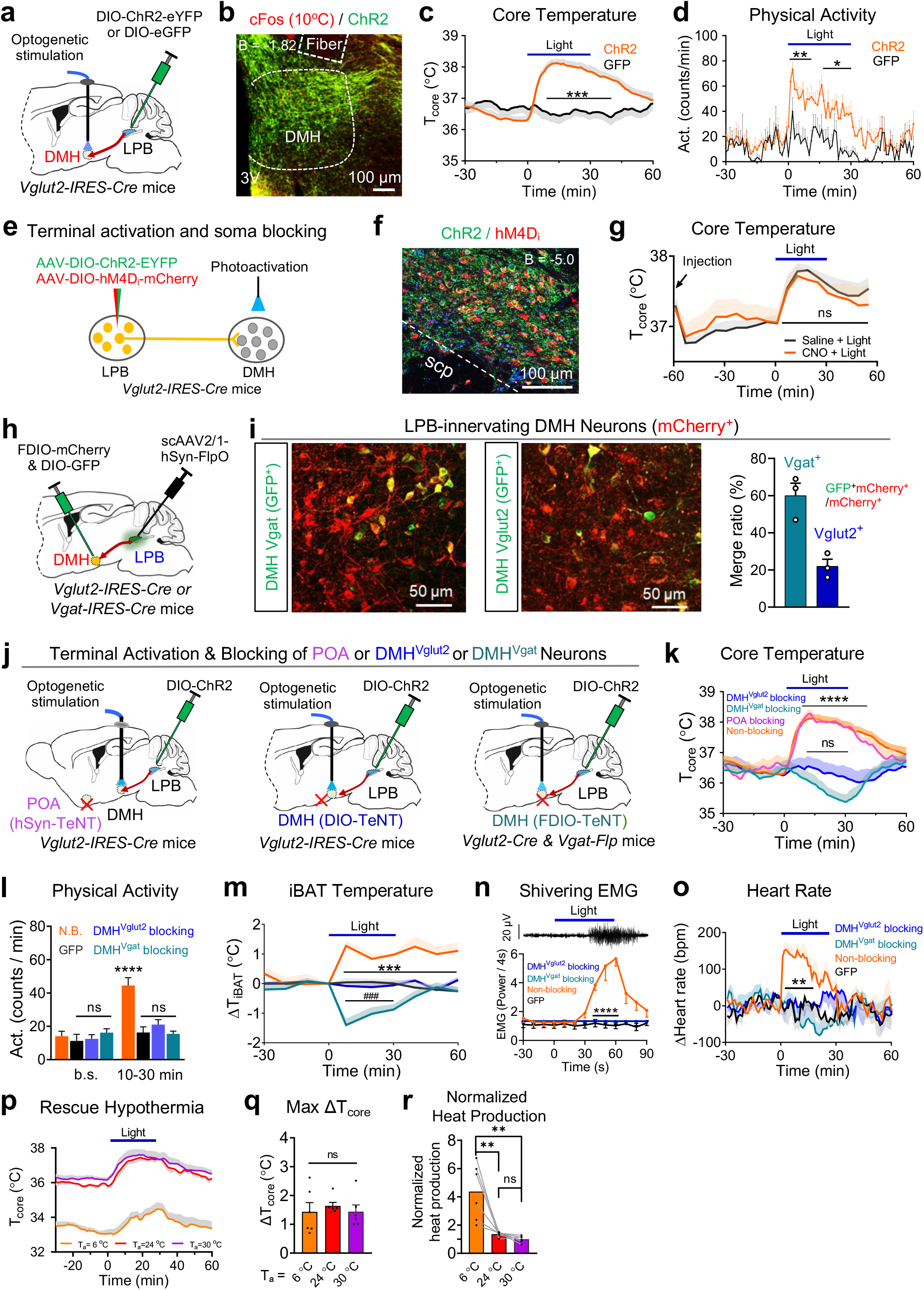
Activation of the LPB^Vglut2^ → DMH projection induces strong cold defense responses independent of the POA. (a) Activating the LPB^Vglut2^ → DMH projection via optogenetic activation of LPB^Vglut2 & ChR2^ neural terminals in the DMH. AAVs carrying Cre-dependent ChR2 (AAV9-hEF1a-DIO-hChR2-EYFP) were injected into the LPB of Vglut2-IRES-Cre mice. An optical fiber was implanted above the DMH and used for optogenetic activation of neural terminals. AAV9-hSyn-Flex-GFP was used as the control. (b) Expression of ChR2 from LPB^Vglut2^ neural terminals in the DMH. Cold-induced cFos (10° C, 2 h) are shown in red. (c-d) Changes of T_core_ (c) and physical activity (d) after photoactivation of LPB^Vglut2 & ChR2^ neural terminals in the DMH (ChR2, n = 10 mice; GFP, n = 8 mice). Light pattern: 473 nm, 12 mW, 10 Hz, 10 ms, 2-s on followed by 2-s off, with the cycles repeating for 30 min (as to g, k-p). (e) Activating the LPB^Vglut2^ → DMH projection while blocking LPB somas with hM4D_i_ . AAVs carrying Cre-dependent ChR2 and Cre-dependent hM4D_i_ (AAV9-EF1α-DIO-hM4D_i_-mCherry) were co-injected into the LPB of Vglut2-IRES-Cre mice. An optical fiber was implanted in the DMH and used for optogenetic activation of neural terminals projecting from LPB^Vglut2^ neurons. (f) Representative ChR2 and hM4Di expression in LPB^Vglut2^ neurons. (g) Changes in T_core_ after activating the LPB^Vglut2^ → DMH projection while blocking LPB neurons (n = 11 mice). CNO was injected at -60 min to silence neurons as indicated (i.p., 10 mg/kg) and saline was used as the control. (h) The neurotransmitter properties of LPB-innervating DMH neurons. Cre-dependent GFP (AAV9-hSyn-DIO-GFP) was injected in the DMH of Vglut2-IRES-Cre or Vgat-IRES-Cre mice to label glutamatergic or GABAergic neurons, respectively. Anterograde trans-synaptic tracing AAV-FlpO (scAAV1-hSyn-FlpO) was injected into the LPB and FlpO-dependent mCherry (AAV9-hSyn-FDIO-mCherry) was injected into the DMH to label LPB-innervating DMH neurons. (i) Representative images showing the overlap between LPB-innervating DMH neurons (mCherry^+^) and GABAergic or glutamatergic markers (GFP^+^). Overlap ratios (GFP^+^ mCherry^+^ /mCherry^+^) are shown on the right (n = 3 mice). GFP^+^ mCherry^+^ /mCherry^+^ in each group refers to the number of double-positive neurons divided by the number of total LPB-innervating DMH neurons. (j) Optogenetic activation of the LPB^Vglut2^ → DMH projection while blocking downstream neurons, including POA neurons, DMH^Vgat^ neurons, or DMH^Vglut2^ neurons. Cre-dependent ChR2 was injected into the LPB of Vglut2-IRES-Cre mice. An optical fiber was implanted above the DMH and used for optogenetic activation of neural terminals from LPB^Vglut2 & ChR2^ neurons. To block POA neurons, pan-neuronal expressing TeNT (AAV9-hSyn-eGFP-TeNT) was injected into the POA. To block DMH^Vglut2^ neurons, Cre-dependent TeNT (AAV9-hSyn-DIO-mCherry-TeNT) was injected in the DMH of Vglut2-IRES-Cre mice. To block DMH^Vga t^ neurons, FlpO-dependent TeNT (AAV9-hSyn-FDIO-mCherry-TeNT) was injected in the DMH of Vglut2-IRES-Cre & Vgat-T2A-FlpO mice. (k) Changes in T_core_ after photoactivation of the LPB^Vglut2^ → DMH projection while blocking POA neurons, DMH^Vgat^ neurons, or DMH^Vglut2^ neurons (n = 5 mice each). The non-blocking group was replotted from Fig. 4c. *, the non-blocking group vs. DMH^Vglut2^ blocking group; ns, DMH^Vglut2^ blocking group vs. DMH^Vgat^ blocking group. (l-o) Blocking of DMH^Vglut2^ or DMH^Vgat^ neurons abolished photoactivation-induced increases in physical activity (l; n = 5 mice for each blocking and GFP group, n = 10 mice for non-blocking group), iBAT thermogenesis (m; n = 5 mice each), nuchal muscle shivering EMG (n; n = 4 mice each), and heart rate (o; n = 5 mice each). *, non-blocking (N.B.) compared to DMH^Vglut2^ blocking group (m-o); #, DMH^Vglut2^ blocking group compared to DMH^Vgat^ blocking group (m). b.s. and (10 - 30 min) represent the averaged physical activity between t = (- 30 - 0 min) and t = (10 - 30 min), respectively. (p-q) Changes in T_core_ after photoactivation of the LPB^Vglut2^ → DMH projection under different T_a_ (6, 24, 30°C; n = 6 mice each) (p). The maximum ΔT_core_ during photoactivation was quantified in (q). (r) Normalized heat production of mice after photoactivation of the LPB^Vglut2^ → DMH projection under different T_a_ as indicated. Heat production was normalized by the formula (T_core_ -T_a_) / ΔT_core_ as reported^47^. All data are the mean ± sem, and were analyzed by two-way RM ANOVA followed by Bonferroni’s multiple comparisons tests (c, g, k-o), or by uncorrected Fisher’s LSD test (d), (q and r) were analyzed by paired t-test. The p-values are calculated based on statistical tests in Extended Data Table 2. *p ≤ 0.05; **p ≤ 0.01; ***p ≤ 0.001; ****p ≤ 0.0001; ^###^ p ≤ 0.001; ns, not significant.

### Cold-defense responses induced by activating the LPB^Vglut2^ → DMH pathway depend on DMH^Vglut2/Vgat^ neurons but not POA neurons

To identify DMH neuronal cell types targeted by the LPB, we used the anterograde transsynaptic AAV1 tracer to label LPB-innervating DMH neurons and co-stained with GABAergic and glutamatergic markers in the DMH. To do so, we simultaneously injected AAV1-hSyn-FlpO into the LPB, and a mixture of AAVs carrying FlpO-dependent mCherry (AAV8-FDIO-mCherry) and Cre-dependent GFP (AAV9-DIO-GFP) into the DMH, in either Vglut2-IRES-Cre or Vgat-IRES-Cre mice (**Fig. 4h**). Interestingly, the mCherry (LPB-innervating DMH neurons) overlapped with both Vgat^+^ (∼60%) and Vglut2^+^ markers (∼20%) (**Fig. 4i**), suggesting that both GABAergic and glutamatergic DMH neurons are innervated by the LPB.

To determine which DMH neural types are required for the hyperthermic function of the LPB^Vglut2^ → DMH pathway, we blocked DMH^Vglut2/Vgat^ neurons using TeNT while photoactivating LPB^Vglut2^ & ^ChR2^ terminals in the DMH (**Fig. 4j** **and Extended Data Fig. 6c-h**). The Vgat-T2A-FlpO used for labeling DMH^Vgat^ neurons was validated by GABA immunostaining (**Extended Data Fig. 6g**). As a comparison, we also blocked POA neurons using TeNT expressed in the MnPO, VMPO, MPA, and LPO (**Fig. 4j** **and Extended Data Fig. 6i-j**). As expected, blocking DMH^Vglut2^, DMH^Vgat^, or POA neurons rendered mice intolerant to cold exposure (4°C) (**Extended Data Fig. 6c-k**); this is consistent with the known functions of these neurons^14^. Interestingly, blocking either DMH^Vglut2^ or DMH^Vgat^ neurons abolished the hyperthermia induced by photoactivating LPB^Vglut2^ terminals in the DMH (**Fig. 4k**). In contrast, POA blocking did not affect this induced hyperthermia (**Fig. 4k**). Unexpectedly, photoactivation caused mild hypothermia after blocking DMH^Vgat^ neurons (**Fig. 4k**), which may be caused by activating unknown hypothermic neural axons. Therefore, the hyperthermic effect of the LPB^Vglut2^ → DMH pathway depends on DMH^Vglut2/Vgat^ neurons but not POA neurons.

To further determine which autonomic cold-defense responses are recruited by the LPB^Vglut2^ → DMH pathway, we measured physical activity and heart rate by telemetry probes, interscapular BAT temperature (T_iBAT_) by infrared thermography, and muscle shivering by nuchal muscle electromyography (EMG). Interestingly, photoactivation was sufficient to increase physical activity, T_iBAT_, shivering EMG, and heart rate (**Fig. 4l-o**). Blocking either DMH^Vglut2^ or DMH^Vgat^ neurons abolished these responses, suggesting that both neural types are essential for these defense responses. Like T_core_ changes, T_iBAT_ decreased in response to photoactivation after blocking DMH^Vgat^ neurons (**Fig. 4m**).

### Activation of the LPB^Vglut2^ DMH pathway rapidly relieves hypothermia

As activation of the LPB^Vglut2^ → DMH pathway induced robust hyperthermic responses, we wondered whether it could alleviate cold-induced hypothermia. Notably, photoactivation of LPB^Vglut2 & ChR2^ terminals in the DMH substantially relieved the hypothermia observed after placing mice at T_a_ of 6°C (**Fig. 4p**). Surprisingly, T_core_ increases were similar for mice placed at T_a_ of 6, 24, or 30°C (**Fig. 4q**), even though heat loss was much greater at lower T_a_ ^47^. Therefore, we normalized heat production by considering the heat-loss effects and found that heat production increased 4-fold at T_a_ of 6°C compared to that seen at T_a_ of 24°C (**Fig. 4r**). These results collectively suggest that the LPB^Vglut2^ → DMH pathway robustly restores T_core_ in mice subjected to cold.

### Activation of the LPB^Vglut2^ → DMH pathway increases iBAT thermogenesis and suppresses body weight gain

An increase in T_iBAT_ may indicate BAT thermogenesis or may only mirror the increase in T_core_. To differentiate between these possibilities, we increased the temporal resolution in T_core_ and T_iBAT_ recording using custom-made wired probes^25^ (**Fig. 5a**). After photoactivating LPB^Vglut2^ → DMH projections, the rise of T_iBAT_ preceded T_core_ and reached a higher value than T_core_ (**Fig. 5b**), suggesting that T_iBAT_ could be a driving force of T_core_ changes. Further, the denervation of iBAT sympathetic nerves suppressed the hyperthermia induced by photoactivation (**Fig. 5b**), suggesting an essential function of the iBAT. Additionally, the expression of thermogenin uncoupling protein 1 (UCP1) in the iBAT was elevated 3-h post photoactivation (**Fig. 5c**). These results collectively show that iBAT thermogenesis is essential for the hyperthermic function of the LPB^Vglut2^ → DMH pathway.

**Fig. 5.**
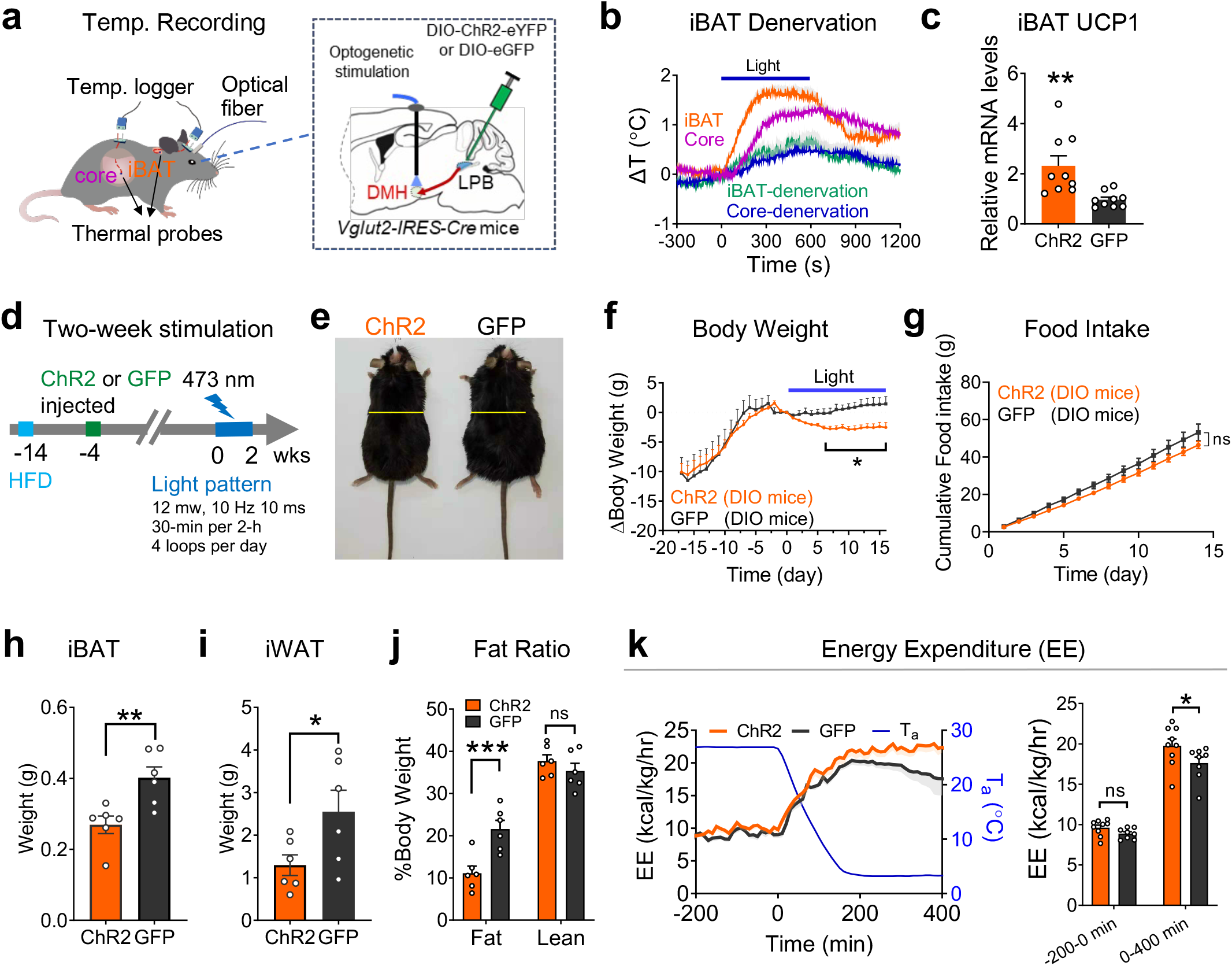
Activation of the LPB^Vglut2^ → DMH projection increases iBAT thermogenesis and suppresses body weight gain. (a) Simultaneously recording T_core_ and T_iBAT_ in freely behaving mice at high temporal resolution using wired probes while activation of the LPB^Vglut2^ → DMH projection. Both temperatures were recorded via small T-type thermal probes connected to the logger via T-type wires. AAVs carrying Cre-dependent ChR2 were injected into the LPB of Vglut2-IRES-Cre mice. An optical fiber was implanted above the DMH and used for optogenetic activation of neural terminals. AAV9-hSyn-Flex-GFP was used as the control. (b) Changes in T_core_ and T_iBAT_ in response to photoactivation of the LPB^Vglut2^ → DMH projection in the sham group and iBAT denervation group (n = 5 mice each). The denervation group had the iBAT sympathetic nerves cut beneath the iBAT pads. (c) Relative UCP1 expression levels in iBAT 3-h after photoactivation of the LPB^Vglut2^ → DMH projection (n = 9 mice each). Light pattern: 473 nm, 12 mW, 10 Hz, 10 ms, 2-s on 2-s off, 30 min. (d) Testing the effect of two-week photoactivation of the LPB^Vglut2^ → DMH projection on body weight of DIO mice fed with HFD. Light pattern is indicated. (e) Mice after two weeks of photoactivation (left, ChR2 group; right, GFP group). (f-g) Changes in body weight (f) and cumulative food intake (g) during two weeks of photoactivation (ChR2, n = 9 mice; GFP, n = 5 mice). The light pattern is shown in (d). (h-k) Changes in iBAT weight (h), iWAT weight (i), body composition (j), and cold-induced energy expenditure (k) after two-week photoactivation (h-j, n = 6 mice each; k, ChR2, n = 9 mice; GFP, n = 8 mice). T_a_ was changed from 25 to 4° C in (k). All data are the mean ± sem, and (f, g) were analyzed by two-way RM ANOVA followed by Bonferroni’s multiple comparisons tests, (c, and h-k) were analyzed by unpaired t-test, The p-values are calculated based on statistical tests in Extended Data Table 2. *p ≤ 0.05; **p ≤ 0.01; ns, not significant.

These increases in BAT thermogenesis prompted us to test whether long-term activation of this pathway would increase energy expenditure to lower body weight. Thus, we used a two-week photostimulation protocol to activate the LPB^Vglut2^ → DMH projection in diet-induced obesity (DIO) mice fed with a high-fat diet (HFD) for 14 weeks (**Fig. 5d**). Although the photostimulation procedure itself appeared to curb weight gains in control mice, photoactivation of the LPB^Vglut2^ → DMH projection further reduced body weight without affecting cumulative food intake (**Fig. 5e-g**). Consistently, photoactivation decreased the weights of iBAT and inguinal white adipose tissue (iWAT), reduced the adipose ratio, and increased EE during cold exposure (**Fig. 5h-k**). Therefore, chronic activation of this pathway increases EE and lowers body weight.

### Projection-specific transcriptomic analysis identifies DMH-projecting LPB^SST^ neurons as cold-activated neurons

After showing that the LPB^Vglut2^ → DMH pathway plays important roles in cold defense, we sought to identify neural types that comprise this pathway. We used a cell type- and projection-specific translating ribosome affinity purification sequencing technique (retroTRAP-seq) we developed^25^, where retrograde AAVs carrying DIO-GFPL10 were injected into the DMH of Vglut2-Cre mice to express GFP-tagged ribosomal protein L10 in the LPB (**Fig. 6a**). The LPB was microdissected and GFP-tagged ribosomes were immunoprecipitated (IP) and used for subsequent sequencing. We analyzed the enrichment fold (IP/input) and found that 413 genes were upregulated (**Fig. 6b** **and Extended Data Table 1**). We compared our upregulated genes with PB-expressed genes obtained from the Allen Brain database and found several genes that overlapped, including *N4bp2os*, *galanin* (*Gal*), *SST*, and *Spint2* (**Fig. 6c**). Since a Cre strain was not available to label *n4bp2os^+^* neurons, and *Gal* was not expressed in the LPB^25^, we used SST-IRES-Cre to label *SST*^+^ neurons and performed further functional studies.

**Fig. 6.**
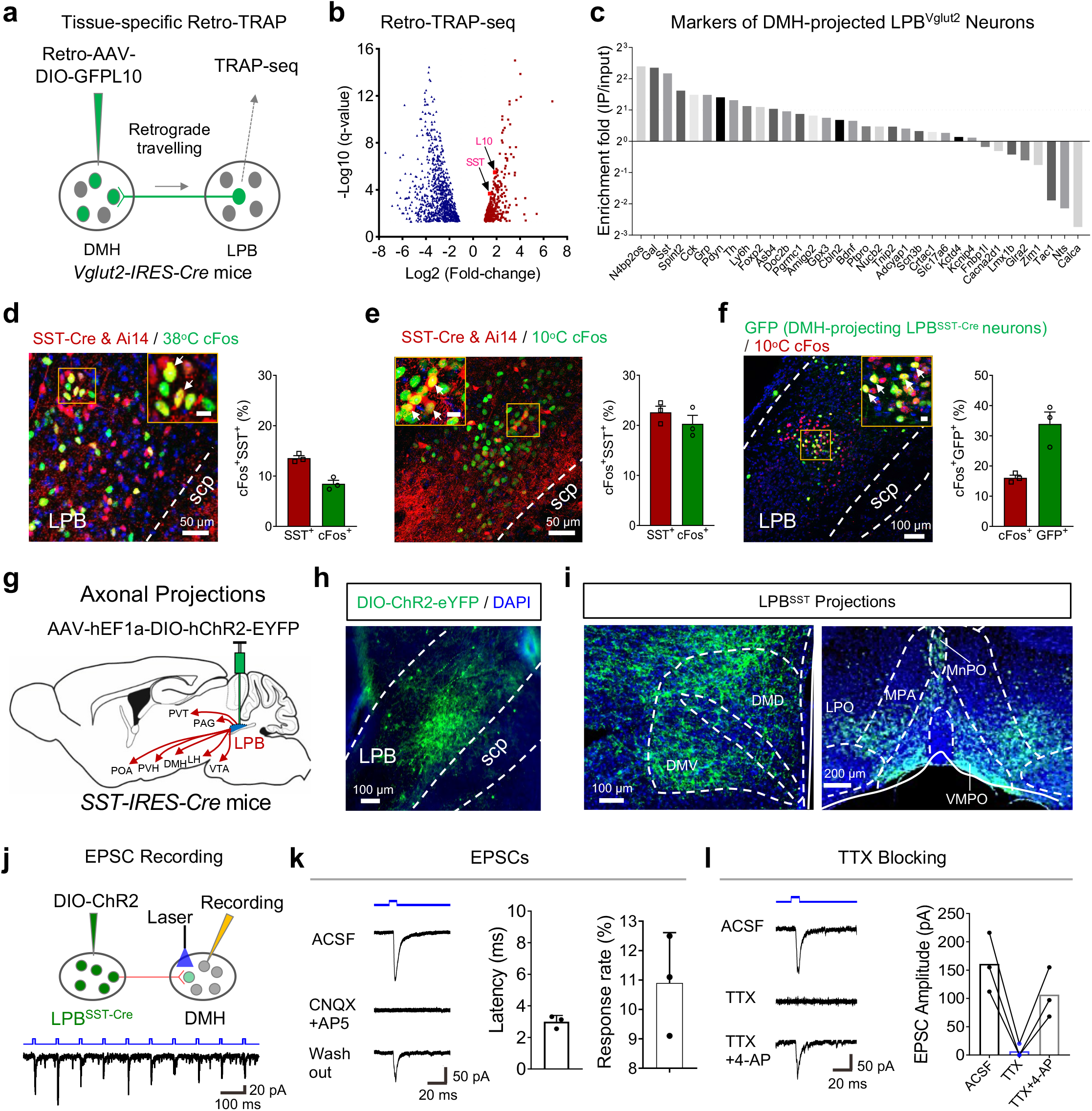
Projection-specific transcriptomic analysis identifies DMH-projecting LPB^SST^ neurons as cold-activated neurons. (a) Projection-specific transcriptomic analysis (retro-TRAP), where GFP-tagged translational ribosomes from DMH-projecting LPB^Vglut2^ neurons were immunoprecipitated and associated mRNAs were sequenced. Retrograde tracing virus carrying Cre-dependent GFP-tagged ribosomal protein L10 (AAV-Retro-hEF1a-FLEX-GFPL10) was injected into the DMH, which traveled to the LPB and expressed GFPL10 after recombination by Vglut2-IRES-Cre. SST, somatostatin. (b) Volcano plots (q value versus log2 fold change) for LPB mRNAs after retro-TRAP sequencing. (c) Retro-TRAP fold enrichment (IP/Input) for PB-expressed genes downloaded from the Allen Institute. (d-e) Overlap between SST-IRES-Cre labeled neurons (Tdt^+^) in the LPB and the following immuno-positive neurons: warm-induced cFos (10° C, 2 h; d), and cold-induced cFos (38° C, 2 h; e). SST-IRES-Cre mice were crossed with Ai14 (Rosa-CAG-LSL-tdTomato-WPRE) to label SST-IRES-Cre. Overlap ratios are shown on the right of each panel (n = 3 mice each). (f) Overlap between DMH-projecting LPB^SST^ neurons and cold-induced cFos (10° C, 2 h) (n = 3 mice each). To label DMH-projecting LPB^SST^ neurons, we injected retrograde AAVs carrying Cre-dependent FlpO (AAV-Retro-CAG-Flex-FlpO) in the DMH of SST-IRES-Cre mice, which drove the expression of FlpO-dependent GFP (AAV9-hEF1a-FDIO-EGFP) in the LPB. (g) Axonal tracing of LPB^SST^ neurons by ChR2, where AAV9-hEF1a-DIO-hChR2-EYFP was injected into the LPB of SST-IRES-Cre mice. PVH, paraventricular hypothalamic. (h-i) Expression of ChR2 in LPB^SST^ neural somas (h) and terminals in the POA or DMH (i). (j) Induction of EPSCs in DMH neurons by light stimulation of LPB^SST & ChR2^ terminals (blue, 6 mW, 10 ms, 10 hz). (k) EPSCs were blocked by GluR antagonists D-AP5 and CNQX. Latency and response rate (3 of 28 randomly recorded neurons from 3 mice) are shown on the right. (l) EPSCs were blocked after TTX treatment but were restored by 4-AP treatment. All data are the mean ± sem. Scale bars in the enlarged panel of (d–f) are 10 μm.

SST-Cre labeled neurons largely overlapped with SST immunoactivity (overlapped/SST^+^ = 89%; **Extended Data Fig. 7a**) and glutamate immunostaining^48^ (overlapped/glu^+^ = 99%; **Extended Data Fig. 7b**), suggesting that this Cre line faithfully labeled *SST*^+^ neurons and that these neurons were glutamatergic. Quantifying the overlapping between LPB^SST^ neurons and cold/warm-induced cFos suggested that LPB^SST^ neurons accounted for only 8% of the neurons activated by warmth in the LPB (38°C, 2 h), but 20% of the neurons activated by cold (10°C, 2 h) (**Fig. 6d**,**e**). Also, more LPB^SST^ neurons were activated in response to cold (∼22%) than to warmth (∼13%) (**Fig. 6d**,**e**). About 33% of DMH-projecting LPB^SST^ neurons were sensitive to cold exposure, accounting for 16% of cold-activated LPB neurons (**Fig. 6f**). As a comparison, DMH-projecting LPB^Vglut2^ neurons accounted for 38% of cold-activated LPB neurons (**Fig. 1h**,**i**). Using SST-Cre, we mapped the projection pattern of LPB^SST^ neurons by injecting Cre-dependent ChR2 into the LPB (**Fig. 6g,h and Extended Data Fig. 7c**). As expected, LPB^SST^ neurons projected to the DMH and other regions, including the MnPO, VMPO, and LPO (**Fig. 6i** **and Extended Data Fig. 7d**). Thus, our data indicate that glutamatergic LPB^SST^ neurons exhibit a biased response to cold temperature, send projections to the DMH, and comprise up to ∼40% (16%/38%) of cold-activated neurons within the LPB→DMH pathway.

### Monosynaptic connections between LPB^SST^ and DMH neurons

To test whether there is a direct connection between LPB^SST^ and DMH neurons, we expressed ChR2 in LPB^SST^ neurons (**Fig. 6h**,**i**) and found that light stimuli induced EPSCs from DMH neurons (**Fig. 6j**). These currents were blocked by glutamate receptor antagonists (CNQX and AP5), and the latency after photoactivation was within the range of a monosynaptic connection (**Fig. 6k**, left two panels). Noticeably, only ∼10% of randomly recorded DMH neurons exhibited EPSCs in response to light stimulation of SST^+^ terminals (**Fig. 6k**, right), making up ∼1/3 of total LPB-innervating DMH neurons (**Fig. 1k**). To further verify this connection was monosynaptic, we first blocked EPSCs with tetrodotoxin (TTX) and then applied 4-aminopyridine (4-AP) to sensitize the postsynaptic current (**Fig. 6l**). Indeed, 4-AP restored EPSCs blocked by TTX, suggesting that LPB^SST^ and DMH neurons form monosynaptic connections.

### The LPB^SST^ → DMH pathway increases T_core_ via iBAT thermogenesis and is required for cold defense

To investigate the function of the LPB^SST^ → DMH pathway, we optogenetically activated LPB^SST & ChR2^ terminals in the DMH (**Fig. 7a,b**). T_core_ and physical activity were measured by telemetry probes and T_iBAT_ was measured by infrared thermography. As expected, photoactivation of these terminals increased T_core_ (∼1.1°C) and T_iBAT_, but not physical activity (**Fig. 7c-e**). To test whether iBAT thermogenesis was required for this hyperthermia, we denervated iBAT sympathetic nerves and found it abolished the hyperthermic effect (**Fig. 7f**). In contrast to activation of the LPB^Vglut2^ → DMH projection, activation of the LPB^SST^ → DMH projection did not affect muscle shivering and heart rate (**Fig. 7g, h**). To rule out a contribution of POA projections from LPB^SST^ neurons to hyperthermic responses, we used retrograde traveling AAVs carrying Cre-dependent FlpO to drive TeNT expression to block POA-projecting LPB^SST^ neurons (**Fig. 7i, j**). Indeed, blocking POA-projecting LPB^SST^ neurons did not affect the increases in T_core_ and T_iBAT_ induced by photoactivation of the LPB^SST^ → DMH projection (**Fig. 7k, l**).

**Fig. 7.**
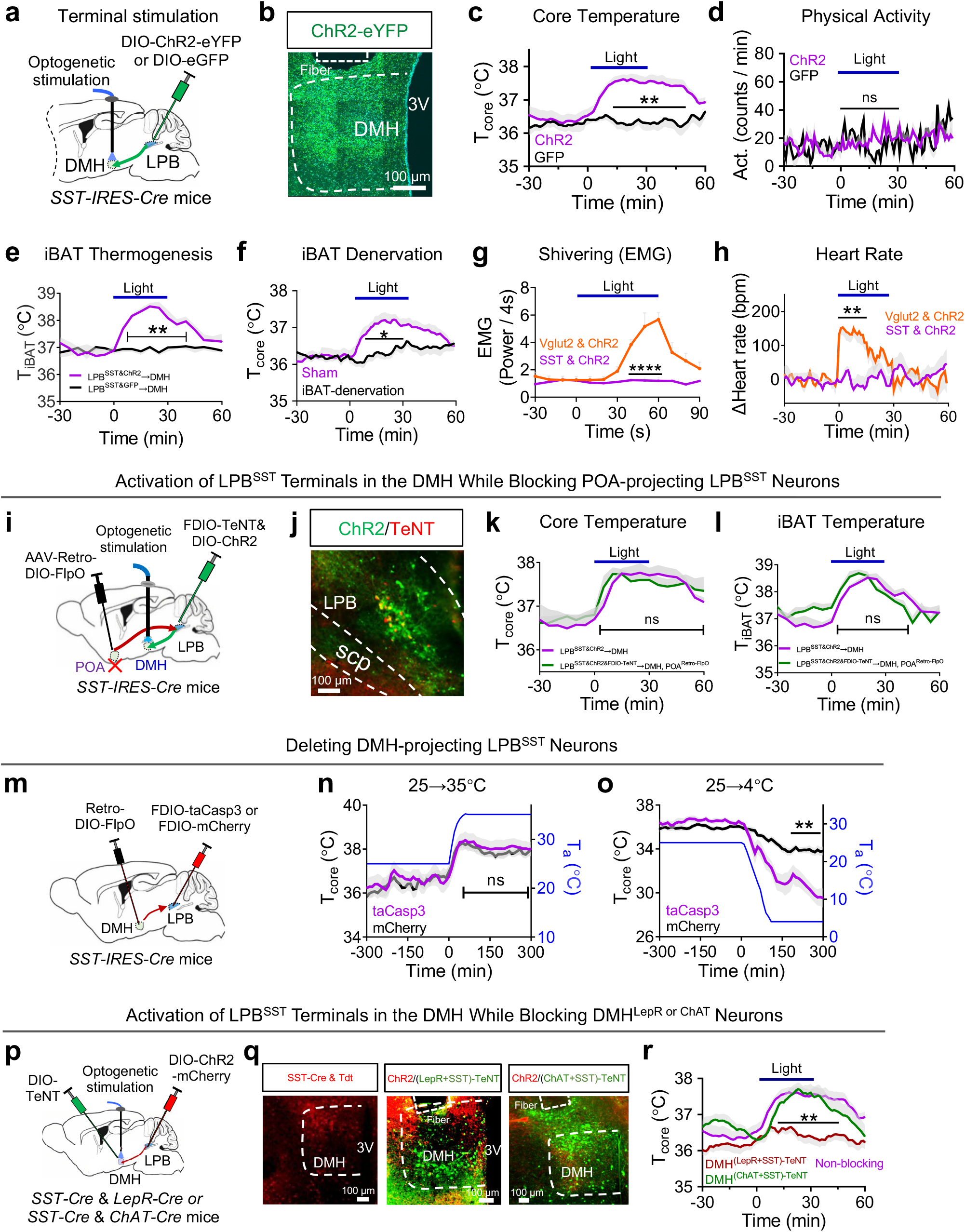
| The LPB^SST^ → DMH projection is selectively required for iBAT thermogenesis during cold defense. (a-b) Design to activate the LPB^SST^ → DMH projection via photostimulating of LPB^SST & ChR2^ terminals in the DMH. AAV9-hEF1a-DIO-hChR2-EYFP was injected into the LPB of SST-IRES-Cre mice. An optical fiber was implanted in the DMH and used to optogenetically activate neural terminals projecting from LPB^SST^neurons. The eGFP (AAV9-hSyn-Flex-GFP) was used as the control for ChR2. The representative expression of ChR2-eYFP in the DMH is shown in (b). (c-e) Changes in T_core_ (c; ChR2, n = 8 mice; GFP, n = 7 mice), physical activity (d; ChR2, n = 20 mice; GFP, n = 9 mice), and T_iBAT_ (e; ChR2, n = 6 mice; GFP, n = 7 mice) after photoactivation of the LPB^SST^ → DMH projection. Light pattern: 473 nm, 6 mW, 10 Hz, 10 ms, 2-s on followed by 2-s off, with the cycles repeating for 30 min. (f) Denervation of the iBAT sympathetic nerves abolished the hyperthermia induced by photoactivation of the LPB^SST^ → DMH projection (n = 6 mice each). (g-h) Photoactivation of the LPB^SST^ → DMH projection did not change nuchal muscle EMG (g; n = 4 mice each) and heart rate (h; SST, n = 8 mice; Vglut2, n = 5 mice). For comparison, the EMG activity and heart rate after photoactivation of LPB^Vglut2^ terminals in the DMH were replotted from Fig. 4n,o. (i) Activating the LPB^SST^ → DMH pathway while blocking POA-projecting LPB^SST^ neurons using TeNT. AAV9-hEF1a-DIO-hChR2-EYFP was injected into the LPB of SST-IRES-Cre mice. An optical fiber was implanted in the DMH and used to optogenetically activate neural terminals projecting from LPB^SST^ neurons. To block POA-projecting LPB^SST^ neurons, retrograde AAVs carrying Cre-dependent FlpO (AAV-Retro-CAG-DIO-FlpO) were injected into the POA of SST-IRES-Cre mice, which drove the expression of FlpO-dependent TeNT (AAV9-hEF1a-FDIO-mCherry-2A-TeNT) in the LPB. (j) Representative expression of ChR2 (green) and TeNT (red) in the LPB. (k-l) Changes in T_core_ (k) and T_iBAT_ (l) after photoactivation of LPB^SST^ terminals in the DMH while blocking POA-projecting LPB^SST^ neurons (n = 6 mice each). Data of the LPB ^SST&CHR2^ → DMH group were replotted from (c) and (e), respectively. (m) Deleting DMH-projecting LPB^SST^ neurons by neural killing with taCasp3. Retrograde AAVs carrying Cre-dependent FlpO (AAV-Retro-CAG-DIO-FlpO) were injected in the DMH, which drove the expression of FlpO-dependent taCaspase3 (AAV9-hEF1a-FDIO-taCasp3-TEVp) injected in the LPB. mCherry (AAV8-hEF1a-FDIO-mCherry) injected in the LPB was used as the control. (n-o) T_core_ changes during warm (25→35°C, n) and cold (25→4° C, o) exposures after deleting DMH-projecting LPB^SST^ neurons (n = 8 mice each). T_a_ changes are indicated. (p) Photoactivation of LPB^SST & ChR2^ terminals in the DMH after blocking DMH^LepR+SST^ neurons or DMH^ChAT+SST^ neurons using neurotoxin TeNT. To block neurons, AAV5-hSyn-DIO-GFP-P2A-TeNT was injected in the DMH of SST-Cre & LepR-Cre or SST-Cre & ChAT-Cre mice. (q) Representative SST-Cre & Tdt (left panel) and LPB^SST & ChR2^ terminals (red) and TeNT (green) expression in the DMH (right two panels). (r) Changes in T_core_ after photoactivation of LPB^SST^ terminal in the DMH while blocking DMH^LepR+SST^ or DMH^ChAT+SST^neurons (n = 7 mice each). The non-blocking group data of T_core_ were replotted from (c). All data are the mean ± sem, and (c-h, k-l, n-o, r) were analyzed by two-way RM ANOVA followed by Bonferroni’s multiple comparisons test. The p-values are calculated based on statistical tests in Extended Data Table 2. *p ≤ 0.05; **p ≤ 0.01; ****p ≤ 0.0001; ns, not significant.

Furthermore, we modulated the activity of LPB^SST^ somas via chemogenetics. Similar to terminal activation, activation of LPB^SST^ somas via hM3D_q_ caused increases in T_core_ but not physical activity (**Extended Data Fig. 8a-d**). Chemogenetic activation also increased EE (**Extended Data Fig. 8e**). Together, we found that the LPB^SST^ → DMH projection within the LPB→DMH pathway increases T_core_ and EE via BAT thermogenesis selectively.

To test the necessity of the LPB^SST^ → DMH pathway in thermoregulation, we blocked DMH-projecting LPB^SST^ neurons via a projection-specific lesion strategy that takes advantage of taCaspase3 (**Fig. 7m**). As expected, this lesion did not affect T_core_ after warm exposure (35°C) (**Fig. 7n**). However, it impaired thermoregulation after cold exposure (4°C) (**Fig. 7o**). Together, these results demonstrate that the LPB^SST^ → DMH pathway is required for cold defense by selectively controlling iBAT thermogenesis.

### LPB^SST^ neurons target DMH^LepR^ neurons to increase T_core_

To determine the DMH cell types that were targeted by LPB^SST^ neurons to increase T_core_, we considered neurons expressing leptin receptor (LepR) and ChAT since they both regulate thermogenesis^36, 38^. To block these two types of neurons while photoactivating LPB^SST^ → DMH projections, we crossed SST-Cre with LepR-Cre or ChAT-Cre to obtain double Cre-positive mice, and then injected DIO-ChR2 and DIO-TeNT into the LPB and DMH, respectively (**Fig. 7p,q**). Although LepR-Cre and ChAT-Cre may also drive ChR2 expression in the LPB, LPB^LepR^ neurons induce very mild hyperthermia^25^ and LPB^ChAT^ neurons barely project to the DMH (Allen Brain Atlas). Therefore, photoactivation of the LPB^SST+LepR/ChAT^ → DMH projection would largely recapitulate the hyperthermia phenotype observed after photoactivating the LPB^SST^ → DMH projection. Then, we photoactivated these terminals while blocking LepR+SST or ChAT+SST neurons in the DMH, since SST-Cre is also expressed in the DMH (**Fig. 7q**, left panel). Interestingly, blocking DMH^LepR+SST^ neurons blunted the induced hyperthermia, whereas blocking DMH^ChAT+SST^ neurons had no effect (**Fig. 7r**). Results from these blocking experiments are consistent with the function of DMH^LepR^ neurons, which increase BAT thermogenesis^36^. Therefore, we suggest that DMH^LepR^ neurons function as downstream targets of LPB^SST^ neurons to increase BAT thermogenesis.

### The raphe pallidus nucleus (RPa) functions downstream of the LPB**→**DMH pathway

In seeking to determine downstream targets of the LPB→DMH pathway, we considered the RPa since it is a synaptic target of DMH thermogenic neurons^10^. Besides, RPa-projecting DMH Vglut2^+^ and Brs3^+^ neurons receive synaptic inputs from the LPB^35, 40^, suggesting the existence of an LPB→DMH→RPa pathway. To demonstrate the functional importance of this pathway, we photostimulated LPB^Vglut2^ ^&^ ^ChR2^ terminals in the DMH while blocking RPa-projecting DMH neurons (**Extended Data Fig. 9a**). To do so, we infected LPB^Vglut2^ somas with AAVs carrying DIO-ChR2, and injected a retrograde AAV carrying FlpO into the RPa to induce the expression of FlpO-dependent TeNT in the DMH (**Extended Data Fig. 9a,b**). The mCherry was used as a control for TeNT. Blocking RPa-projecting DMH neurons itself did not affect basal body temperature or physical activity compared with controls measured during the light cycle (**Extended Data Fig. 9c**). Next, we photoactivated LPB^Vglut2^ ^&^ ^ChR2^ terminals in the DMH while blocking RPa-projecting DMH neurons. Strikingly, the hyperthermia induced by photoactivation was abolished (**Extended Data Fig. 9d**), as well as autonomic cold defense responses associated with photoactivation, including physical activity, iBAT thermogenesis, and muscle shivering (**Extended Data Fig. 9e-g**). Hence, we show that RPa-projecting DMH neurons are required for the LPB→DMH pathway to mediate cold-defense function. We, therefore, suggest that the RPa is the primary downstream target of the LPB→DMH pathway to promote BAT thermogenesis and muscle shivering since the RPa is known to regulate these activities downstream of the DMH^10, 35, 40^. Yet, a concern was raised about whether physical activity changes were also mediated by the RPa since it has not been reported before.

We then tested whether the RPa plays a role in regulating physical activity by neural blocking (**Extended Data Fig. 9h,i**). Consistent with its role in thermogenesis, blocking RPa neurons via TeNT decreased the basal T_core_ during the dark cycle compared with GFP controls (**Extended Data Fig. 9j**). As expected, the hyperthermia induced by photoactivation of LPB^Vglut2^ ^&^ ^ChR2^ neural terminals in the DMH was abolished after blocking RPa neurons (**Extended Data Fig. 9k**), further supporting that the RPa is the downstream target of the LPB→DMH pathway. To our surprise, blocking the RPa itself caused a compulsive-like circling behavior in the cage (**Extended Data Fig. 9l**), which resulted in hyperactivity and prevented us from judging its role in regulating physical activity during cold defense. Therefore, the roles of the RPa in regulating physical activity require further studies.

## Discussion

Hypothermia caused by cold or starvation is a major threat to life^4, 5, 49^. Therefore, the ability to prevent hypothermia under normal conditions and to rapidly recover from a bout of hypothermia is critical for animal survival and fitness. Here, we found that a previously uncharacterized pathway in mice, namely the LPB→DMH pathway, functions in parallel to the LPB→POA pathway to rapidly increase thermogenesis during cold defense and quickly recover from hypothermia caused by cold exposure in mice (**Fig. 4**). This parallel-circuit model (illustrated in **Fig. 8**) deepens our understanding of how neuronal circuits may affect thermoregulation and may represent a basic network in regulating other essential homeostatic behaviors.

**Fig. 8.**
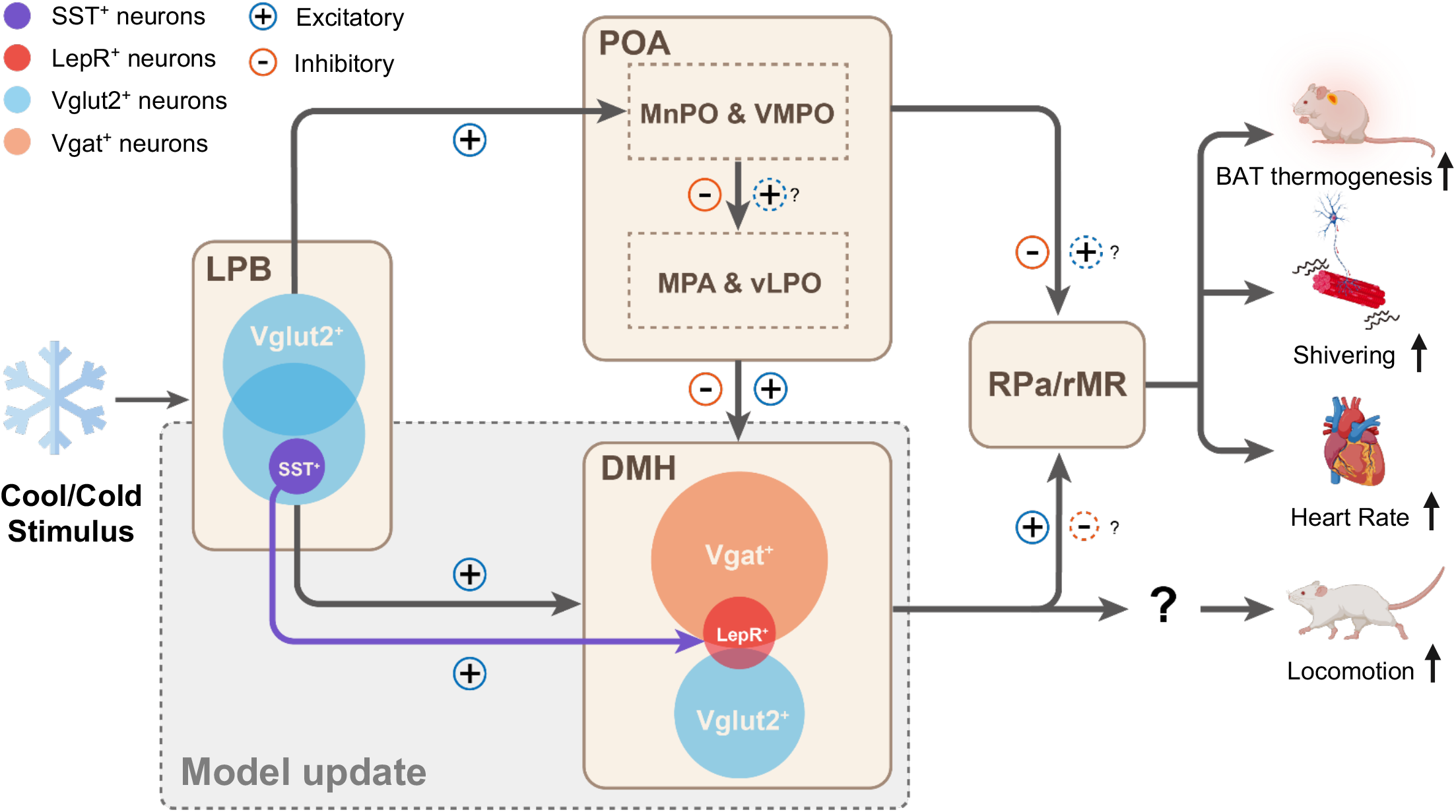
Proposed parallel circuits for cold defense. The LPB→POA→DMH/RPa circuitry has been proposed before^10^ and the model updated by this study is shaded in grey color. Afferent cool/cold signals activate Vglut2^+^ neurons in the LPB. Among these cold-activated neurons (cFos^+^ ; 100%), 45% projected to the POA (mainly the MnPO, VMPO, and LPO), 38% projected to the DMH, and 20% projected to both regions. Projection-specific neural blockings suggest that both the LPB →POA and LPB→DMH pathways are required in cold defense, where the two pathways contribute equivalently and cumulatively to cold defense and therefore form a parallel circuit. Within the LPB→DMH pathway, 60% of LPB-innervated DMH neurons are Vgat^+^, while 20% of them are Vglut2^+^ . Activation of the LPB→DMH pathway induces strong cold-defense responses, including increases in BAT thermogenesis, muscle shivering, heart rate, and locomotion. Both the DMH^Vgat^ and DMH^Vglut2^ neurons are required to support the cold-defense function of the LPB→DMH pathway. Additionally, a subpopulation of SST^+^ neurons in the LPB targets DMH^LepR^ neurons to promote BAT thermogenesis selectively, suggesting a genetically defined neural projection controls specific cold-defense activities. Downstream of the DMH, the RPa or rostral medullary raphe region (rMR) is known to regulate BAT thermogenesis, muscle shivering, and heart rate, while the regulation of locomotion is not clear.

### The role of a parallel circuit in cold defense

The LPB→POA pathway has been suggested to be important for cold defense in rats. Pharmacological activation of LPBel neurons with N-methyl-D-aspartic acid (NMDA) increases BAT thermogenesis and HR in rats, and these phenotypes are suppressed by antagonizing glutamate receptors in the MnPO^39^. Consistent with this, our data suggest that LPB neurons mainly innervate the MnPO, VMPO, and LPO in mice (**Fig. 1b**). However, recent optogenetic manipulations in mice failed to verify a cold-defense function of the LPB→POA pathway, whose activation caused hypothermia alone^25, 26^. These conflicting results raise the issue of whether there are species-specific roles played by the LPB→POA pathway in cold defense. To address this issue in mice, we mapped the temperature sensitivity of the LPB→POA/DMH pathways via cFos staining. As expected, warm-activated neurons predominantly reside in the LPB→POA pathway. In contrast, cold-activated neurons are equally distributed between the two pathways (**Fig. 1e-h**). Blocking either of these two pathways impaired cold defense to a similar extent (**Fig. 3**), suggesting both pathways are required for physiological defense responses to cool/cold temperatures. The reason for not seeing a hyperthermia phenotype after bulk optogenetic activation of the LPB→POA pathway might be due to a masking effect of heat-defense neurons within this pathway^25, 26^. Thus, more selective genetic manipulations are needed to reveal the identity of cold-defense neurons within the LPB→POA pathway. Most notably, blocking both LPB→POA/DMH pathways exhibited a much stronger impairment in cold defense (**Fig. 3h-k**), suggesting a cumulative or additive effect of the two pathways. Thus, these two pathways assemble into a parallel circuit.

Interestingly, this LPB→DMH pathway may also function in rats since cold exposure also induces cFos expression in DMH-projecting LPB neurons in rats^39^. Hence, mice and rats may use an evolutionarily conserved neural circuit that we describe here, namely the parallel circuit comprised of the LPB→POA/DMH pathways, to control cold defense. The evolution of parallel neural circuits in cold defense not only enables resilience to hypothermia but also provides a scalable, robust, and efficient network in heat production when both pathways are recruited.

### Genetically defined LPB neurons relay thermal afferent signals to differentially control thermal effector activities

The LPB is considered the ‘relay station’ for transmitting warm and cold signals from the spinal cord to the POA. Recent findings in mice reveal that warm signals are relayed by different types of neurons to control different warm defense activities^25, 26^. For instance, we have shown that warm-activated Prodynorphin^+^ (Pdyn^+^) and cholecystokinin^+^ (CCK^+^) neurons in the LPB are responsible for inhibiting BAT thermogenesis and promoting skin vasodilation, respectively^25^. Here, we show that cold signals were relayed by a parallel circuit, namely the LPB→ POA/DMH pathways, to control a variety of cold defense activities (**Fig. 3** **and** **Fig. 4**). Among these activities, we showed that the LPB^SST^→DMH^LepR^ pathway governs iBAT thermogenesis, suggesting a genetically defined projection controls specific cold defense activities. We reasonably speculate that other cold defense activities, including heart rate and muscle shivering, are also controlled by genetically defined neural projections. Together, these findings suggest that the LPB is a critical thermoregulation center that relays thermal afferent signals via different genetically defined neurons or projections as part of a feed-forward mechanism to control different thermal effector activities.

Whether LPB neurons simply relay these afferent signals or actively process them remains an interesting question. Considering that the LPB receives diverse inputs from brain regions with functions related to internal state sensing and fear, including the nucleus of the solitary tract (NTS)^25, 50^, the rostral ventrolateral medulla^51^, and the paraventricular hypothalamus^52^, we propose that the LPB may also incorporate other types of signals to modify thermal afferent signals. In support of this hypothesis, it has recently been suggested that the LPBel receives input from the NTS and sends projections to the posterior subthalamic nucleus to mediate innate fear-associated hypothermia^50^.

## Acknowledgments

We thank Drs. Ji Hu, Zhenge Luo for sharing reagents; Dr. Xiaoming Li and the Molecular Imaging Core Facility (MICF) of School of Life Science and Technology, ShanghaiTech University for Microscopic imaging; Dr. Ying Xiong and the Molecular Cellular Core for slices and staining; all Shen lab members and “Shen Xian Hui (NPC)” Wechat group for valuable discussion. This study was funded by the National Key Research and Development Program of China (2019YFA0801900), National Nature Science Foundation of China (32122039 to W. Shen; 32100825 to W. Yang), Shanghai Science and Technology Committee of Shanghai City (19140903800 and 21XD1422700 to W. Shen), Shanghai Sailing Program (21YF1429800 to W. Yang), “Chen Guang” project supported by Shanghai Municipal Education Commission and Shanghai Education Development Foundation (20CG71 to W. Yang), Ministry of Science and Technology of China (2017YFA0205903), and Shanghai Frontiers Science Center for Biomacromolecules and Precision Medicine at ShanghaiTech University. We thank the staff members of the Animal Facility at the National Facility for Protein Science in Shanghai (NFPS), Zhangjiang Lab, China, for providing technical support and assistance.

## Author contributions

W.Z.Y., H.X., X.D. and Q.Z. performed most of the experiments; Y.X., Z.L. and Z.Z. performed behavioral evaluations; R.H., H.S., Q.S., Q.X. and X.N. performed the immunostaining; X.J. performed retro-TRAP analysis; J.X. and W.Z. performed the electrophysiology; X.F. designed and performed a concurrent recording of BAT and core temperatures; W.Z.Y. and H.T. designed the graphical abstract; W.Z.Y., R.Z., X.X., H.W., Y.F., L.W., X.L., H.Y., Q.Y. T.Y., Q.S., and W.L.S. designed the experiments; W.Z.Y., H.X., X.D., Q.Z. and W.L.S. wrote the manuscript.

## Data and materials availability

All data presented in the main text or supplementary materials are available from the corresponding author upon reasonable request.

## Code availability

No customized code was generated in this study.

## Reference

1. Braun, T. & Mosinger, B. Effect of hypothermia on death by starvation. Nature 181, 968 (1958).

2. Gasparrini, A., et al. Mortality risk attributable to high and low ambient temperature: a multicountry observational study. Lancet 386, 369–375 (2015).

3. Webb, P., et al. Hunger and malnutrition in the 21st century. Bmj 361, k2238 (2018).

4. Boyd, S.C. & Caul, W.F. Evidence of state dependent learning of brightness discrimination in hypothermic mice. Physiol Behav 23, 147–153 (1979).

5. Polderman, K.H. Mechanisms of action, physiological effects, and complications of hypothermia. Crit Care Med 37, S186–202 (2009).

6. Tan, C.L. & Knight, Z.A. Regulation of Body Temperature by the Nervous System. Neuron 98, 31–48 (2018).

7. Teague, R.S. & Ranson, S.W. The Role Of The Anterior Hypothalamus In Temperature Regulation. American Journal of Physiology-Legacy Content 117, 562–570 (1936).

8. Clark, G., Magoun, H.W. & Ranson, S.W. Hypothalamic Regulation Of Body Temperature. Journal of neurophysiology 2, 61–80 (1939).

9. H T Hammel, a. & Pierce, J.B. Regulation of Internal Body Temperature. Annual review of physiology 30, 641–710 (1968).

10. Morrison, S.F. & Nakamura, K. Central Mechanisms for Thermoregulation. Annual review of physiology 81, 285–308 (2019).

11. Song, K., et al. The TRPM2 channel is a hypothalamic heat sensor that limits fever and can drive hypothermia. Science 353, 1393–1398 (2016).

12. Tan, C.L., et al. Warm-Sensitive Neurons that Control Body Temperature. Cell 167, 47-+ (2016).

13. Yu, S.H., et al. Glutamatergic Preoptic Area Neurons That Express Leptin Receptors Drive Temperature-Dependent Body Weight Homeostasis. Journal of Neuroscience 36, 5034–5046 (2016).

14. Zhao, Z.D., et al. A hypothalamic circuit that controls body temperature. Proceedings of the National Academy of Sciences of the United States of America 114, 2042–2047 (2017).

15. Harding, E.C., et al. A Neuronal Hub Binding Sleep Initiation and Body Cooling in Response to a Warm External Stimulus. Current biology : CB 28, 2263–2273 e2264 (2018).

16. Kroeger, D., et al. Galanin neurons in the ventrolateral preoptic area promote sleep and heat loss in mice. Nature communications 9, 4129 (2018).

17. Padilla, S.L., Johnson, C.W., Barker, F.D., Patterson, M.A. & Palmiter, R.D. A Neural Circuit Underlying the Generation of Hot Flushes. Cell reports 24, 271–277 (2018).

18. Zhang, K.X., et al. Violet-light suppression of thermogenesis by opsin 5 hypothalamic neurons. Nature (2020).

19. Abbott, S.B.G. & Saper, C.B. Median preoptic glutamatergic neurons promote thermoregulatory heat loss and water consumption in mice. J Physiol 595, 6569–6583 (2017).

20. Moffitt, J.R., et al. Molecular, spatial, and functional single-cell profiling of the hypothalamic preoptic region. Science 362 (2018).

21. Krajewski-Hall, S.J., Miranda Dos Santos, F., McMullen, N.T., Blackmore, E.M. & Rance, N.E. Glutamatergic Neurokinin 3 receptor neurons in the median preoptic nucleus modulate heat-defense pathways in female mice. Endocrinology (2019).

22. Nakamura, K. & Morrison, S.F. A thermosensory pathway mediating heat-defense responses. Proceedings of the National Academy of Sciences of the United States of America 107, 8848–8853 (2010).

23. Nakamura, K. & Morrison, S.F. Preoptic mechanism for cold-defensive responses to skin cooling. The Journal of physiology 586, 2611–2620 (2008).

24. Geerling, J.C., et al. Genetic identity of thermosensory relay neurons in the lateral parabrachial nucleus. American journal of physiology. Regulatory, integrative and comparative physiology 310, R41–54 (2016).

25. Yang, W.Z., et al. Parabrachial neuron types categorically encode thermoregulation variables during heat defense. Sci Adv 6, eabb9414 (2020).

26. Norris, A.J., Shaker, J.R., Cone, A.L., Ndiokho, I.B. & Bruchas, M.R. Parabrachial opioidergic projections to preoptic hypothalamus mediate behavioral and physiological thermal defenses. eLife 10 (2021).

27. Machado, N.L.S., Bandaru, S.S., Abbott, S.B.G. & Saper, C.B. EP3R-Expressing Glutamatergic Preoptic Neurons Mediate Inflammatory Fever. The Journal of neuroscience : the official journal of the Society for Neuroscience 40, 2573–2588 (2020).

28. Pinol, R.A., et al. Preoptic BRS3 neurons increase body temperature and heart rate via multiple pathways. Cell metabolism (2021).

29. Tabarean, I.V. Activation of Preoptic Arginine Vasopressin Neurons Induces Hyperthermia in Male Mice. Endocrinology 162 (2021).

30. Saper, C.B. & Machado, N.L. The search for thermoregulatory neurons is heating up. Cell metabolism 33, 1269–1271 (2021).

31. Buijs, R.M., Swaab, D.F., Dogterom, J. & van Leeuwen, F.W. Intra- and extrahypothalamic vasopressin and oxytocin pathways in the rat. Cell Tissue Res 186, 423–433 (1978).

32. Almeida, M.C., Steiner, A.A., Branco, L.G. & Romanovsky, A.A. Neural substrate of cold-seeking behavior in endotoxin shock. PLoS One 1, e1 (2006).

33. da Conceicao, E.P.S., Morrison, S.F., Cano, G., Chiavetta, P. & Tupone, D. Median preoptic area neurons are required for the cooling and febrile activations of brown adipose tissue thermogenesis in rat. Scientific reports 10, 18072 (2020).

34. Houtz, J., Liao, G.Y., An, J.J. & Xu, B. Discrete TrkB-expressing neurons of the dorsomedial hypothalamus regulate feeding and thermogenesis. Proceedings of the National Academy of Sciences of the United States of America 118 (2021).

35. Pinol, R.A., et al. Brs3 neurons in the mouse dorsomedial hypothalamus regulate body temperature, energy expenditure, and heart rate, but not food intake. Nature neuroscience (2018).

36. Rezai-Zadeh, K., et al. Leptin receptor neurons in the dorsomedial hypothalamus are key regulators of energy expenditure and body weight, but not food intake. Mol Metab 3, 681–693 (2014).

37. Zhou, Q., et al. Cooling-activated dorsomedial hypothalamic BDNF neurons control cold defense in mice. Journal of neurochemistry (2022).

38. Jeong, J.H., et al. Cholinergic neurons in the dorsomedial hypothalamus regulate mouse brown adipose tissue metabolism. Mol Metab 4, 483–492 (2015).

39. Nakamura, K. & Morrison, S.F. A thermosensory pathway that controls body temperature. Nature neuroscience 11, 62–71 (2008).

40. Machado, N.L.S., et al. A Glutamatergic Hypothalamomedullary Circuit Mediates Thermogenesis, but Not Heat Conservation, during Stress-Induced Hyperthermia. Current biology : CB 28, 2291–2301 e2295 (2018).

41. Petreanu, L., Huber, D., Sobczyk, A. & Svoboda, K. Channelrhodopsin-2-assisted circuit mapping of long-range callosal projections. Nature neuroscience 10, 663–668 (2007).

42. Chen, T.W., et al. Ultrasensitive fluorescent proteins for imaging neuronal activity. Nature 499, 295–300 (2013).

43. Ran, C., Hoon, M.A. & Chen, X. The coding of cutaneous temperature in the spinal cord. Nature Neuroscience 19, 1201–1209 (2016).

44. Yarmolinsky, D.A., et al. Coding and Plasticity in the Mammalian Thermosensory System. Neuron 92, 1079–1092 (2016).

45. Fenno, L.E., et al. Targeting cells with single vectors using multiple-feature Boolean logic. Nature methods 11, 763–U116 (2014).

46. Zingg, B., et al. AAV-Mediated Anterograde Transsynaptic Tagging: Mapping Corticocollicular Input-Defined Neural Pathways for Defense Behaviors. Neuron (2016).

47. Abreu-Vieira, G., Xiao, C.Y., Gavrilova, O. & Reitman, M.L. Integration of body temperature into the analysis of energy expenditure in the mouse. Molecular Metabolism 4, 461–470 (2015).

48. Shang, C., et al. Divergent midbrain circuits orchestrate escape and freezing responses to looming stimuli in mice. Nature communications 9, 1232 (2018).

49. Peiris, A.N., Jaroudi, S. & Gavin, M. Hypothermia. Jama 319, 1290 (2018).

50. Liu, C., et al. Posterior subthalamic nucleus (PSTh) mediates innate fear-associated hypothermia in mice. Nature communications 12, 2648 (2021).

51. Krukoff, T.L., Vu, T., Harris, K.H., Aippersbach, S. & Jhamandas, J.H. Neurons in the rat medulla oblongata containing neuropeptide Y-, angiotensin II-, or galanin-like immunoreactivity project to the parabrachial nucleus. Neuroscience 47, 175–184 (1992).

52. Ryan, P.J., Ross, S.I., Campos, C.A., Derkach, V.A. & Palmiter, R.D. Oxytocin-receptor-expressing neurons in the parabrachial nucleus regulate fluid intake. Nature neuroscience 20, 1722–1733 (2017).

