## Supplementary Information for "A parabrachial-hypothalamic parallel circuit governs cold defense in mice"

#### Supplementary Information List

**STAR METHODS**

- KEY RESOURCES TABLE
- CONTACT FOR REAGENT AND RESOURCE SHARING
- EXPERIMENTAL MODEL AND SUBJECT DETAILS
- METHOD DETAILS
    - Stereotaxic brain injection
    - Immunohistochemistry
    - Triple-labeling of warm/cold-activated cFos, POA- and DMH-projecting $LPB^{Vglut2}$ neurons
    - Slice physiological recording
    - Calcium fiber photometry
    - Fiber photometry data analysis
    - Cold plate test
    - Temperature preference test
    - Metabolic measurement
    - Core body temperature, heart rate, physical activity, and iBAT temperature measurement
    - Heat production calculation
    - Electromyogram (EMG) recording
    - iBAT sympathetic nerve denervation
    - Quantitative (or real-time) PCR
    - Food intake, body weight, and body composition analysis
    - Cell-type specific retro-TRAP sequencing
    - TRAP-seq data analysis

**Extended Data Fig. 1-9**

**Extended Data Table 1:** Summary of TRAP-seq data

**Extended Data Table 2:** Summary of statistical analyses

#### STAR METHODS

##### KEY RESOURCES TABLE

| REAGENT or RESOURCE | SOURCE | IDENTIFIER |
|---|---|---|
| **Antibodies** | | |
| Chicken anti-GFP | Abcam | Cat# ab13970<br>RRID: AB_300798 |
| Guinea pig anti-cFos | Synaptic systems | Cat# 226004<br>RRID: AB_2619946 |
| Rabbit anti-cFos | Synaptic systems | Cat# 226003<br>RRID: AB_2231974 |
| Rat anti-RFP | Chromotek | Cat# 5F8<br>RRID: AB_2336064 |
| Rabbit anti-Somatostatin | ImmunoStar | Cat# 20067<br>RRID: AB_572264 |
| Rabbit anti-Glutamate | Sigma | Cat# G6642<br>RRID: AB_259946 |
| Rabbit anti-GABA | Sigma | Cat# A2052<br>RRID: AB_477652 |
| DyLight 488 conjugated goat anti-chicken | Invitrogen | Cat# SA5-10070<br>RRID: AB_2556650 |
| Alexa Fluor 594 conjugated goat anti-rat IgG | Invitrogen | Cat# A-11007<br>RRID: AB_141374 |
| Alexa Fluor 594 conjugated goat anti-Guinea pig IgG | Invitrogen | Cat#A-11076<br>RRID: AB_ 141930 |
| Alexa Fluor 594 conjugated goat anti-rabbit IgG | Jackson | Cat# 111-585-144<br>RRID: AB_2307325 |
| Alexa Fluor 488 conjugated goat anti-rabbit IgG | Jackson | Cat# 111-545-003<br>RRID: AB_2338046 |
| Alexa Fluor 647 conjugated goat anti-rabbit | Invitrogen | Cat# A21244<br>RRID: AB_141663 |
| anti-EGFP antibody | Monoclonal Antibody Core Facility at Memorial Sloan-Kettering cancer center | an equal mixture of clones 19C8 and 19F7 |
| **Chemicals, Peptides, and Recombinant Proteins** | | |
| DAPI Fluoromount-G mounting medium | SouthernBiotech | Cat# 0100-20 |
| TRIzol reagent | Sangon | #B511311-0100 |

| | | |
|---|---|---|
| 0.5% Blocking reagent | Roche | Cat# 11096176001 |
| Clozapine N-oxide (CNO) | ENZO | Cat# BML-NS105-0025 |
| CNQX | Sigma | Cat# C127 |
| DL-AP5 | Tocris | Cat# 0105 |
| TTX | Sigma | Cat# T8024 |
| 4-AP | Sigma | Cat# 275875 |
| **Critical Commercial Assays** | | |
| Pierce™ Recombinant Protein L, Biotinylated | ThermoFisher Scientific | Cat# 29997 |
| RNeasy Mini Kit | QIAGEN | Cat# 74104 |
| Qubit® 2.0 Fluorometer | Life Technologies, USA | |
| SIGMAFAST™ Fast Red TR/Naphthol AS-MX Tablets | Sigma | Cat# F4648 |
| Reverse Transcription Kit | Takara | #RR047A |
| SYBR Green PCR Kit | Takara | #RR820A |
| **Deposited Data** | | |
| **Experimental Models: Organisms/Strains** | | |
| Vgat-IRES-Cre mice | Jackson Laboratory (USA) | JAX: 028862 |
| Vgat-T2A-FlpO mice | Jackson Laboratory (USA) | JAX: 029591 |
| Vglut2-IRES-Cre mice | Jackson Laboratory (USA) | JAX: 028863 |
| SST-IRES-Cre mice | Jackson Laboratory (USA) | JAX: 028864 |
| LepR-Cre mice | Jackson Laboratory (USA) | JAX: 008320 |
| C57BL/6J mice | Silaike Experiment Animal Co., Ltd. (Shanghai) | JAX: 000664 |
| Ai14 mice | Jackson Laboratory (USA) | JAX: 007908 |
| ChAT-Cre mice | Jackson Laboratory (USA) | JAX: 006410 |
| **Oligonucleotides** | | |
| **Recombinant DNA** | | |
| AAV2-Retro-hEF1a-DIO-GFPL10-WPRE-hGHpA | Shanghai Taitool Bioscience Co. | N/A |
| AAV2-Retro-hSyn-DIO-mCherry-WPRE-pA | Shanghai Taitool Bioscience Co. | N/A |

| | | |
|---|---|---|
| AAV2/9-hSyn-DIO-GCaMP6s-WPRE-SV40pA | Shanghai Taitool Bioscience Co. | N/A |
| AAV2-Retro-hSyn-DIO-GCaMP6s-WPRE-SV40pA | Shanghai Taitool Bioscience Co. | N/A |
| AAV2/9-hSyn-DIO-jGCaMP7b-WPRE-pA | Shanghai Taitool Bioscience Co. | N/A |
| AAV2/9-hEF1a-DIO-hChR2(H134R)-mCherry | Shanghai Taitool Bioscience Co. | N/A |
| AAV2/9-hSyn-DIO-hM3Dq-EGFP | Shanghai Taitool Bioscience Co. | N/A |
| AAV2/9-hEF1a-DIO-hChR2(H134R)-EYFP | Shanghai Taitool Bioscience Co. | N/A |
| AAV2/9-hEF1a-FDIO-hChR2(H134R)-EYFP | Shanghai Taitool Bioscience Co. | N/A |
| AAV2/9-hEF1a-DIO-mCherry-P2A-TeNT-WPRE-pA | Shanghai Taitool Bioscience Co. | N/A |
| AAV2/8-hSyn-DIO-mCherry-WPRE-pA | Shanghai Taitool Bioscience Co. | N/A |
| AAV2/9-hEF1a-FDIO-taCasp3-TEVp | Shanghai Taitool Bioscience Co. | N/A |
| AAV2/8-hEF1a-FDIO-mCherry | Shanghai Taitool Bioscience Co. | N/A |
| scAAV2/1-hSyn-FlpO-pA | Shanghai Taitool Bioscience Co. | N/A |
| AAV2/8-hEF1a-FDIO-hM3Dq-mCherry | Shanghai Taitool Bioscience Co. | N/A |
| scAAV2/1-hSyn-Cre-pA | Shanghai Taitool Bioscience Co. | N/A |
| AAV2/9-hSyn-DIO-EGFP-WPRE-hGHpA | Shanghai Taitool Bioscience Co. | N/A |
| AAV2/5-hSyn-DIO-GFP-P2A-TeNT | Shanghai Taitool Bioscience Co. | N/A |
| AAV2/9-hEF1a-FDIO-mCherry-2A-TeNT-WPRE-pA | Shanghai Taitool Bioscience Co. | N/A |
| AAV2/9-hSyn-eGFP-2A-TeNT-WPRE-hGHpA | Shanghai Taitool Bioscience Co. | N/A |
| AAV2/9-hSyn-MCS-EGFP-3FLAG | Shanghai Taitool Bioscience Co. | N/A |
| AAV2-Retro-CAG-DIO-FlpO | Shanghai Taitool Bioscience Co. | N/A |
| pAAV2/5-CAG-DIO-GtACR1-P2A-EGFP | Shanghai Taitool Bioscience Co. | N/A |
| AAV2/9-EF1α-DIO-hM4D(Gi)-mCherry | BrainVTA (Wuhan) Co., Ltd | N/A |

| | | |
|---|---|---|
| AAV2/9-hSyn-Cre-off-FlpO-on-EGFP-T2A-TeNT | Shanghai Taitool Bioscience Co. | N/A |
| AAV2/9-hSyn-Cre-off-FlpO-on-EGFP | Shanghai Taitool Bioscience Co. | N/A |
| **Software and Algorithms** | | |
| Digbehv video-recording software | Jiliang, Shanghai, China | http://www.ecgonline.cn/main |
| Clampex 10 data acquisition software | Molecular Devices | https://www.moleculardevices.com/products/axon-patch-clamp-system/acquisition-and-analysis-software/pclamp-software-suite |
| EV Capture | Hunan Yiwei LTD, China | https://www.ieway.cn/evcapture |
| MATLAB | MathWorks | https://www.mathworks.com/ |
| ImageJ bundled with Java 1.8.0_172 | NIH ImageJ | https://imagej.nih.gov/ij |
| SPIKES2 | CED, Cambridge, UK | http://ced.co.uk/products/spkovin |
| VITALVIEW | Starr Life Sciences Corp., Oakmont, USA | https://www.starrlifesciences.com/ |
| CLAMS | Columbus, USA | http://www.colinst.com |
| GraphPad Prism 8 | GraphPad | https://www.graphpad.com/scientific-software/prism |
| Excel | Microsoft | https://www.microsoft.com/microsoft-365/excel |
| **Other** | | |
| Regular Chow Diet | SLAC | Cat# M03 |
| High Fat Diet | Research Diets | Cat# D12492 |

#### CONTACT FOR REAGENT AND RESOURCE SHARING

#### EXPERIMENTAL MODEL AND SUBJECT DETAILS

#### Animals

Animal care and use conformed to institutional guidelines of ShanghaiTech University, Shanghai Biomodel Organism Co., and governmental regulations. All experiments were performed on male adult mice (8–16 weeks old). Mice were housed under controlled temperature (22–25 °C) unless specified in a 12-hour reverse light/dark cycle (light time, 9 pm to 9 am) with food and water ad libitum. Mice were fed either a high fat diet (HFD, Research Diets, #D12492) or regular chow food as described in the text and figure legends. The mice strains were listed in the Key Resources Table.

#### METHOD DETAILS

##### Stereotaxic brain surgeries and viral injection

Mice were given general anesthesia with isoflurane during stereotaxic injection. The injection was performed using a small animal stereotaxic instrument (David Kopf Instruments, #PF-3983; RWD Life Science, #68030; Thinker Tech Nanjing Biotech, #SH01A). AAV virus (~0.15 μl, unless specified) was delivered through a pulled-glass pipette and a pressure micro-injector (Nanoject II, #3-000-205A, Drummond) at a slow rate (23 nl $min^{-1}$) with customized controllers. The coordinates of viral injection sites include the LPB (AP, -4.95 mm; ML, ± 1.5mm; DV, -3.6 mm), the DMH (AP, -1.25 mm; ML, ± 0.3 mm; DV, -5.2 mm), the VMPO (AP, 0.75 mm; ML, 0 mm; DV, -5.0 mm), the vLPO (AP, 0.35 mm; ML, ± 0.75 mm; DV, -5.5 mm) and the RPa (AP, -5.8 mm; ML, 0 mm; DV, -5.7 mm). The injection needle was withdrawn 10 min after the end of the injection. During surgeries, a feedback heater was used to maintain core body temperature at 36 ± 1°C. The optical fiber (200 μm in diameter, Inper Inc., China) was chronically implanted in the LPB (AP, -4.95 mm; ML, ± 1.5mm; DV, -3.4 mm), and the DMH (AP, -1.25 mm; ML, ± 0.3 mm; DV, -4.9 mm) and secured with dental cement (C&B Metabond®, Parkell, Japan). The bregma sites and anatomy are indicated directly on relevant figures. Mice were transferred to housing cages for 3-4 weeks before performing behavioral evaluations. After behavioral tests were finished, mice were perfused to check the virus expression. Data from mice (often 0-20%) with few or no viral expression were removed. We used a mouse brain atlas (Franklin and

Paxinos, 2008, 4th edition) to determine coordinates for injection sites. To construct the heatmap in **Extended Data Fig. 5**, we converted the photomicrograph of the injection site into a binary one, applied a Gaussian filter to remove the remaining noise, and stacked them together using ImageJ (version 1.8.0).

**Immunohistochemistry**

Mice were anesthetized with isoflurane and perfused transcardially with PBS and 4% paraformaldehyde (PFA) in PBS. Brain tissues were post-fixed overnight at 4°C, then sectioned at 50 μm using a vibratome (Leica, VT1200S). Brain slices were collected and blocked with blocking solution (Roche, #11096176001) for 2-h at room temperature and subsequently incubated with primary antibodies (1:1000, unless specified) for 2 days at 4°C. Then, the samples were washed three times in PBST (PBS with 0.1% Triton X-100, v/v) before incubating in secondary antibodies (1:1000, unless specified) overnight at 4°C. For glutamate, GABA, and somatostatin staining, mice were perfused transcardially with PBS followed by 50 ml 4% PFA without post-fixation. Brains were dehydrated in 20% sucrose for 1 day and 30% sucrose for 2 days at 4°C, then sectioned at 40 μm thicknesses on a cryostat microtome (Leica, CM3050s). Brain slices were collected and blocked with the blocking solution containing 10% normal goat serum (v/v) and 0.1% Triton X-100 (v/v) in PBS overnight at 4°C and subsequently incubated with primary antibodies (1:500) for 8-h at room temperature. After that, the slices were washed three times in PBST (PBS with 0.7% Triton X-100, v/v) before being incubated in secondary antibodies (1:1000) for 2-h at room temperature. The primary and secondary antibodies were listed in the Key Resources Table. Brain sections were washed three times in PBST and cover-slipped with DAPI Fluoromount-G mounting medium (SouthernBiotech, #0100-20). Images were captured on a Nikon A1R or Leica SP8 confocal microscope or Olympus VS120 Virtual Microscopy Slide Scanning System.

**Triple-labeling of warm/cold-activated cFos, POA- and DMH-projecting $LPB^{Vglut2}$ neurons**

Vglut2-IRES-Cre mice were injected with AAV-retro-DIO-GFPL10 into the POA and AAV-retro-DIO-mCherry into the DMH. Four weeks following the injection, mice were exposed to a warm (38°C) or cold (10°C) environment in an incubator for two hours, then were perfused with PBS and 4% PFA and processed following the immunohistochemistry protocol described above. POA-projecting $LPB^{Vglut2}$ neurons were labeled with GFP. DMH-projecting $LPB^{Vglut2}$ neurons were labeled with mCherry. Warm- or cold-activated cFos in the LPB neurons were labeled by cFos immunostaining (rabbit anti-cFos, Synaptic systems, #226003, 1:10000) colored with Fluor 647-conjugated secondary antibody (Alexa Fluor 647 conjugated goat anti-rabbit, Invitrogen, #A21244, 1:1000). Images were captured on a Nikon A1R confocal microscope.

**Slice physiological recording**

Brain slice recording and data analysis were performed similarly as described in[1, 2]. Briefly, slices containing the DMH regions were prepared from adult mice anesthetized before decapitation. Brains were removed and placed in ice-cold oxygenated (95% $O_2$ and 5% $CO_2$) cutting solution (228 mM sucrose, 11 mM glucose, 26 mM $NaHCO_3$, 1 mM $NaH_2PO_4$, 2.5 mM KCl, 7 mM $MgSO_4$, and 0.5 mM $CaCl_2$). Coronal brain slices (250 µm) were cut using a vibratome (VT 1200S, Leica Microsystems, Germany). The slices were incubated at 32°C in oxygenated artificial cerebrospinal fluid (ACSF: 119 mM NaCl, 2.5 mM KCl, 1 mM $NaH_2PO_4$, 1.3 mM $MgSO_4$, 26 mM $NaHCO_3$, 10 mM glucose, and 2.5 mM $CaCl_2$) for 1 h, and were then kept at room temperature under the same conditions before transfer to the recording chamber. The ACSF was perfused at 2 ml/min. The acute brain slices were visualized with a 40x Olympus water immersion lens, differential interference contrast optics (Olympus Inc., Japan), and a CCD camera (IR1000, Dage-MTI, USA).

Patch pipettes were pulled from borosilicate glass capillary tubes (#BF150-150-86-10, Sutter Instruments, USA) using a P-97 pipette puller (Sutter Instruments, USA). For EPSC recordings, pipettes were filled with solution (in mM: 130 $CsMeSO_3$, 1 $MgCl_2$, 1 $CaCl_2$, 10 HEPES, 2 QX- 314, 11 EGTA, 2 Mg-ATP, and 0.3 Na-GTP; pH 7.3; 295

mOsm). Cells were clamped at -70 mV. To block EPSCs, 10 μM CNQX (Sigma) and 50 μM DL-AP5 (Tocris) were applied to the bath solution. For IPSC recordings, cells were clamped at 0 mV. 10 μM CNQX (Sigma) and 50 μM DL-AP5 (Sigma) were applied to the bath solution. To block IPSCs, 25 μM bicuculline (Tocris) was applied to the bath solution. The resistance of pipettes varied between 3–5 MΩ. The signals were recorded with MultiClamp 700B, Digidata 1440A interface, and Clampex 10 data acquisition software (Molecular Devices). After the establishment of the whole-cell configuration, series resistance was measured. Recordings with series resistances of > 20 MΩ were rejected. Blue light pulses were delivered through the 40X objective of the microscope with the X-Cite LED light source. The light power density was adjusted to 7 mW/mm$^2$.

**Calcium fiber photometry**

Following injection of an AAV2/9-hSyn-DIO-GCaMP6s or AAV2-Retro-hSyn-DIO-GCaMP6s (Shanghai Taitool Bioscience Co.) viral vector, an optical fiber (200 μm O.D., 0.37 numerical aperture, Inper Inc., China) was placed 150 μm above the viral injection site. Post-surgery mice were transferred to housing cages for at least three weeks before any experiment. Fluorescence signals were captured with a dual-channel fiber photometry system (Fscope, Biolinkoptics, China) equipped with a 488-nm excitation laser (OBIS, Coherent), 505-544-nm emission filter, and a photomultiplier tube (Hamamatsu, #R3896). The gain (voltage) on PMT was set to 600 V. The laser power at the tip of the optical fiber was coordinated to 25 - 40 μW to minimize bleaching. The analog voltage signals were low-pass filtered at 30 Hz, digitalized at 100 Hz, and then acquired by Fscope software (Biolinkoptics, China).

We also used a fiber photometry system from Inper Ltd. to record the fluorescence signals from GCaMP6s. Light from a 470-nm LED was bandpass filtered, collimated, reflected by dichroic mirrors, focused by a 20× objective, and delivered at a power of 25 - 40 μW on the tip of the fiber optic cannula. Emitted fluorescence from GCaMP6s was bandpass filtered and focused on the sensor of a CMOS camera. The end of the fiber was imaged at a frame rate of 60 fps with the Inper Studio, and the mean value of the ROI of an end-face of the fiber was calculated using Inper Analysis software. To serve as an isosbestic control channel, 410-nm LED light was delivered alternately with 470-nm LED light.

To sterilize the surface, the optical fiber and the distal ends of the cord were cleaned using 75% ethanol. All animals were allowed to acclimate for at least 30 min after the fiber cord attachment. To control cage floor temperature changes, we used a Peltier controller (#5R7-001, Oven Industry) with a customized Labview code (National Instrument) to control Peltier floor plates (15 x 15 cm) for each mouse.

**Fiber photometry data analysis**

To calculate the fluorescence change ratios, we analyzed the raw data using Fscope software or Inper Analysis software with customized MATLAB code. We segmented the data based on behavioral events within individual trials. The values of fluorescence change ($\Delta F/F_0$) were derived by calculating $(F-F_0)/F_0$, where $F_0$ is the baseline fluorescence signal averaged in a 120 s time window prior to temperature changes. The peak $\Delta F/F_0$ in (**Fig. 2e**) represents the maximum $\Delta F/F_0$ after warming or cooling the floor. The cooling phase of $T_{floor}$ in (**Fig. 2f**) is defined as the period that $T_{floor}$ changes from the starting temperature to a target temperature, and the steady phase is defined as the period that $T_{floor}$ maintains at the target temperature. Finally, we plotted the average fluorescence changes ($\Delta F/F_0$) with different events using Customized MATLAB code and GraphPad Prism 8 (GraphPad).

**Cold Plate Test**

For quantification of spontaneous pain, the cold plate test was performed by using the test chamber with a homemade thermostatic plate, which is able to reach variable temperatures. Mice were placed on the plate fixed at 4 °C, free to move and walk. Spontaneous nociceptive behavior was quantified by counting the number of paw lifts recorded in a trial of 5 minutes. We also measured the limb withdrawal latency, which defines the latency of the mice's first limb lift after the mice had been placed on the cold plate.

**Temperature preference test**

Mice were individually housed at least seven days before testing. Mice with the blocking of LPB-innervating POA/DMH neurons (**Extended Data Fig. 5j**) were placed in a chamber containing two identical adjacent plates with one set to 30 ± 0.5 °C and the other adjusted to various temperatures as indicated (30, 16, 10, 6, and 35 °C). Mice were free to explore for five minutes, and each test was videotaped for later analysis. The mice moving trajectories were analyzed by Digbehv video-recording software (Jiliang, Shanghai, China) to calculate the total time spent on each side. All tests were performed in the dark phase between 9 am and 6 pm.

**Metabolic measurement**

For DREADDs and TeNT mice, energy expenditure, locomotor activity, and core body temperature were monitored by the Comprehensive Lab Animal Monitoring System with Temperature Telemetry Transmitter (CLAMS; Columbus Instruments, with G2 E-Mitter transponders) and ambient temperature was labeled in related figures. The data was acquired at a 10-min interval, as shown in the figures. Temperature transponders were implanted into the peritoneal cavity 3 – 5 days before testing. Mice were adapted in the chambers for 2 days before giving saline (volume (μl) = 10 x body weight (grams)), and CNO (ENZO, #BML-NS105-0025, I.P., 2.5 mg/kg body weight). Stimuli (drugs or temperature) were delivered in the dark phase.

**Core body temperature, heart rate, physical activity, and iBAT temperature measurement**

The core temperature, heart rate, and physical activity were recorded at a 1-min interval by VitalView Data Acquisition System Series 4000 (Starr Life Sciences Corp., Oakmont, USA) unless specified. To record core temperature and physical activity, we used G2 E-Mitter transponders implanted intraperitoneally. To record core temperature, heart rate, and physical activity, we used the G2-HR E-Mitter transponders. The HR E-Mitter was slipped into the abdominal cavity along the sagittal plane, and both leads were brought out of the abdominal incision. We then secured the negative lead (black) against a chest muscle on the animal's right. We secured the positive lead (shorter - clear) against a chest muscle on the animal's left. After recovery, mice were placed in their homecages on top of the ER-4000 Receivers. The heart rate is then recorded by the VitalView system as a beats-per-minute value based on computation from the R-R (R wave of the QRS complex) interval. Temperature challenges, laser stimulations and drugs were delivered in the dark phase.

The interscapular brown adipose tissue (iBAT) temperature ($T_{iBAT}$) was measured using a thermal infrared camera (A655sc, FLIR). And we analyzed the infrared images using the FLIR Tools software (Teledyne FLIR). To measure the iBAT temperature, the hair on top of the iBAT was shaved 3-5 days before measurement. The $T_{iBAT}$ was evaluated by measuring the average temperature in the interscapular region. We used the factory settings to convert raw pixel counts to units of temperature and analyzed the pictures taken from the same angle.

For wired recording with high temporal resolution shown in **Fig. 5a**,**b**, we applied a modified thermocouple unit as we reported before[1]. Two T-type thermocouples (TT-40, Omega) were implanted in the abdomen and within the iBAT to measure the core and iBAT temperatures, respectively. Mice were recovered for 3 - 5 days after surgery. To ensure these probes were accurate, we calibrated them using water baths and a standard thermal probe. The NI data acquisition card (USB-TC01, National Instrument) and supporting software were used for data collection, with a 1-Hz sampling rate.

**Heat production calculation**

The normalized heat production of mice after photoactivation of $LPB^{Vglut2\ \&\ ChR2}$ terminals in the DMH under different ambient temperatures, as shown in **Fig. 4r**. The heat production was calculated according to the paper[3]. The original equation is: heat loss rate (HLS) = $Cp_{mouse}*(T_{core2}-T_{core1})/(t_2-t_1)/[(T_{core1}+T_{core2})/2-(T_{a1}+T_{a2})/2]$, which $Cp_{mouse}$ represents the heat production capacity; in other form: $Cp_{mouse}$ = $[(T_{core1}+T_{core2})/2-(T_{a1}+T_{a2})/2]$ * HLS * $(t_2-t_1)$ / $(T_{core2}-T_{core1})$. While in our situation: $t_2$ = 30 min, $t_1$ = 0 min, and $T_{a1} = T_{a2}$, thus, the equation can be simplified as $Cp_{mouse}$ = 30 *

HLS * (($T_{core\_mean}$ -$T_a$) / $\Delta T_{core}$), and the parameter – HLS relatively constant; so, the heat production capacity somehow correlated with the rate of ($T_{core\_mean}$-$T_a$) / $\Delta T_{core}$. Then the heat production of each $T_a$ was normalized by dividing the mean value of the heat production of $T_a$ = 30 °C.

**Electromyogram (EMG) recording**

Mice were anesthetized and fixed with the animal stereotaxic instrument to avoid movement. An aluminum thermal pad with circulating water was used to maintain mice body temperature at 36 ± 1.5 °C. The rectal temperature was monitored by a thermocouple to reflect the body temperature under anesthesia. Another thermocouple to record skin temperature was taped onto the abdominal skin. Temperatures were recorded by NI data acquisition card and supporting software (National Instrument). Handmade electrodes for EMG recording were inserted into nuchal muscles after scissoring the skin, and the signal was amplified (×1000) and filtered (10 - 1000 Hz) with an 1800 2-Channel Microelectrode AC Amplifier (A-M Systems, Sequim, WA, USA). The laser stimulation pattern was 6 mW, 10 Hz, 10 ms on, for 60 s or 120 s as shown in the figures. The EMG amplitude was quantified (Spike 2, CED, Cambridge, UK) in sequential 4-sec bins as the square root of the total power (root mean square) in the 0 – 500 Hz band of the auto spectra of each 4-s segment [4].

**iBAT sympathetic nerve denervation**

Mice were anesthetized with isoflurane. The hair in the targeted area was shaved, and the area was sterilized with 95% ethanol-soaked sterile gauze. A midline incision was made in the upper dorsal skin to expose both iBAT pads, and styptic powder (KELC, USA) was applied to the area to prevent bleeding. We gently exposed the medial, ventral surface of both pads to visualize nerves beneath the pads. All five nerves innervating both pads were cut off a length of 2-3 mm to avoid nerve regeneration under a stereoscope. Mice were housed in the thermoneutral environment for 5-7 days before tests.

**Quantitative (or real-time) PCR**

RNA was extracted using TRIzol reagent (Sangon, #B511311-0100). The RNA quality and quantity were determined using a NanoDrop 5500 (Thermo). mRNA was reverse transcribed using a High-Capacity cDNA Reverse Transcription Kit (Takara, #RR047A) according to the manufacturers' instructions and processed for quantitative real-time PCR using the SYBR Green PCR system (Takara, #RR820A). The primers used to amplify a fragment of Ucp1 were 5'-ACTGCCACACCTCCAGTCATT-3' and 5'-CTTTGCCTCACTCAGGATTGG-3'. Mouse β-actin was used as the endogenous control to which sample values were normalized. The sequences used to amplify a fragment of β-actin were 5'- GTGACGTTGACATCCGTAAAGA -3' and 5'-GCCGGACTCATCGTACTCC -3'. Expression levels were calculated based on the $2^{-\Delta\Delta Ct}$ method.

**Food intake, body weight, and body composition analysis**

For the optogenetic experiments shown in **Fig. 5d-j**, the body weight and food intake were measured in the home cage. For DIO mice, the 5-week-old mice were fed with an ad libitum high-fat diet (HFD, Research Diets, #D12492) to drive body weight gain. AAVs carrying Cre-dependent GFP control or ChR2-tdTomato were injected in the DMH of Vglut2-IRES-Cre mice ten weeks after HFD feeding. Mice's body weight and food intake were measured daily (~ 4 pm). Four weeks after viral injections, mice were photo-stimulated to activate the $LPB^{Vglut2}$ terminals in the DMH for two weeks, as indicated in the figure.

After two-week photoactivation of the $LPB^{Vglut2}\rightarrow$DMH projections, the body composition of both two groups of mice was assessed with a minispec whole-body composition analyzer (Burker Minispec CMR LF50). And the fat and lean mass were normalized to total body mass. On the final day, the iBAT (interscapular brown adipose tissue) and iWAT (inguinal white adipose tissue) were collected, weighed, and frozen in liquid nitrogen from all mice after activation of the $LPB^{Vglut2}\rightarrow$DMH projections and then perfused transcardially to collect brain tissues.

#### Cell-type specific retro-TRAP sequencing

The procedures were detailed in our previous reports[1]. Briefly, a recombinant Cre-dependent AAV plasmid expressing GFP-tagged ribosomal subunit L10a (AAV2-EF1a-DIO-EGFP-L10a) was gifted by Dr. Jeffrey Friedman and then packaged using rAAV2-retro capsids by Taitool. 300 nl of the viral aliquot was unilaterally injected into the DMH of Vglut2-IRES-Cre mice. Animals were sacrificed for immunoprecipitation four weeks after injection.

For TRAP experiments, brain slices (300 μm) containing the LPB regions were prepared from virus-injected mice and placed in ice-cold DEPC-PBS (VT 1200S, Leica Microsystems, Germany). Then, the LPB regions were cut out with microsurgical forceps. Tissue from 6 brains was pooled for each experimental repeat, and three experimental repetitions were performed. Ribosomes were immunoprecipitated using anti-EGFP antibodies that were conjugated to Protein L-coated magnetic beads (ThermoFisher Scientific) for 16-h at 4°C with rotating. An aliquot of input RNAs taken before immunoprecipitation and total immunoprecipitated RNAs were then purified and enriched with PCR to create the final cDNA library. Purified libraries were quantified and validated to confirm the insert size and calculate the mole concentration. The library construction and sequencing were performed by Shanghai Sinotech Genomics Corporation (China).

#### TRAP-seq data analysis

The RNA quantification data after sequencing were analyzed as the following. To determine the statistical significance and fold enrichment, we divided the RNAs from the IP by those from the input for each gene (IP/Input). Moreover, the hits were narrowed down by only analyzing the PB-enriched gene list downloaded from Allen Institute (https://alleninstitute.org/; Tool one: MOUSE BRAIN CONNECTIVITY-source search-filter source structure: Parabrachial nucleus; Tool two: MOUSE BRAIN-Fine Structure Search: Parabrachial nucleus). The top candidate genes were then selected for further analysis.

#### Data availability

The data that support the findings of this study are available from the corresponding author upon reasonable request.

#### Code availability

The code that supports the findings of this study is available from the corresponding author upon reasonable request.

#### Methods References

1. Yang, W.Z.*, et al.* Parabrachial neuron types categorically encode thermoregulation variables during heat defense. *Sci Adv* **6**, eabb9414 (2020).
2. Zhao, Z.D.*, et al.* A hypothalamic circuit that controls body temperature. *Proceedings of the National Academy of Sciences of the United States of America* **114**, 2042-2047 (2017).
3. Abreu-Vieira, G., Xiao, C., Gavrilova, O. & Reitman, M. Integration of body temperature into the analysis of energy expenditure in the mouse. *Molecular metabolism* **4** (2015).
4. Nakamura, K. & Morrison, S.F. Central efferent pathways for cold-defensive and febrile shivering. *The Journal of physiology* **589**, 3641-3658 (2011).

**Extended Data Fig. 1 | Projections of LPB glutamatergic neurons throughout the brain.**

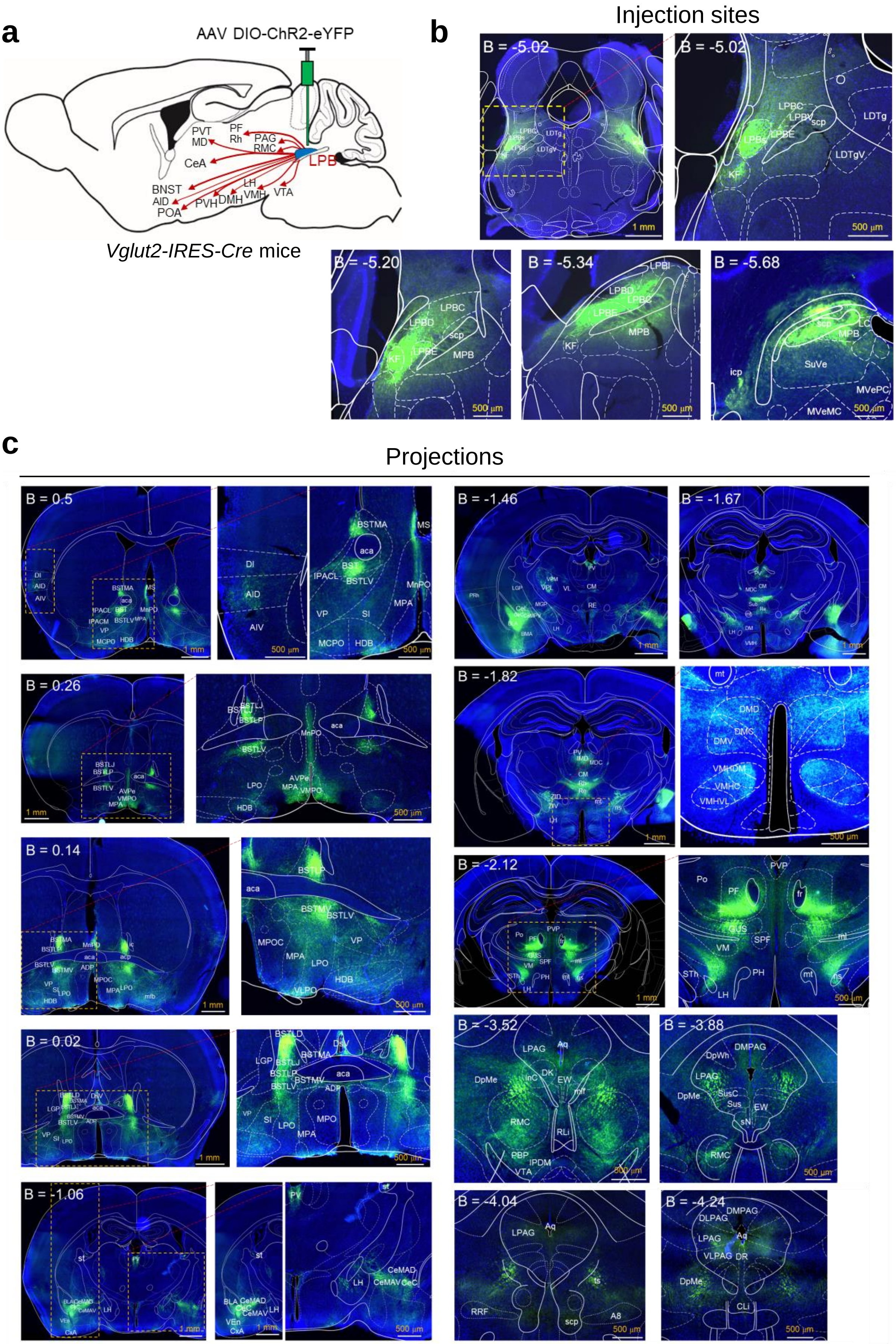

**Extended Data Fig. 1 | Projections of LPB glutamatergic neurons throughout the brain.**

(a) The axonal projection pattern of LPB glutamatergic neurons throughout the brain, revealed by anterograde tracing AAVs. Injection sites of AAV9-DIO-ChR2-eYFP in the LPB are indicated in the figure. Thicker lines indicate stronger eYFP expression.

(b) Representative images are showing ChR2-eYFP expression in the injection sites.

(c) Representative images are showing ChR2-eYFP expression in axonal terminals at various brain sites.

B, bregma; LPBC, lateral parabrachial nucleus, central part; LPBD, lateral parabrachial nucleus, dorsal part; LPBE, lateral parabrachial nucleus, external part; LPBV, lateral parabrachial nucleus, ventral part; LPBI, lateral parabrachial nucleus, internal part; LDTg, laterodorsal tegmental nucleus; LDTgV, laterodorsal tegmental nucleus, ventral part; KF, Kolliker-Fuse nucleus; MPB, medial parabrachial nucleus; LC, locus coeruleus; MVeMC, medial vestibular nucleus, magnocellular part; MVePC, medial vestibular nucleus, parvicellular part; icp, inferior cerebellar peduncle; scp, superior cerebellar peduncle; SuVe, superior vestibular nucleus. PAG, periaqueductal gray matter; SC, superior colliculus; RVM, rostral ventromedial medulla; PnO, the oral part of pontine reticular nucleus; TH, thalamus; Po, posterior thalamic nuclear group; POA, preoptic nucleus; PaF, parafascicular thalamic nucleus; SPF, subparafascicular thalamic nucleus; VPM, ventral posteromedial thalamic; ZID, zona incerta, dorsal part; DLPAG, dorsolateral periaqueductal gray; DMPAG, dorsomedial periaqueductal gray; LPAG, lateral periaqueductal gray; InG, intermediate gray layer of the superior colliculus; InWh, intermediate white layer of the superior colliculus; DpG, deep gray layer of the superior colliculus; Gi, gigantocellular reticular nucleus; GiA, gigantocellular reticular nucleus, alpha part; LPGi, lateral paragigantocellular nucleus; rmg, raphe magnus nucleus; LPO, lateral preoptic area; StHY, striohypothalamic nucleus; MPA, medial preoptic area; MPOM, medial preoptic nucleus, medial part; SIB, substantia innominata, basal part; HDB, nucleus of the horizontal limb of the diagonal band; D3V, dorsal 3rd ventricle; PVT, paraventricular thalamic nucleus.

### Extended Data Fig. 2 | Mapping the collateral projections of LPB$^{Vglut2}$ neurons to the POA (MnPO & VMPO) and the DMH.

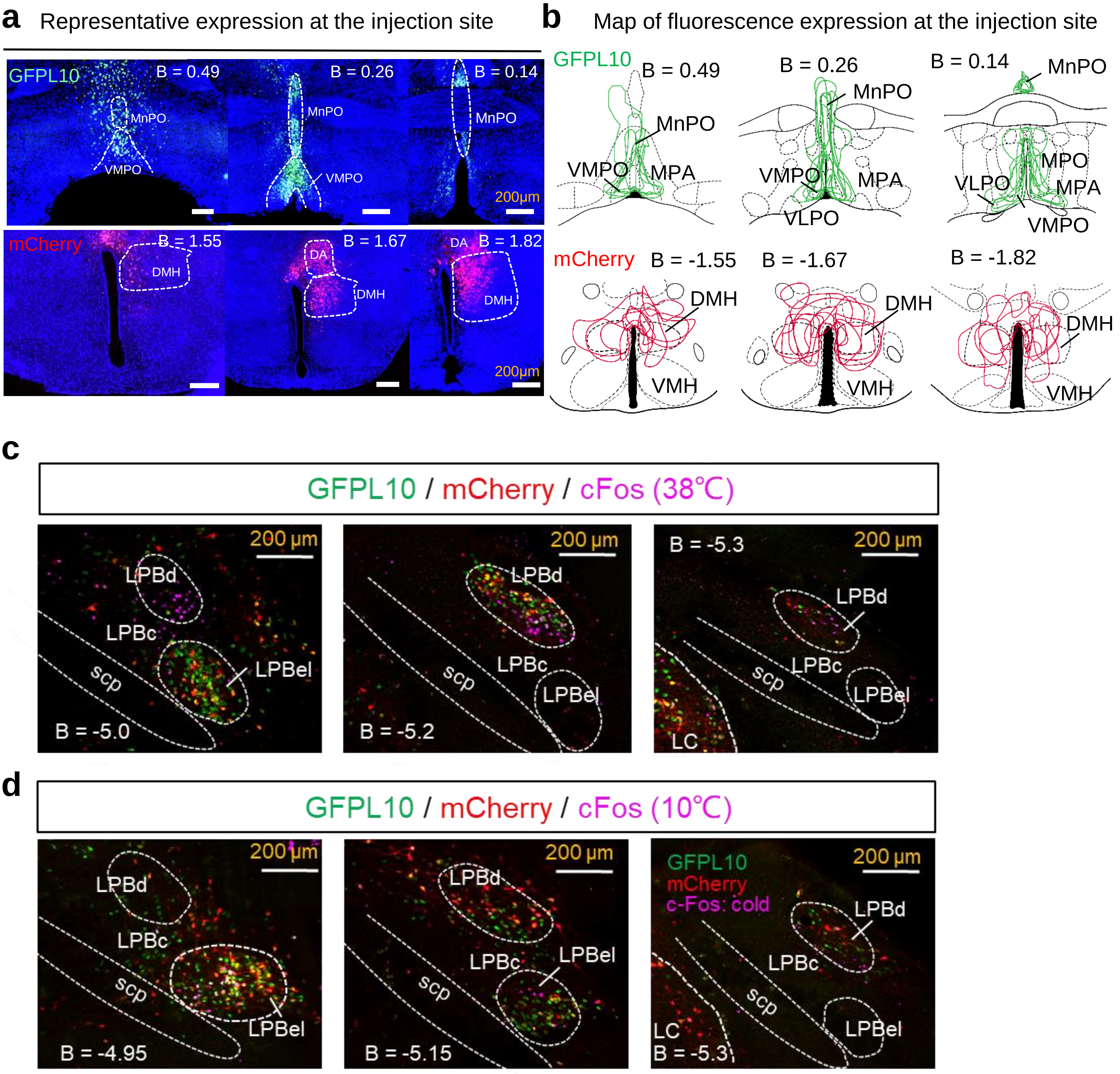

**Extended Data Fig. 2 | Mapping the collateral projections of LPB$^{Vglut2}$ neurons to the POA (MnPO & VMPO) and the DMH.**

(a) Representative expression of GFP (AAV-Retro-DIO-GFPL10) and mCherry (AAV-Retro-DIO-mCherry) in the injection sites. AAV-Retro-DIO-GFPL10 was injected in the VMPO & MnPO of Vglut2-IRES-Cre mice (n = 6 mice) and AAV-Retro-DIO-mCherry was injected in the DMH of Vglut2-IRES-Cre mice (n = 6 mice). The expression boundaries of each mouse were overlaid in the right as indicated.

(b) Map of fluorescence expression at the injection sites from (a). The expression boundaries of each mouse were overlaid and showed together as indicated (n = 6 mice).

(c) Representative images showing the overlapping between POA- and DMH-projecting LPB$^{Vglut2}$ neurons and warm-activated cFos at different bregma sites. Mice were exposed to warm stimuli (38$^{o}$C, 2 hrs) before sacrificing for cFos immunostaining. The overlapping ratios were indicated in Fig. 1e. A total of 924 cFos$^{+}$ neurons from 3 mice were scored.

(d) Representative images showing the overlapping between POA/DMH-projecting LPB neurons and cold-activated cFos immunoactivity at different bregma sites. Mice were exposed to cold stimuli (10$^{o}$C, 2 hrs) before sacrificing for cFos immunostaining. The overlapping ratios were indicated in Fig. 1g. A total of 693 cFos$^{+}$ neurons from 3 mice were scored.

B, bregma; LPBc, lateral parabrachial nucleus, central part; LPBd, lateral parabrachial nucleus, dorsal part; LPBel, lateral parabrachial nucleus, external and internal part; LC, locus coeruleus; scp, superior cerebellar peduncle; DMH, dorsomedial hypothalamus nucleus; DA, dorsal hypothalamus area VMH, ventromedial hypothalamus nucleus; MnPO, median preoptic area; VLPO, ventrolateral preoptic area; MPA, medial preoptic area; MPO, medial preoptic nucleus; VMPO, ventromedial preoptic area.

**Extended Data Fig. 3 | Projection pattern of DMH-projecting LPB$^{Vglut2}$ neurons throughout the brain.**

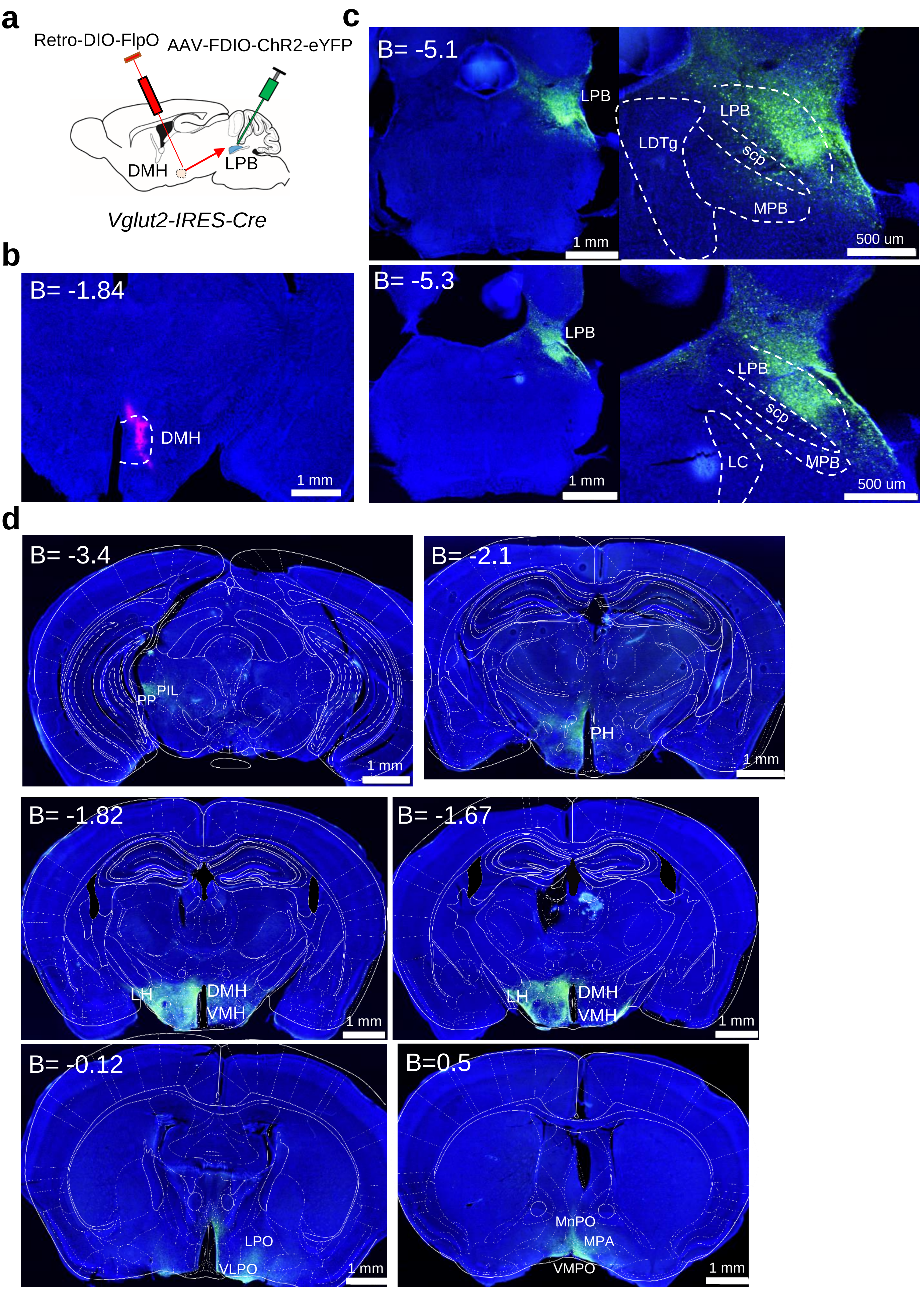

**Extended Data Fig. 3 | Projection pattern of DMH-projecting LPB$^{Vglut2}$ neurons throughout the brain.**

(a) Scheme for mapping the axonal projections of DMH-projecting LPB$^{Vglut2}$ neurons. For labeling DMH-projecting LPB$^{Vglut2}$ neurons, retrograde AAVs carrying Cre-dependent FlpO were injected in the DMH, which drove the expression of FlpO-dependent ChR2-eYFP in the LPB. A red tracer (CTB647) was co-injected into the DMH to indicate the injection sites.

(b) Representative image showing the injection sites in the DMH viewed by red tracer (CTB647).

(c) Representative images showing ChR2-eYFP expression in DMH-projecting LPB$^{Vglut2}$ neurons in the LPB.

(d) Representative images are showing ChR2-eYFP expression in axonal terminals at various brain sites.

B, bregma; LPB, lateral parabrachial nucleus; LDTg, laterodorsal tegmental nucleus; MPB, medial parabrachial nucleus; LC, locus coeruleus; scp, superior cerebellar peduncle; POA, preoptic nucleus; DMH, dorsomedial hypothalamus; VMH, ventromedial hypothalamus; LH, lateral hypothalamus; PH, poster hypothalamus; LPO, lateral preoptic area; MnPO, median preoptic nucleus; VMPO, ventromedial preoptic nucleus; MPA, medial preoptic area; VLPO, ventrolateral preoptic nucleus; PP, peripeduncular nucleus; PIL, posterior intralaminar thalamic nucleus.

### Extended Data Fig. 4 | Phenotypes associated with blocking of DMH$^{only}$-projecting or POA$^{only}$-projecting LPB neurons.

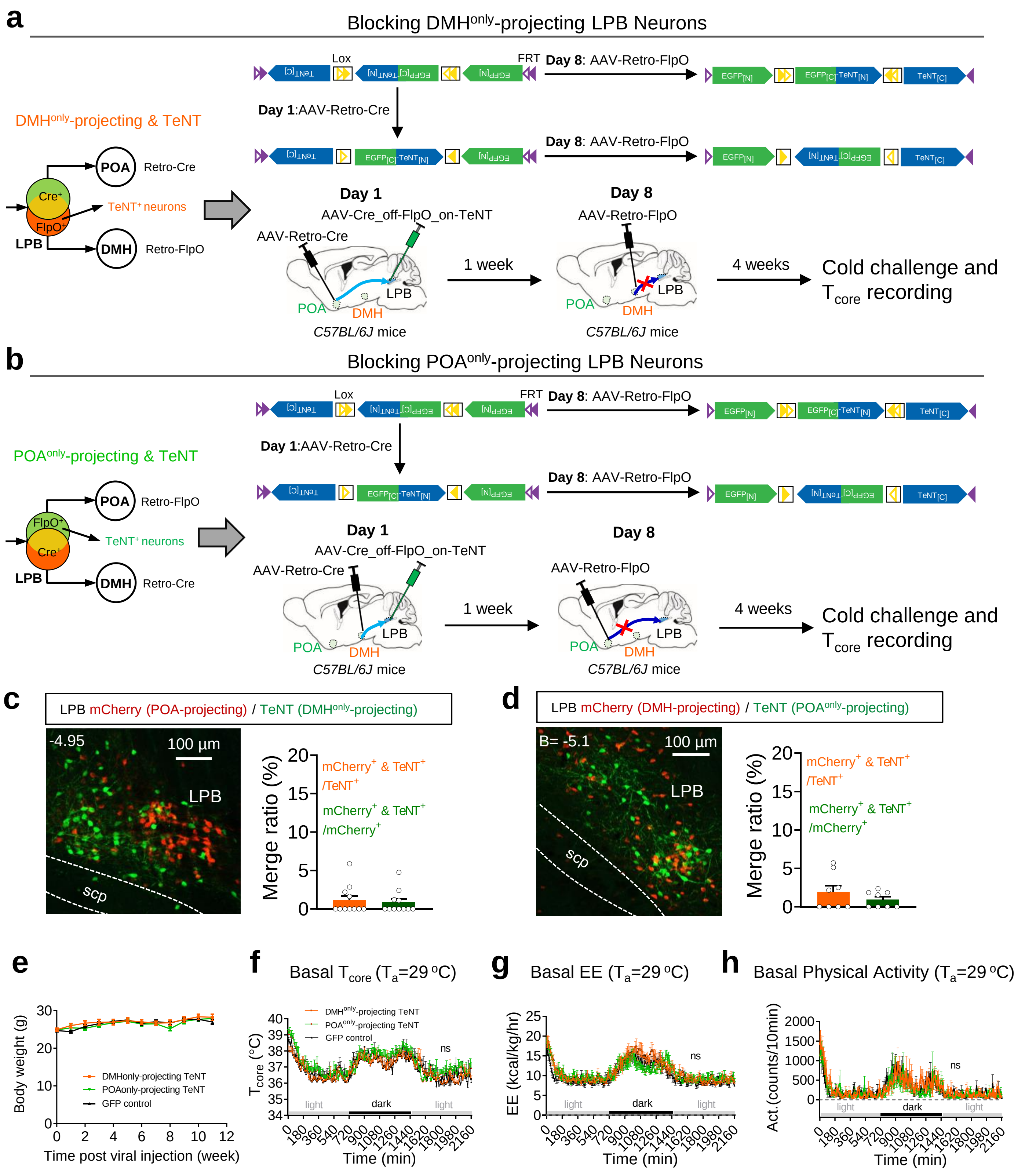

**Extended Data Fig. 4 | Phenotypes associated with blocking of DMH$^{only}$-projecting or POA$^{only}$-projecting LPB neurons.**

(a) The scheme to block LPB neurons that projected to the DMH but did not collaterally project to the POA (DMH$^{only}$-projecting LPB neurons). On day 1, the retrograde traveling AAV-Retro-hSyn-Cre virus was injected into the POA (MnPO & VMPO) and Cre_off-FlpO_on-TeNT (AAV9-hSyn-Cre_off-FlpO_on-EGFP-T2A-TeNT) injected in the LPB of C57BL/6J mice. Thus, AAV-Retro-Cre would travel to the LPB to turn off TeNT in turning off the TeNT expression in POA-projecting LPB neurons, thus unblocking POA-projecting LPB neurons. On day 8, AAV-Retro-hSyn-FlpO virus was injected into the DMH, which then traveled to the LPB to turn on the TeNT expression in DMH-projecting but not co-projecting LPB neurons (DMH$^{only}$-projecting LPB neurons). Cre_off-FlpO_on-EGFP (AAV9-hSyn-Cre_off-FlpO_on-EGFP) was used as the control for Cre_off-FlpO_on-TeNT.

(b) The scheme to block LPB neurons that projected to the POA but did not collaterally project to the DMH (POA$^{only}$-projecting LPB neurons). On day 1, the retrograde traveling AAV-Retro-hSyn-Cre virus was injected into the DMH and Cre_off-FlpO_on-TeNT (AAV9-hSyn-Cre_off-FlpO_on-EGFP-T2A-TeNT) injected in the LPB of C57BL/6J mice. Thus, AAV-Retro-Cre would travel to the LPB to turn off TeNT in turning off the TeNT expression in DMH-projecting LPB neurons, thus unblocking DMH-projecting LPB neurons. On day 8, AAV-Retro-hSyn-FlpO virus was injected into the POA (MnPO & VMPO), which then traveled to the LPB to turn on the TeNT expression in POA-projecting but not co-projecting LPB neurons (POA$^{only}$-projecting LPB neurons). The same control was used as in (a).

(c) Representative images showing the little overlapping between POA-projecting LPB neurons (mCherry$^{+}$) and DMH$^{only}$-projecting LPB neurons (TeNT$^{+}$). To label the POA-projecting LPB neurons, AAV-hSyn-DIO-mCherry was co-injected with Coff/Fon-TeNT-EGFP in the LPB in (a) so that AAV-Retro-Cre injected in the POA would travel to the LPB to turn on mCherry expression. Scale bar, 100 µm. The overlapping ratios were quantified in the right panel (11 brain slices from 3 mice).

(d) Representative images showing the little overlapping between DMH-projecting LPB neurons (mCherry$^{+}$) and POA$^{only}$-projecting LPB neurons (TeNT$^{+}$). To label the DMH-projecting LPB neurons, AAV-hSyn-DIO-mCherry was co-injected with Coff/Fon-TeNT-EGFP in the LPB in (b) so that AAV-Retro-Cre injected in the DMH would travel to the LPB to turn on mCherry expression. Scale bar, 100 µm. The overlapping ratios were quantified in the right panel (8 brain slices from 3 mice).

(e-h) Changes of body weight (e), basal $T_{core}$ (f), basal energy expenditure (g) and basal physical activity (h) at thermoneutral temperature (f-h) after blocking the DMH$^{only}$-projecting LPB neurons or POA$^{only}$-projecting LPB neurons, as indicated (n = 6 mice each).

LPB, lateral parabrachial nucleus; POA, preoptic area; DMH, dorsomedial hypothalamic nucleus; TeNT, Tetanus toxin. All data are shown as mean ± sem. The p-values are calculated based on two-way ANOVA (f-h). ns, not significant.

### Extended Data Fig. 5 | Heatmaps of TeNT expression and baseline measurement after blocking LPB-innervating POA or DMH neurons.

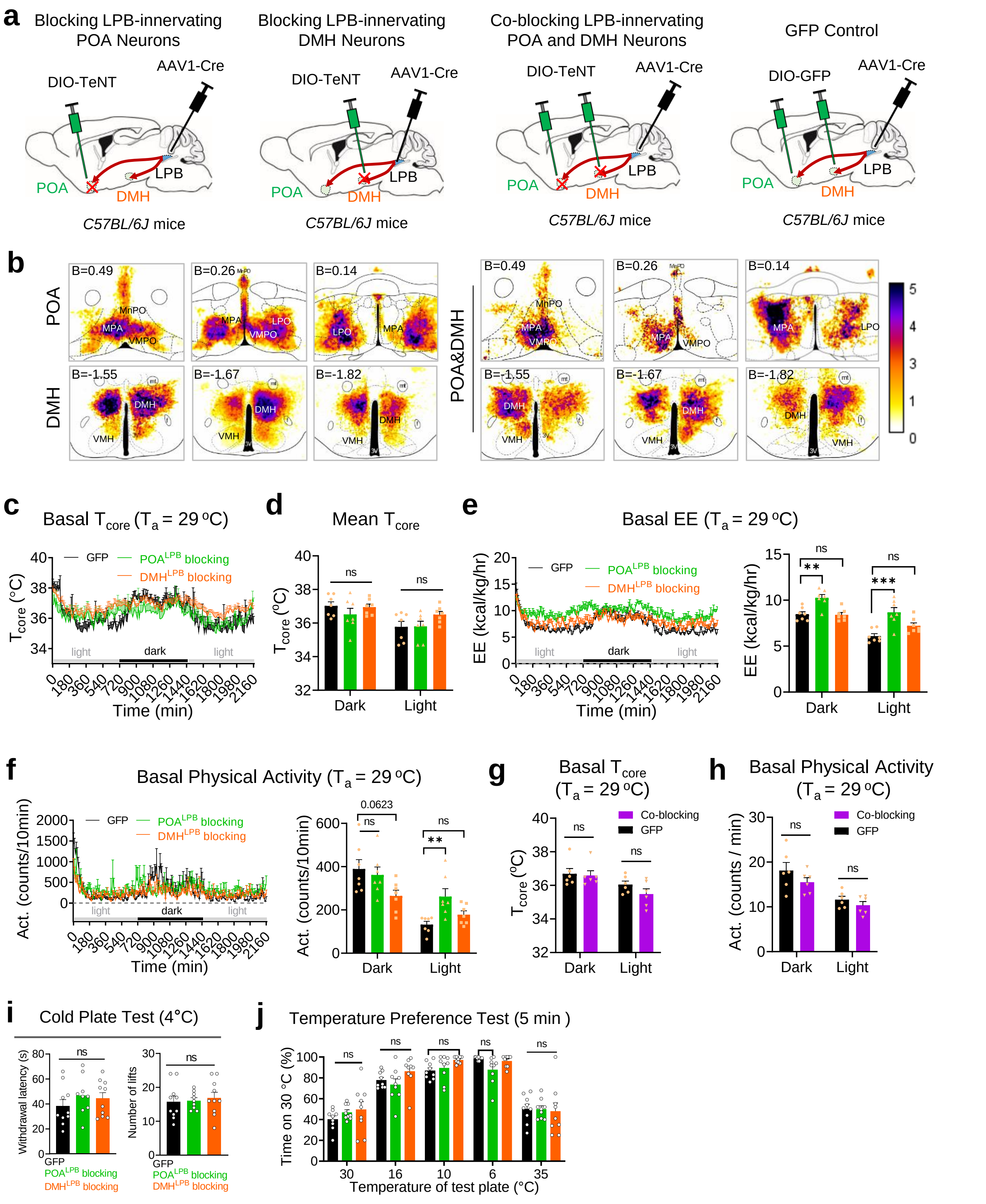

**Extended Data Fig. 5 | Heatmaps of TeNT expression and baseline measurement after blocking LPB-innervating POA or DMH neurons.**

(a) The scheme to block LPB-innervating POA (mainly MnPO, VMPO, MPA, and LPO) neurons, LPB-innervating DMH neurons, LPB-innervating POA and DMH Neurons (Co-blocking), and GFP control. The anterograde transsynaptic Cre carried by AAV1 (AAV1-hSyn-Cre) was injected in the LPB, which drove the expression of AAV9-hEF1a-DIO-mCherry-P2A-TeNT injected in the POA ($POA^{LPB}$ blocking), the DMH ($DMH^{LPB}$ blocking), or both (co-blocking), as indicated. AAV9-hSyn-DIO-GFP was injected as the control (GFP), where AAV1-hSyn-Cre was injected in the LPB and AAV9-hSyn-DIO-GFP was injected in both the POA and DMH.

(b) Heatmaps of TeNT expression at different Bregma sites from all experimental mice. $POA^{LPB}$ blocking, n = 9 mice; $DMH^{LPB}$ blocking, n = 10 mice; co-blocking, n = 6 mice. The relative scale for the expression intensity (measured by fluorescence intensity) was shown in the right.

(c-f) Changes of basal $T_{core}$ (c-d), basal energy expenditure (e), and basal physical activity (f) at thermoneutral temperature (29$^{o}$C) after blocking LPB-innervating DMH neurons ($DMH^{LPB}$ blocking) or LPB-innervating POA neurons ($POA^{LPB}$ blocking), as indicated (n = 7 each).

(g-h) Changes of basal $T_{core}$ (g) and basal physical activity (h) at thermoneutral temperature (29$^{o}$C) after blocking LPB-innervating POA and DMH neurons (Co-blocking), as indicated (n = 6 each).

(i) Withdrawal latency and the number of lifts in the cold plate test (4$^{o}$C) after blocking LPB-innervating POA/DMH neurons (n = 10 for GFP and $DMH^{LPB}$ blocking group; n = 9 for $POA^{LPB}$ blocking group).

(j) Changes in preference to 30$^{o}$C versus 35/16/10/6$^{o}$C after blocking LPB-innervating POA/DMH neurons (n = 9 mice each).

All data are shown as mean ± sem. The p-values are calculated based on one-way ANOVA (d-h). *p ≤ 0.05; **p ≤ 0.01; ***p ≤ 0.001; ns, not significant.

### Extended Data Fig. 6 | Phenotypes associated with neural blocking of POA neurons, DMH$^{Vglut2}$ neurons and DMH$^{Vgat}$ neurons.

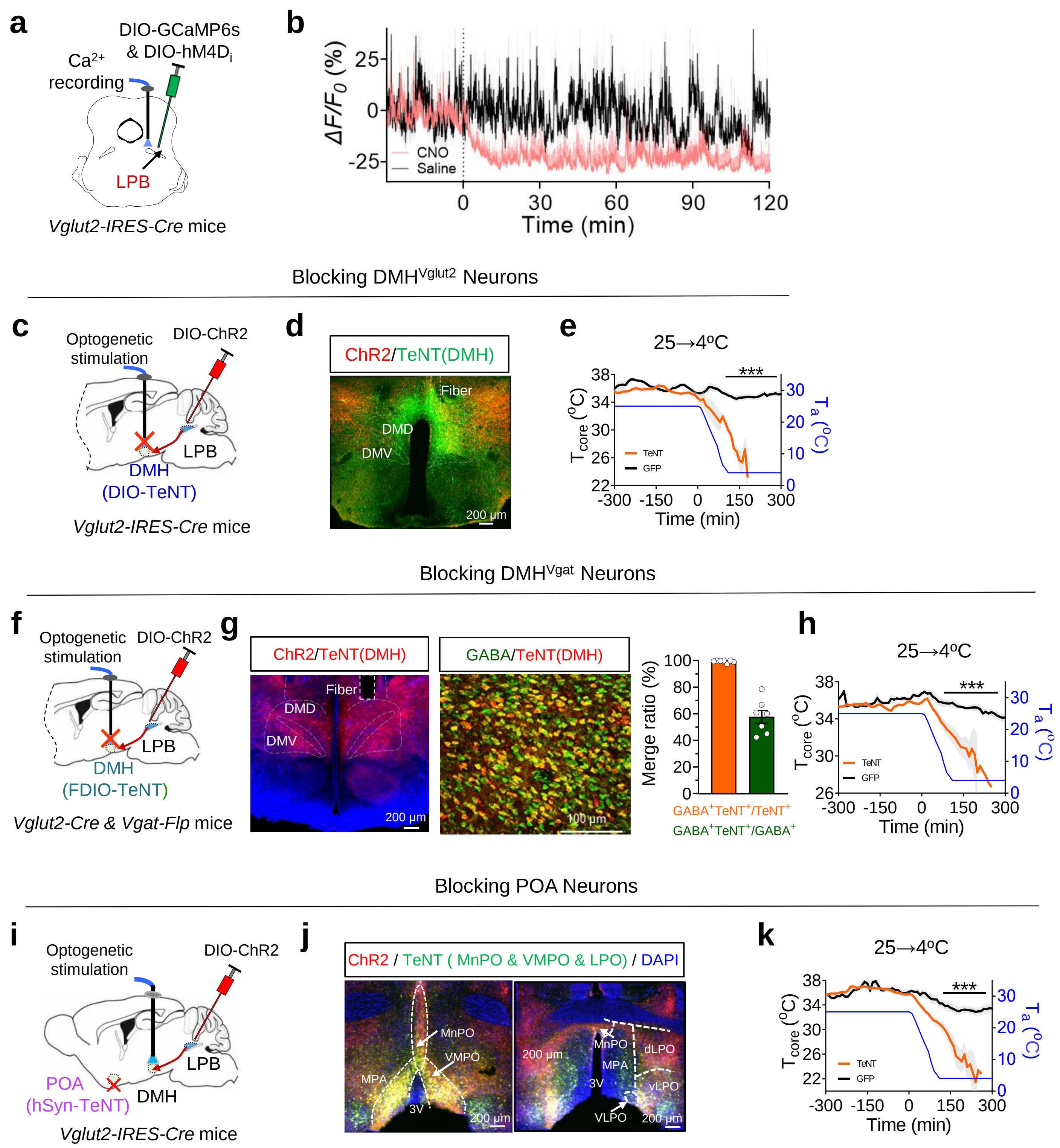

**Extended Data Fig. 6 | Phenotypes associated with neural blocking of POA neurons, $DMH^{Vglut2}$ neurons and $DMH^{Vgat}$ neurons.**

(a) Scheme for calcium photometry of $LPB^{Vglut2}$ neurons before and after chemogenetic inhibition. AAVs carrying inhibitory chemogenetic tool hM4D(Gi) (AAV9-EF1α-DIO-hM4D(Gi)-mCherry) and GCaMP6s (AAV9-hSyn-DIO-GCaMP6s) were co-injected in the LPB of Vglut2-Cre mice to block neural activity and record calcium dynamics of $LPB^{Vglut2}$ neurons, respectively. The optical fiber for calcium photometry was implanted in the LPB.

(b) Calcium dynamics of $LPB^{Vglut2}$ neurons before and after chemogenetic inhibition. CNO was injected (i.p., 10mg/kg) at t = 0 min. $\Delta F/F_0$ represents the change in GCaMP6s fluorescence from the mean level (n = 5 mice).

(c) Scheme for blocking $DMH^{Vglut2}$ neurons using TeNT. AAV5-hSyn-DIO-GFP-P2A-TeNT was injected in the DMH of Vglut2-IRES-Cre mice to block $DMH^{Vglut2}$ neurons.

(d) Representative TeNT expression in the $DMH^{Vglut2}$ neurons.

(e) Changes of $T_{core}$ during cold challenges (25→4$^{o}$C) after blocking $DMH^{Vglut2}$ neurons (TeNT, n = 9 mice; GFP, n = 7 mice).

(f) Scheme for blocking $DMH^{Vgat}$ neurons using TeNT. AAV9-hEF1a-FDIO-mCherry-P2A-TeNT was injected in the DMH of Vglut2-IRES-Cre & Vgat-T2A-FlpO mice to block $DMH^{Vgat}$ neurons.

(g) Representative TeNT expression in $DMH^{Vgat\text{-}FlpO}$ neurons and the colocalization of TeNT and GABA immunostaining. The overlapping ratios were quantified in the right.

(h) Changes of $T_{core}$ during cold challenges (25→4$^{o}$C) after blocking $DMH^{Vgat}$ neurons (TeNT, n = 7 mice; GFP, n = 5 mice).

(i) Scheme for blocking POA neurons using TeNT. AAV9-hSyn-eGFP-2A-TeNT were injected in the POA of Vglut2-IRES-Cre mice to block POA neurons.

(j) Representative TeNT expression in the POA (MnPO, VMPO, MPA and LPO) neurons.

(k) Changes of $T_{core}$ during cold challenges (25→4$^{o}$C) after blocking bulk POA neurons (TeNT, n = 7 mice; GFP, n = 4 mice).

All data are shown as mean ± sem. The p-values are calculated based on two-way ANOVA statistical tests. $^{**}p \leq 0.01$; $^{***}p \leq 0.001$.

### Extended Data Fig. 7 | Properties and whole-brain axonal projections of LPB$^{SST}$ neurons.

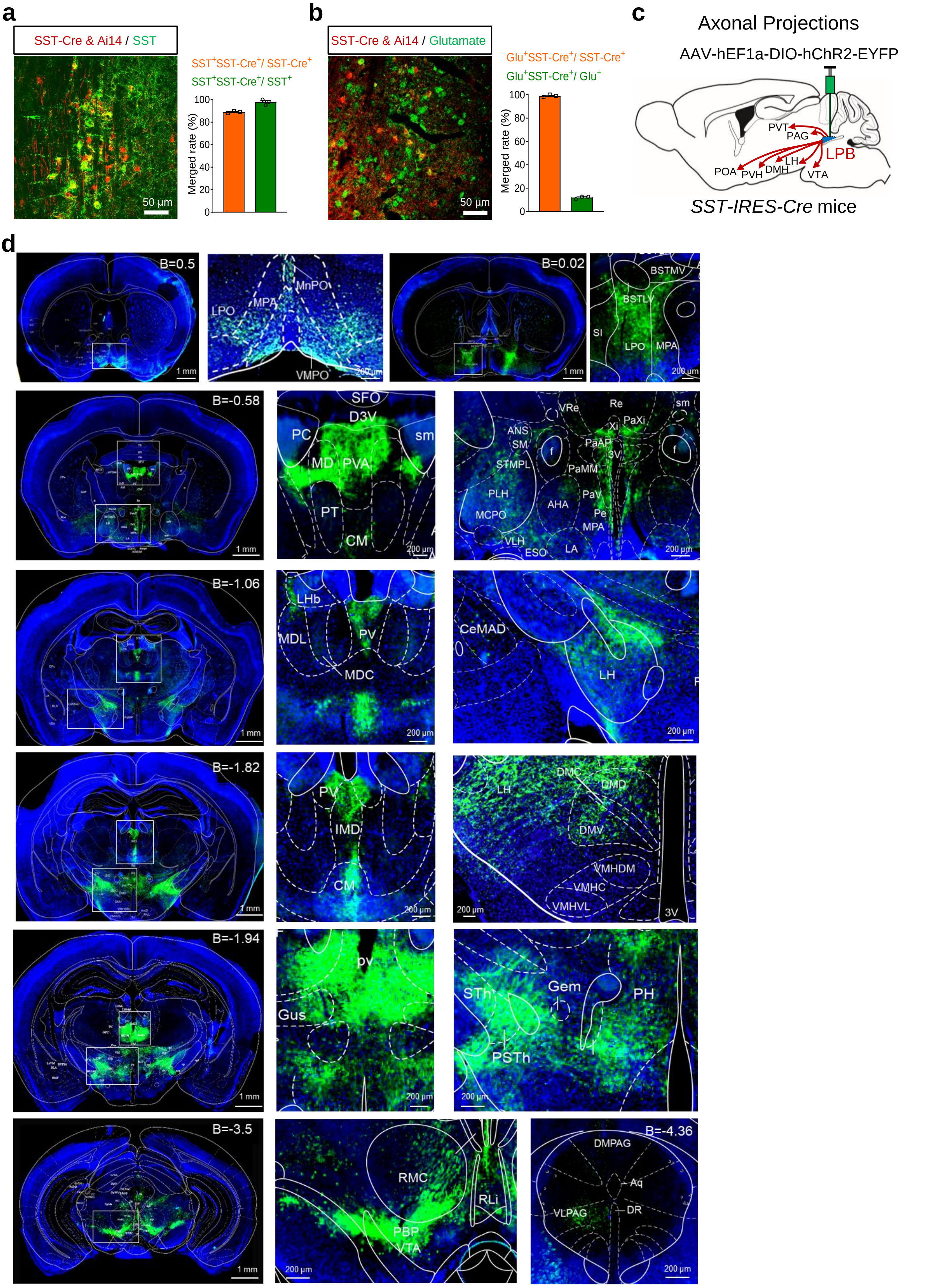

**Extended Data Fig. 7 | Properties and whole-brain axonal projections of LPB$^{SST}$ neurons.**

(a-b) Overlap between SST-IRES-Cre labeled neurons (Tdt$^+$) in the LPB and the following immuno-positive neurons: SST peptide (a) and glutamate (b). SST-IRES-Cre mice were crossed with Ai14 (Rosa-CAG-LSL-tdTomato-WPRE) to label SST-IRES-Cre in the LPB. Overlap ratios are shown on the right of each panel (n = 3 mice each).

(c) The axonal projection pattern of LPB$^{SST}$ neurons throughout the brain, revealed by anterograde tracing AAVs. Injection sites of AAV9-DIO-ChR2-eYFP in the LPB were indicated in the figure. Thicker lines indicated stronger eYFP expressions.

(d) Representative images showing ChR2-eYFP expression in axonal terminals at various brain sites.

AHA, anterior hypothalamic, anterior; B, bregma; BNST, bed nucleus of the stria terminalis; SFO, subfornical organ; sm, stria medullaris of the thalamus; PVA, paraventricular thalamic nucleus, anterior part; PT, paratenia thalamic nucleus; PaAP, paraventricular hypothalamic nucleus, anterior parvic; PaMM, paraventricular hypothalamic nucleus, medial magnocellular part; MDL, mediodorsal thalamic nucleus, lateral part; LHb, lateral habenular nucleus; Pe, periventricular hypothalamic nucleus; PaV, paraventricular hypothalamic nucleus, ventral part; PV, paraventricular thalamic nucleus; BLA, basolateral amygdaloid nucleus, anterior part; CeMAD central amygdaloid nucleus, medial division, anterodorsal part; MD, intermediodorsal thalamic nucleus; DM, central medial thalamic nucleus; PH, posterior hypothalamic nucleus; VMH, ventromedial hypothalamic nucleus; VMHC, ventromedial hypothalamic nucleus, central part; VMHDM, ventromedial hypothalamic nucleus, dorsomedial part; VMHVL, ventromedial hypothalamic nucleus, ventrolateral part; SC, superior colliculus; PSTh parasubthalamic nucleus; STh subthalamic nucleus; POA, preoptic nucleus; Gem, gemini hypothalamic nucleus; Gus gustatory thalamic; VPM, ventral posteromedial thalamic; DMC, dorsomedial hypothalamic nucleus, compact part; DMD, dorsomedial hypothalamic nucleus, dorsal part; RMC, red nucleus, magnocellular part; ZID, zona incerta, dorsal part; Aq, aqueduct(Sylvius); DLPAG, dorsolateral periaqueductal gray; DMPAG, dorsomed periaqueductal; lnG, intermediate gray layer of the superior colliculus; LPO, lateral preoptic area; MPA, medial preoptic area; SIB, substantia innominata, basal part; HDB, nucleus of the horizontal limb of the diagonal band; D3V, dorsal 3rd ventricle; PVT, paraventricular thalamic nucleus.

**Extended Data Fig. 8 | Phenotypes associated with activation of LPB$^{SST}$ neurons.**

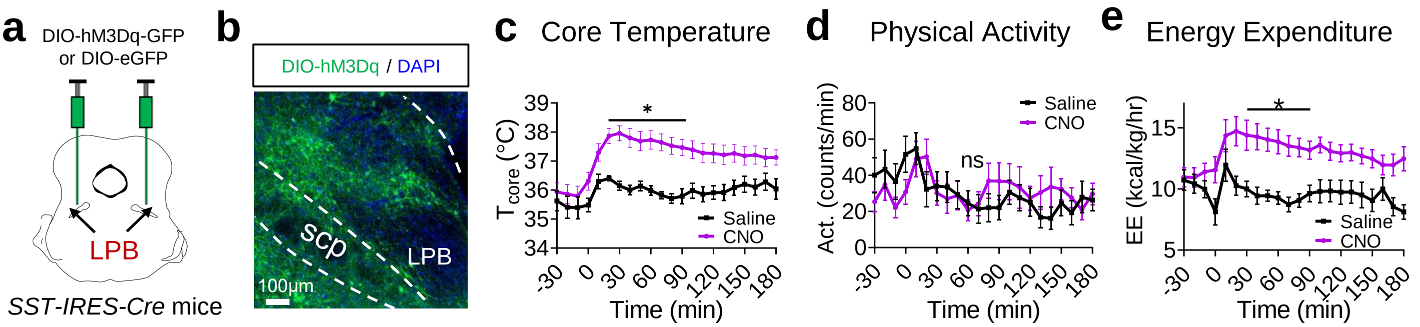

(a) Scheme for chemogenetic activation of LPB$^{SST}$ neurons using hM3D$_q$. AAVs carrying hM3D$_q$ (AAV9-hSyn-DIO-hM3D(G$_q$)-eGFP) were injected in the LPB to activate LPB$^{SST}$ neurons.

(b) Representative expression of hM3D$_q$ in LPB$^{SST}$ neurons.

(c-e) Changes of T$_{core}$ (c), physical activity (d), and energy expenditure (e) after chemogenetic activation of LPB$^{SST}$ neurons (n = 12 mice each). CNO was injected (i.p., 2.5 mg/kg) at t = 0 min.

All data are shown as mean ± sem. The p-values are calculated based on RM two-way ANOVA. *$p \leq 0.05$; ns, not significant.

**Extended Data Fig. 9 | The DMH→RPa pathway is downstream of the LPB→DMH pathway in regulating cold defense.**

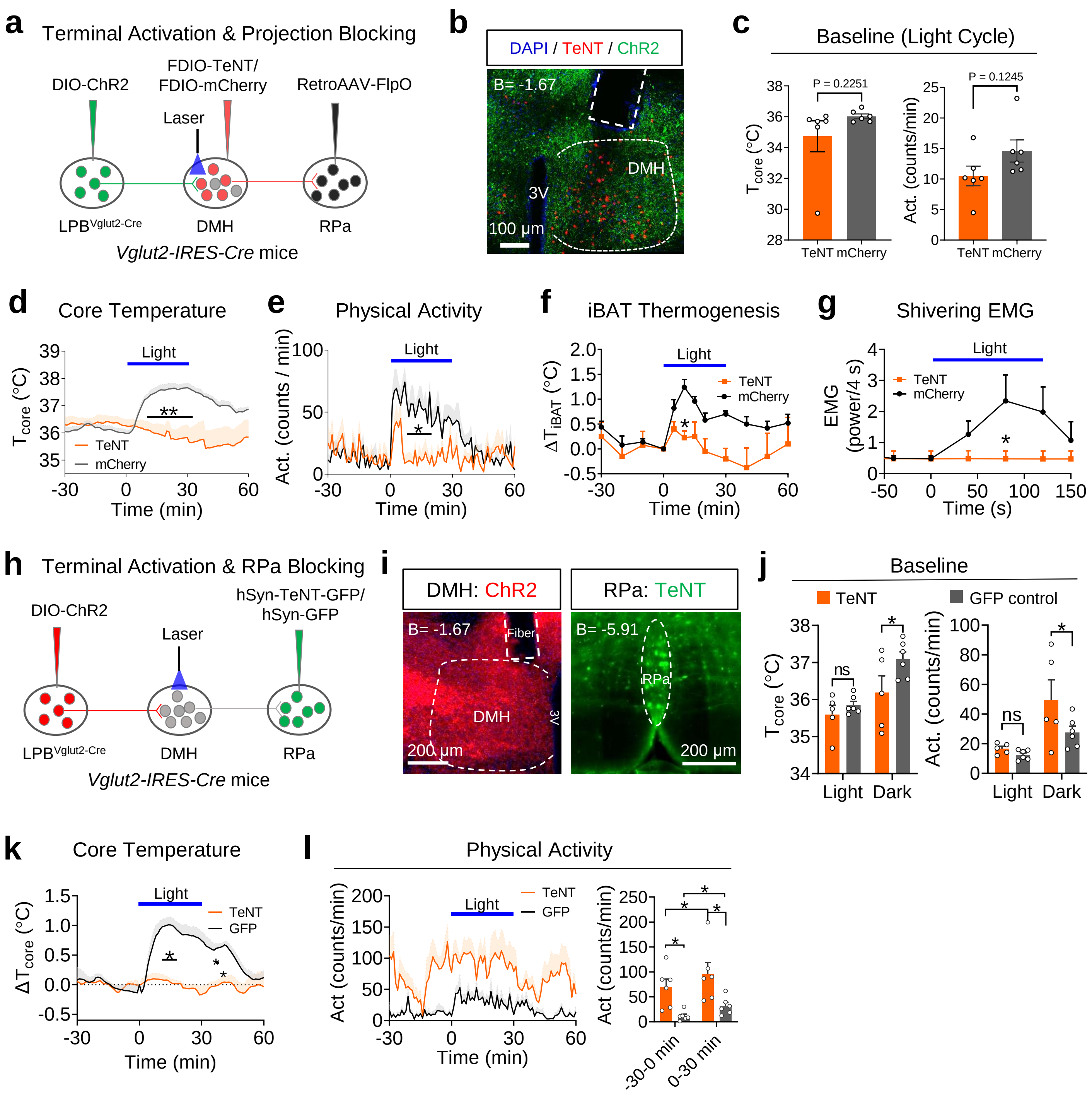

**Extended Data Fig. 9 | The DMH→RPa pathway is downstream of the LPB→DMH pathway in regulating cold defense.**

(a) Photoactivation of $LPB^{Vglut2}$ terminals in the DMH while blocking RPa-projecting DMH neurons. To block RPa-projecting DMH neurons, retrograde AAVs carrying FlpO (AAV-Retro-hSyn-FlpO) were injected in the RPa, which drove the expression of FlpO-dependent TeNT (AAV9-hEF1a-FDIO-mCherry-2A-TeNT) in the DMH.

(b) Representative images of $LPB^{Vglut2\ \&\ ChR2}$ terminals in the DMH (green) and TeNT expression from RPa-projecting DMH neurons (red).

(c) Average basal $T_{core}$ and physical activity after blocking RPa-projecting DMH neurons during the light cycle at $T_a = 26\ ^oC$ (n = 6 mice each).

(d-g) Changes in $T_{core}$ (d), physical activity (e), $T_{iBAT}$ (f), and nuchal muscle EMG (g) in response to photoactivation of $LPB^{Vglut2}$ terminals in the DMH while blocking of RPa-projecting DMH neurons (d-e: TeNT, n = 6 mice; mCherry, n = 7 mice; f-g: TeNT, n = 4 mice; mCherry, n = 5 mice). Light pattern: 473 nm, 6 mW, 10 Hz, 10 ms, 2-s on followed by 2-s off, with the cycles repeating for 30 min in (d-f) and 120 s in (g).

(h) Photoactivation of $LPB^{Vglut2}$ terminals in the DMH while blocking RPa neurons.

(i) Representative images of $LPB^{Vglut2\ \&\ ChR2}$ terminals (red) in the DMH (left panel) and TeNT expression (green) in RPa neurons (right panel). Scale bar, 200 μm.

(j) Average basal $T_{core}$ and physical activity after blocking RPa neurons during the dark and light cycle at $T_a = 26\ ^oC$ (TeNT, n = 5 mice; GFP, n = 6 mice).

(k-l) Changes in $\Delta T_{core}$ (k) and physical activity (l) in response to photoactivation of $LPB^{Vglut2}$ terminals in the DMH while blocking of RPa neurons (n = 6 for each group). Light pattern: 473 nm, 6 mW, 10 Hz, 10 ms, 2-s on followed by 2-s off, with the cycles repeating for 30 min. Note, RPa blocking caused compulsive-like circling behavior at the basal level.

LPB, lateral parabrachial nucleus; POA, preoptic area; DMH, dorsomedial hypothalamic nucleus. RPa, raphe pallidus nucleus. All data are the mean ± SEM, (c) was analyzed by unpaired t-test, and (c-e, j-i) were analyzed by two-way RM ANOVA. ns, not significant; $^*p \le 0.05$ ; $^{**}p \le 0.01$.

### Extended Data Table 2. Summary of statistical analyses

| Figure | Sample size (n) | Statistical test | P values |
|---|---|---|---|
| 1i | Mice: 3<br>Brain slices: 20 | Two-tailed unpaired t-test | t=5.056, df=38<br>P<0.0001 |
| 2e | Warming: 14 mice<br>Cooling: 14 mice | Two-tailed paired t test | t=3.398, df=13, P=0.0048 |
| 2h | GCaMP6: 10 mice<br>21ºC: 30 trials<br>16ºC: 34 trials<br>10ºC: 32 trials<br>4ºC: 33 trials | Brown-Forsythe and Welch ANOVA tests<br>Don’t correct for multiple comparisons test | F* (DFn, DFd) 0.4612 (3.000, 122.7), P=0.7099<br>Multiple comparisons:<br>21ºC vs. 16ºC,P=0.4077<br>21ºC vs. 10ºC, P=0.3677<br>21ºC vs. 4ºC, P=0.2756<br>16ºC vs. 10ºC, P=0.9587<br>16ºC vs. 4ºC, P=0.7521<br>10ºC vs. 4ºC, P=0.7838 |
| 2i | GCaMP6: 10 mice<br>21ºC: 30 trials<br>16ºC: 34 trials<br>10ºC: 32 trials<br>4ºC: 33 trials | Brown-Forsythe and Welch ANOVA tests<br>Don’t correct for multiple comparisons test | F* (DFn, DFd) 17.70 (3.000, 116.8), P<0.0001<br>Multiple comparisons:<br>21ºC vs. 16ºC,P=0.2288<br>21ºC vs. 10ºC, P<0.0001<br>21ºC vs. 4ºC, P<0.0001<br>16ºC vs. 10ºC, P=0.0021<br>16ºC vs. 4ºC, P<0.0001<br>10ºC vs. 4ºC, P=0.0547 |
| 2k | GCaMP6: 10 mice<br>35-20ºC: 20 trials<br>32-17ºC: 17 trials<br>25-10ºC: 26 trials<br>29-14ºC: 17 trials | Brown-Forsythe and Welch ANOVA tests<br>Don’t correct for multiple comparisons test | F* (DFn, DFd) 0.1478 (3.000, 66.04), P=0.9308<br>Multiple comparisons:<br>35-20ºC vs. 32-17ºC, P=0.6186<br>35-20ºC vs. 25-10ºC, P=0.4975<br>35-20ºC vs. 29-14ºC, P=0.9098<br>32-17ºC vs. 25-10ºC, P=0.8965<br>32-17ºC vs. 29-14ºC, P=0.7182<br>25-10ºC vs. 29-14ºC, P=0.6058 |
| 2l | GCaMP6: 10 mice<br>35-20ºC: 20 trials<br>32-17ºC: 17 trials<br>25-10ºC: 26 trials<br>29-14ºC: 17 trials | Brown-Forsythe and Welch ANOVA tests<br>Don’t correct for multiple comparisons test | F* (DFn, DFd) 25.48 (3.000, 103.9), P=0.1478<br>Multiple comparisons:<br>35-20ºC vs. 32-17ºC, P=0.0484<br>35-20ºC vs. 25-10ºC, P<0.0001<br>35-20ºC vs. 29-14ºC, P<0.0001<br>32-17ºC vs. 25-10ºC, P<0.0001<br>32-17ºC vs. 29-14ºC, P=0.0084<br>25-10ºC vs. 29-14ºC, P=0.0033 |
| 2n | GCaMP6: 10 mice<br>0.1ºC/s:19 trials<br>0.2ºC/s: 20 trials<br>0.4ºC/s: 20 trials<br>0.7ºC/s: 20 trials | Brown-Forsythe and Welch ANOVA tests<br>Don’t correct for multiple comparisons test | F* (DFn, DFd) 1.693 (3.000, 57.55), P=0.1785<br>Multiple comparisons:<br>0.1ºC/s vs. 0.2ºC/s, P=0.0356<br>0.1ºC/s vs. 0.4ºC/s, P=0.0696<br>0.1ºC/s vs. 0.7ºC/s, P=0.0853<br>0.2ºC/s vs. 0.4ºC/s, P=0.7986<br>0.2ºC/s vs. 0.7ºC/s, P=0.4623<br>0.4ºC/s vs. 0.7ºC/s, P=0.6562 |
| 2o | GCaMP6: 10 mice<br>0.1ºC/s:19 trials<br>0.2ºC/s: 20 trials<br>0.4ºC/s: 20 trials<br>0.7ºC/s: 20 trials | Brown-Forsythe and Welch ANOVA tests<br>Don’t correct for multiple comparisons test | F* (DFn, DFd) 0.4130 (3.000, 71.93), P=0.7442<br>Multiple comparisons:<br>0.1ºC/s vs. 0.2ºC/s, P=0.8573<br>0.1ºC/s vs. 0.4ºC/s, P=0.3253<br>0.1ºC/s vs. 0.7ºC/s, P=0.6648<br>0.2ºC/s vs. 0.4ºC/s, P=0.4287<br>0.2ºC/s vs. 0.7ºC/s, P=0.8226<br>0.4ºC/s vs. 0.7ºC/s, P=0.5019 |
| 2s | Warming: 6 mice<br>Cooling: 6 mice | Two-tailed paired t test | t=5.560, df=5, P=0.0026 |
| 2t | Warming: 6 mice<br>Cooling: 6 mice | Two-tailed paired t test | t=4.319, df=5, P=0.0076 |
| 3c | GFP: 10 mice<br>DMH$^{only}$ projecting LPB<br>TeNT: 9 mice | RM two-way ANOVA<br>factor one: virus (GFP, DMH$^{only}$ projecting LPB TeNT)<br>factor two: time<br>Uncorrected Fisher’s LSD test | Virus: F (1, 17) = 2.664, P=0.1210<br>Time: F (2.199, 37.38) = 5.516,P=0.0065<br>Interaction: F (360, 6120) = 2.338, P<0.0001<br>Multiple comparisons:<br>GFP vs. DMH$^{only}$ projecting LPB TeNT:<br>95 min – 108 min, P<0.0493<br>117 min – 120 min, P<0.0488<br>123 min – 130 min, P<0.0495 |
| | DMH$^{only}$ projecting LPB<br>TeNT: 9 mice<br>POA$^{only}$ projecting LPB<br>TeNT: 7mice | RM two-way ANOVA<br>factor one: virus ( DMH$^{only}$ projecting LPB TeNT, POA$^{only}$ projecting LPB TeNT)<br>factor two: time<br>Uncorrected Fisher’s LSD test | Virus: F (1, 14) = 0.09332, P=0.7645<br>Time: F (2.201, 30.82) = 8.153, P=0.0011<br>Interaction: F (360, 5040) = 0.7286, P>0.9999<br>Multiple comparisons:<br>POA$^{only}$ projecting LPB TeNT vs. DMH$^{only}$ projecting LPB TeNT:<br>P>0.1575 |
| 3d | GFP: 10 mice<br>DMH$^{only}$ projecting LPB<br>TeNT: 8 mice<br>POA$^{only}$ projecting LPB<br>TeNT: 7mice | RM two-way ANOVA<br>factor one: virus (GFP, DMH$^{only}$ projecting LPB TeNT, POA$^{only}$ projecting LPB TeNT)<br>factor two: time<br>Uncorrected Fisher’s LSD test | Virus: F (2, 22) = 2.757, P=0.0854<br>Time: F (2.837, 62.40) = 14.82, P<0.0001<br>Interaction: F (720, 7920) = 1.745, P<0.0001<br>Multiple comparisons:<br>GFP vs. DMH$^{only}$ projecting LPB TeNT:<br>15 min – 74 min, P<0.0466<br>220 min – 300 min, P<0.0193 |

(*table continued on the next page*)

### Extended Data Table 2. Summary of statistical analyses

| Figure | Sample size (n) | Statistical test | P values |
|---|---|---|---|
| 3e | GFP: 10 mice<br>DMH$^{only}$ projecting LPB TeNT: 8 mice | RM two-way ANOVA<br>factor one: virus (GFP, DMH$^{only}$ projecting LPB TeNT)<br>factor two: time<br>Uncorrected Fisher's LSD test | Virus: $F (1, 16) = 10.66$, $P=0.0049$<br>Time: $F (360, 5760) = 28.00$, $P<0.0001$<br>Interaction: $F (360, 5760) = 5.486$, $P<0.0001$<br>Multiple comparisons:<br>GFP vs. DMH$^{only}$ projecting LPB TeNT:<br>19 min – 69 min, $P<0.0498$<br>159 min – 300 min, $P<0.0098$ |
| | DMH$^{only}$ projecting LPB TeNT: 8 mice<br>POA$^{only}$ projecting LPB TeNT: 7mice | RM two-way ANOVA<br>factor one: virus (DMH$^{only}$ projecting LPB TeNT, POA$^{only}$ projecting LPB TeNT)<br>factor two: time<br>Uncorrected Fisher's LSD test | Virus: $F (1, 13) = 0.03077$, $P=0.8635$<br>Time: $F (360, 4680) = 36.17$, $P<0.0001$<br>Interaction: $F (360, 4680) = 0.4357$, $P>0.9999$<br>Multiple comparisons:<br>DMH$^{only}$ projecting LPB TeNT vs. POA$^{only}$ projecting LPB TeNT: $P>0.1381$ |
| 3f | GFP: 10 mice<br>DMH$^{only}$ projecting LPB TeNT: 8 mice | RM two-way ANOVA<br>factor one: virus (GFP, DMH$^{only}$ projecting LPB TeNT)<br>factor two: time<br>Uncorrected Fisher's LSD test | Virus: $F (1, 16) = 8.921$, $P=0.0087$<br>Time: $F (360, 5760) = 32.63$, $P<0.0001$<br>Interaction: $F (360, 5760) = 5.171$, $P<0.0001$<br>Multiple comparisons:<br>GFP vs. DMH$^{only}$ projecting LPB TeNT: 57 min – 300 min, $P<0.0088$ |
| | DMH$^{only}$ projecting LPB TeNT: 8 mice<br>POA$^{only}$ projecting LPB TeNT: 7mice | RM two-way ANOVA<br>factor one: virus (DMH$^{only}$ projecting LPB TeNT, POA$^{only}$ projecting LPB TeNT)<br>factor two: time<br>Uncorrected Fisher's LSD test | Virus: $F (1, 13) = 2.382$, $P=0.1467$<br>Time: $F (360, 4680) = 47.96$, $P<0.0001$<br>Interaction: $F (360, 4680) = 1.279$, $P=0.0004$<br>Multiple comparisons:<br>DMH$^{only}$ projecting LPB TeNT vs. POA$^{only}$ projecting LPB TeNT:<br>16 min – 62 min, $P<0.0483$ |
| 3h | GFP: 10<br>DMH$^{LPB}$ blocking: 10 | RM two-way ANOVA<br>factor one: virus (GFP, DMH$^{LPB}$ blocking)<br>factor two: time<br>Uncorrected Fisher's LSD test | Virus: $F (1, 18) = 29.88$, $P<0.0001$<br>Time: $F (4.518, 81.32)$ = 15.14, $P<0.0001$<br>Interaction: $F (360, 4680) = 6.670$, $P<0.0001$<br>Multiple comparisons: 37 min – 300 min, $P<0.0075$ |
| | POA$^{LPB}$ blocking: 9<br>Co-blocking: 6 | RM two-way ANOVA<br>factor one: virus (POA$^{LPB}$ blocking, Co-blocking)<br>factor two: time<br>Uncorrected Fisher's LSD test | Virus: $F (1, 13) = 3.875$, $P=0.0707$<br>Time: $F (2.126, 27.64) = 24.71$, $P<0.0001$<br>Interaction: $F (360, 4680) = 1.089$, $P=0.1264$<br>Multiple comparisons: 75 min – 149 min, $P<0.0487$ |
| | POA$^{LPB}$ blocking: 9<br>DMH$^{LPB}$ blocking: 10 | RM two-way ANOVA<br>factor one: virus (POA$^{LPB}$ blocking, DMH$^{LPB}$ blocking)<br>factor two: time<br>Uncorrected Fisher's LSD test | Virus: $F (1, 17) = 0.9149$, $P=0.3522$<br>Time: $F (2.349, 39.93) = 27.73$, $P<0.0001$<br>Interaction: $F (360, 6120) = 1.761$, $P<0.0001$<br>Multiple comparisons: 35 min – 54 min, $P<0.0453$ |
| 3i | GFP: 10<br>DMH$^{LPB}$ blocking: 10 | RM two-way ANOVA<br>factor one: virus (GFP, DMH$^{LPB}$ blocking)<br>factor two: time<br>Uncorrected Fisher's LSD test | Virus: $F (1, 18) = 22.52$, $P=0.0002$<br>Time: $F (3.450, 62.11) = 37.24$, $P<0.0001$<br>Interaction: $F (360, 6480) = 11.66$, $P<0.0001$<br>Multiple comparisons: 7 min – 300 min, $P<0.0095$ |
| | POA$^{LPB}$ blocking: 9<br>Co-blocking: 6 | RM two-way ANOVA<br>factor one: virus (POA$^{LPB}$ blocking, Co-blocking)<br>factor two: time<br>Uncorrected Fisher's LSD test | Virus: $F (1, 13) = 4.561$, $P=0.0523$<br>Time: $F (2.316, 30.10) = 53.89$, $P<0.0001$<br>Interaction: $F (360, 4680) = 6.098$, $P<0.0001$<br>Multiple comparisons: 116 min – 300 min, $P<0.0495$ |
| | POA$^{LPB}$ blocking: 9<br>DMH$^{LPB}$ blocking: 10 | RM two-way ANOVA<br>factor one: virus POA$^{LPB}$ blocking, DMH$^{LPB}$ blocking)<br>factor two: time<br>Uncorrected Fisher's LSD test | Virus: $F (1, 17) = 0.06235$, $P=0.8058$<br>Time: $F (2.683, 45.62) = 39.88$, $P<0.0001$<br>Interaction: $F (360, 6120) = 0.6782$, $P>0.9999$<br>Multiple comparisons: 19 min – 35 min, $P<0.0471$ |
| 3j | GFP: 10<br>DMH$^{LPB}$ blocking: 10 | RM two-way ANOVA<br>factor one: virus (GFP, DMH$^{LPB}$ blocking)<br>factor two: time<br>Uncorrected Fisher's LSD test | Virus: $F (1, 18) = 34.35$, $P<0.0001$<br>Time: $F (2.993, 53.87) = 106.5$, $P<0.0001$<br>Interaction: $F (360, 6480) = 25.91$, $P<0.0001$<br>Multiple comparisons: 31 min – 300 min, $P<0.0008$ |
| | POA$^{LPB}$ blocking: 9<br>DMH$^{LPB}$ blocking: 10 | RM two-way ANOVA<br>factor one: virus (POA$^{LPB}$ blocking, DMH$^{LPB}$ blocking)<br>factor two: time<br>Uncorrected Fisher's LSD test | Virus: $F (1, 17) = 0.2659$, $P=0.6128$<br>Time: $F (2.421, 41.16) = 122.5$, $P<0.0001$<br>Interaction: $F (360, 6120) = 0.6928$, $P>0.9999$<br>Multiple comparisons: $P>0.2914$ |
| | POA$^{LPB}$ blocking: 9<br>Co-blocking: 6 | RM two-way ANOVA<br>factor one: virus (POA$^{LPB}$ blocking, Co-blocking)<br>factor two: time<br>Uncorrected Fisher's LSD test | Virus: $F (1, 13) = 2.316$, $P=0.1520$<br>Time: $F (1.727, 22.45) = 137.5$, $P<0.0001$<br>Interaction: $F (154, 2002) = 3.619$, $P<0.0001$<br>Multiple comparisons: 61 min – 94 min, $P<0.0482$ |
| 3k | GFP: 10<br>DMH$^{LPB}$ blocking: 10 | RM two-way ANOVA<br>factor one: virus (GFP, DMH$^{LPB}$ blocking)<br>factor two: time<br>Uncorrected Fisher's LSD test | Virus: $F (1, 18) = 93.14$, $P<0.0001$<br>Time: $F (2.270, 40.86) = 158.3$, $P<0.0001$<br>Interaction: $F (277, 4986) = 46.53$, $P<0.0001$<br>Multiple comparisons: 31 min – 217 min, $P<0.0009$ |
| | POA$^{LPB}$ blocking: 9<br>DMH$^{LPB}$ blocking: 10 | RM two-way ANOVA<br>factor one: virus (POA$^{LPB}$ blocking, DMH$^{LPB}$ blocking)<br>factor two: time<br>Uncorrected Fisher's LSD test | Virus: $F (1, 17) = 0.02254$, $P=0.8824$<br>Time: $F (277, 4709) = 192.6$, $P<0.0001$<br>Interaction: $F (277, 4709) = 1.468$, $P<0.0001$<br>Multiple comparisons: 22 min – 37 min, $P<0.0429$ |
| | POA$^{LPB}$ blocking: 9<br>Co-blocking: 6 | RM two-way ANOVA<br>factor one: virus (POA$^{LPB}$ blocking, Co-blocking)<br>factor two: time<br>Uncorrected Fisher's LSD test | Virus: $F (1, 13) = 5.089$, $P=0.0419$<br>Time: $F (1.814, 23.58) = 166.2$, $P<0.0001$<br>Interaction: $F (120, 1560) = 4.740$, $P<0.0001$<br>Multiple comparisons: 36 min – 59 min, $P<0.0417$ |

(*table continued on the next page*)

### Extended Data Table 2. Summary of statistical analyses

| Figure | Sample size (n) | Statistical test | P values |
|---|---|---|---|
| 4c | ChR2: 10 mice<br>GFP: 8 mice | RM two-way ANOVA<br>factor one: virus (ChR2, GFP)<br>factor two: time<br>Bonferroni's multiple comparisons test | Virus: $F(1, 16) = 11.26$, $P=0.0040$<br>Time: $F(90, 1440) = 7.309$, $P<0.0001$<br>Interaction: $F(90, 1440) = 12.99$, $P<0.0001$<br>Multiple comparisons: 8 min – 40 min, $P<0.0001$ |
| 4d | ChR2: 10 mice<br>GFP: 8 mice | RM two-way ANOVA<br>factor one: virus (ChR2, GFP)<br>factor two: time<br>Uncorrected Fisher's LSD test | Virus: $F(1, 16) = 30.92$, $P<0.0001$<br>Time: $F(90, 1440) = 4.328$, $P<0.0001$<br>Interaction: $F(90, 1440) = 1.737$, $P<0.0001$<br>multiple comparisons: Before-After: 0 min – 38 min, $P<0.0396$ |
| 4k | Non-blocking: 10 mice<br>$DMH^{Vglut2}$ blocking: 5 mice<br>$DMH^{Vgat}$ blocking: 5 mice<br>POA blocked: 5 mice | RM two-way ANOVA<br>factor one: treatment (Non-blocking, $DMH^{Vglut2}$ blocking, $DMH^{Vgat}$ blocking, POA blocking)<br>factor two: time<br>Bonferroni's multiple comparisons test | Treatment: $F(3, 21) = 9.102$, $P=0.0005$<br>Time: $F(90, 1890) = 11.60$, $P<0.0001$<br>Interaction: $F(270, 1890) = 10.46$, $P<0.0001$<br>Multiple comparisons:<br>Non-blocking vs. $DMH^{Vglut2}$ blocking: 9 min – 45 min, $P<0.0001$<br>Non-blocking vs. $DMH^{Vgat}$ blocking: 7 min – 43 min, $P<0.0001$<br>Non-blocking vs. POA blocking: $P>0.4970$ |
| 4l | Non-blocking: 10 mice<br>$DMH^{Vglut2}$ blocking: 5 mice<br>$DMH^{Vgat}$ blocking: 5 mice<br>GFP:5 mice | RM two-way ANOVA<br>factor one: treatment (Non-blocking, $DMH^{Vglut2}$ blocking, $DMH^{Vgat}$ blocking, GFP)<br>factor two: time<br>Bonferroni's multiple comparisons test | Treatment: $F(3, 21) = 9.566$, $P=0.0003$<br>Time: $F(1, 21) = 13.59$, $P=0.0014$<br>Interaction: $F(3, 21) = 7.305$, $P=0.0015$<br>Multiple comparisons:<br>b.s. vs. 10 min – 30 min:<br>Non-blocking: $P=0.0005$<br>GFP: $P>0.9999$<br>$DMH^{Vglut2}$ blocking: $P=0.8948$<br>$DMH^{Vgat}$ blocking: $P>0.9999$ |
| 4m | 5 mice each | RM two-way ANOVA<br>factor one: treatment (Non-blocking, $DMH^{Vglut2}$ blocking, $DMH^{Vgat}$ blocking, GFP)<br>factor two: time<br>Bonferroni's multiple comparisons test | Treatment: $F(3, 16) = 23.36$, $P<0.0001$<br>Time: $F(9, 144) = 1.652$, $P=0.1060$<br>Interaction: $F(27, 144) = 8.832$, $P<0.0001$<br>Multiple comparisons:<br>GFP vs. Non-blocking: 8 min – 60 min, $P<0.0002$<br>GFP vs. $DMH^{Vgat}$ blocking: 8 min – 30 min, $P<0.016$<br>GFP vs. $DMH^{Vglut2}$ blocking: $P>0.3106$ |
| 4n | 4 mice each | RM two-way ANOVA<br>factor one: treatment (Non-blocking, $DMH^{Vglut2}$ blocking, $DMH^{Vgat}$ blocking)<br>factor two: time<br>Bonferroni's multiple comparisons test | Treatment: $F(3, 12) = 26.85$, $P<0.0001$<br>Time: $F(12, 144) = 9.665$, $P<0.0001$<br>Interaction: $F(36, 144) = 9.762$, $P<0.0001$<br>Multiple comparisons:<br>Non-blocking vs. $DMH^{Vglut2}$ blocking: 40 s – 60 s, $P<0.0001$<br>Non-blocking vs. $DMH^{Vgat}$ blocking: 40 s – 60 s, $P<0.0001$<br>Non-blocking vs. GFP: 40 – 60 s, $P<0.0001$ |
| 4o | 5 mice each | RM two-way ANOVA<br>factor one: treatment (Non-blocking, $DMH^{Vglut2}$ blocking, $DMH^{Vgat}$ blocking,GFP)<br>factor two: time<br>Bonferroni's multiple comparisons test | Treatment: $F(3, 16) = 3.472$, $P=0.0410$<br>Time: $F(90, 1440) = 1.897$, $P<0.0001$<br>Interaction: $F(270, 1440) = 2.218$, $P<0.0001$<br>Multiple comparisons:<br>Non-blocking vs. GFP: 1 – 14 min, $P<0.01$<br>Non-blocking vs. $DMH^{Vglut2}$ blocking: 1 min – 16 min, $P<0.01$<br>Non-blocking vs. $DMH^{Vgat}$ blocking: 1 min – 27 min, $P<0.01$ |
| 4q | ChR2: 6mice | RM one-way ANOVA<br>factor one: treatment (ChR2: 24$^{o}$C, ChR2: 6$^{o}$C, ChR2: 30$^{o}$C, )<br>Bonferroni's multiple comparisons test | Treatment: $F(2, 15) = 1.890$, $P=0.1854$<br>Multiple comparisons:<br>30 $^{o}$C vs. 24 $^{o}$C: $P=0.7829$<br>24 $^{o}$C vs. 6 $^{o}$C: $P=0.7706$ |
| 4r | ChR2: 6mice | RM one-way ANOVA<br>factor one: treatment (ChR2: 24$^{o}$C, ChR2: 6$^{o}$C, ChR2: 30$^{o}$C, )<br>Bonferroni's multiple comparisons test | Treatment: $F(2, 10) = 14.57$, $P=0.0011$<br>Multiple comparisons:<br>30 $^{o}$C vs. 6 $^{o}$C: $P=0.0012$<br>24 $^{o}$C vs. 6 $^{o}$C: $P=0.0027$ |
| 5c | GFP: 9 mice<br>ChR2: 9 mice | Two-tailed unpaired t test | t=3.205, df=16, $P=0.0055$ |
| 5f | ChR2: 9 mice<br>GFP: 5 mice | RM two-way ANOVA<br>factor one: virus (ChR2, GFP)<br>factor two: time<br>Uncorrected Fisher's LSD test | Virus: $F(1, 12) = 1.604$, $P=0.2294$<br>Time: $F(33, 396) = 25.36$, $P<0.0001$<br>Interaction: $F(33, 396) = 1.439$, $P=0.0590$<br>Multiple comparisons: 10 -16 day, $P<0.0261$ |
| 5g | ChR2: 9 mice<br>GFP: 5 mice | RM two-way ANOVA<br>factor one: treatment (GFP, ChR2)<br>factor two: time<br>Uncorrected Fisher's LSD test | Treatment: $F(1, 12) = 3.167$, $P=0.1004$<br>Time: $F(13, 156) = 490.1$, $P<0.0001$<br>Interaction: $F(13, 156) = 2.177$, $P=0.0128$<br>Multiple comparisons:<br>$P>0.0948$ |
| 5h | GFP: 6 mice<br>ChR2: 6 mice | Two-tailed unpaired t test | t=3.348, df=10, $P=0.0074$ |
| 5i | GFP: 6 mice<br>ChR2: 6 mice | Two-tailed unpaired t test | t=2.235, df=10, $P=0.0494$ |
| 5j | ChR2: 6 mice<br>GFP: 6 mice | Ordinary two-way ANOVA<br>factor one: virus (ChR2, GFP)<br>factor two: tissues (fat, lean)<br>Bonferroni's multiple comparisons test | Virus: $F(1, 20) = 5.225$, $P=0.0333$<br>Tissues: $F(1, 20) = 130.0$, $P<0.0001$<br>Interaction: $F(1, 20) = 13.16$, $P=0.0017$<br>Multiple comparisons:<br>Fat: ChR2 vs. GFP: $P=0.0009$<br>Lean: ChR2 vs. GFP: $P=0.583$ |
| 5k | ChR2: 6 mice<br>GFP: 6 mice | Ordinary two-way ANOVA<br>factor one: virus (ChR2, GFP)<br>factor two: time (-200-0, 0-400)<br>Bonferroni's multiple comparisons test | Virus: $F(1, 30) = 1.384$, $P=0.2486$<br>Time: $F(1, 30) = 249.4$, $P<0.0001$<br>Interaction: $F(1, 30) = 1.384$, $P=0.2486$<br>Multiple comparisons:<br>-200 min - 0 min: $P=0.7904$<br>0 min - 400 min: $P=0.0341$ |

(*table continued on the next page*)

### Extended Data Table 2. Summary of statistical analyses

| Figure | Sample size (n) | Statistical test | P values |
|---|---|---|---|
| 7c | ChR2: 8 mice<br>GFP: 7 mice | RM two-way ANOVA<br>factor one: virus (ChR2, GFP)<br>factor two: time<br>Bonferroni's multiple comparisons test | Virus: *F (1, 13) = 8.669*, *P=0.0114*<br>Time: *F (90, 1170) = 10.24*, *P<0.0001*<br>Interaction: *F (90, 1170) = 8.463*, *P<0.0001*<br>Multiple comparisons:<br>14 min – 49 min: *P<0.0061* |
| 7d | ChR2: 20 mice<br>GFP:9 mice | RM two-way ANOVA<br>factor one: virus (ChR2, GFP)<br>factor two: time<br>Bonferroni's multiple comparisons test | Virus: *F (1, 27) = 0.0095*, *P=0.9228*<br>Time: *F (90, 2430) = 1.638*, *P=0.0125*<br>Interaction: *F (90, 2430) = 1.373*, *P=0.0002*<br>Multiple comparisons: *P>0.9999* |
| 7e | ChR2: 6 mice<br>GFP: 7 mice | RM two-way ANOVA<br>factor one: virus (ChR2, GFP)<br>factor two: time<br>Bonferroni's multiple comparisons test | Virus: *F (1, 11) = 31.63*, *P=0.0002*<br>Time: *F (18, 198) = 7.932*, *P<0.0001*<br>Interaction: *F (18, 198) = 7.532*, *P<0.0001*<br>Multiple comparisons:<br>5 min – 40 min: *P<0.0037* |
| 7f | Sham: 6 mice<br>Denervation: 6 mice | RM two-way ANOVA<br>factor one: treatment (Sham, Denervation)<br>factor two: time<br>Bonferroni's multiple comparisons test | Treatment: *F (1, 10) = 3.611*, *P=0.0866*<br>Time: *F (89, 890) = 12.18*, *P<0.0001*<br>Interaction: *F (89, 890) = 5.664*, *P<0.0001*<br>Multiple comparisons:<br>22 min – 34 min: *P<0.0390* |
| 7g | SST: 4 mice<br>Vglut2: 4 mice | RM two-way ANOVA<br>factor one: mice (SST, Vglut2)<br>factor two: time<br>Bonferroni's multiple comparisons test | Mice: *F (1, 6) = 259.6*, *P<0.0001*<br>Time: *F (12, 72) = 9.856*, *P<0.0001*<br>Interaction: *F (12, 72) = 8.922*, *P<0.0001*<br>Multiple comparisons:<br>40 s – 60 s, *P<0.0001* |
| 7h | SST: 8 mice<br>Vglut2: 5 mice | RM two-way ANOVA<br>factor one: mice (SST, Vglut2)<br>factor two: time<br>Bonferroni's multiple comparisons test | Mice: *F (2, 15) = 1.564*, *P=0.2416*<br>Time: *F (89, 1335) = 1.907*, *P<0.0001*<br>Interaction: *F (178, 1335) = 2.628*, *P<0.0001*<br>Multiple comparisons:<br>0 min – 16 min, *P<0.0012* |
| 7k | ChR2: 6 mice<br>ChR2 + TeNT:6 mice | RM two-way ANOVA<br>factor one: virus (ChR2, ChR2 + TeNT)<br>factor two: time<br>Bonferroni's multiple comparisons test | Virus: *F (1, 10) = 0.04145*, *P=0.8428*<br>Time: *F (2.483, 24.83) = 0.04145, P<0.0001*<br>Interaction: *F (18, 180) = 0.8825*, *P=6003*<br>Multiple comparisons: *P>0.9878* |
| 7l | ChR2: 6 mice<br>ChR2 + TeNT:6 mice | RM two-way ANOVA<br>factor one: virus (ChR2, ChR2 + TeNT)<br>factor two: time<br>Bonferroni's multiple comparisons test | Virus: *F (1, 10) = 0.4483*, *P=0.5183*<br>Time: *F (4.334, 43.34) = 21.94*, *P<0.0001*<br>Interaction: *F (18, 180) = 3.162*, *P<0.0001*<br>Multiple comparisons: *P>0.9999* |
| 7n | taCasp3: 8 mice<br>mCherry: 8 mice | RM two-way ANOVA<br>factor one: virus (taCasp3, mCherry)<br>factor two: time<br>Bonferroni's multiple comparisons test | Virus: *F (1, 14) = 0.01322*, *P=0.7216*<br>Time: *F (60, 840) = 35.12*, *P<0.0001*<br>Interaction: *F (60, 840) = 0.5366*, *P=0.9985*<br>Multiple comparisons: *P>0.9999* |
| 7o | taCasp3: 8 mice<br>mCherry: 8 mice | RM two-way ANOVA<br>factor one: virus (taCasp3, mCherry)<br>factor two: time<br>Bonferroni's multiple comparisons test | Virus: *F (1, 14) = 12.95*, *P=0.0029*<br>Time: *F (60, 455) = 47.2*, *P<0.0001*<br>Interaction: *F (60, 775) = 12.17*, *P<0.0001*<br>Multiple comparisons: 180 min – 300 min, *P<0.0014* |
| 7r | Non-blocking: 8 mice<br>$DMH^{Lep\&SST\text{-}TeNT}$: 7 mice<br>$DMH^{ChAT\&SST\text{-}TeNT}$: 7 mice | RM two-way ANOVA<br>factor one: treatment (Non-blocking, $DMH^{Lep\&SST\text{-}TeNT}$, $DMH^{ChAT\&SST\text{-}TeNT}$)<br>factor two: time<br>Bonferroni's multiple comparisons test | Virus: *F (1, 13) = 11.18*, *P=0.0053*<br>Time: *F (90, 1170) = 13.96*, *P<0.0001*<br>Interaction: *F (90, 1170) = 4.768*, *P<0.0001*<br>Multiple comparisons:<br>Non-blocking vs. $DMH^{Lep\&SST\text{-}TeNT}$, 11 min – 51 min, *P<0.0381* |
